# The influence of trait intolerance of uncertainty on behavioural flexibility

**DOI:** 10.1101/2024.10.30.621039

**Authors:** Brendan Williams, Claudia Rodriguez-Sobstel, Lily FitzGibbon, Jayne Morriss, Anastasia Christakou

## Abstract

Identifying and responding adaptively to a change in our environment is an essential skill. However, differences in our ability to detect and our disposition to react to these changes mean that some individuals are better equipped to deal with change than others. Here, we investigate whether intolerance of uncertainty, a transdiagnostic dimension of psychopathology, is associated with performance during a reversal learning task. We assessed task performance of 145 subjects using behavioural measures and computational modelling across two time points (approximately 12 days apart). Intolerance of uncertainty and its prospective and inhibitory subscales were associated with task performance, irrespective of self-reported levels of trait anxiety. Intolerance of uncertainty and its inhibitory subscale were positively associated with increased sensitivity to reinforcement from positive, but not negative, feedback. Furthermore, the inhibitory subscale was positively associated with better performance, both overall (as indexed by accuracy) and immediately following a change in reward contingencies (as indexed by perseveration). Lastly, the prospective subscale interacted with the extent to which choice was driven by expected value across time points. These findings provide novel evidence for how trait intolerance of uncertainty may modulate behavioural flexibility in changeable environments. The study points towards exciting avenues for further research into the development of IU-related behaviours across the lifespan and their implications for mental health.

## Introduction

Intolerance of uncertainty (IU) is the tendency to interpret and react to uncertainty negatively (Carleton, 2016; Freeston et al., 1994). Prior research has shown that during uncertain situations individuals with higher IU report greater subjective ratings of threat (Cupid et al., 2021; Pepperdine et al., 2018) and negative affect (Morriss, Goh, et al., 2023), as well as display greater physiological arousal (Morriss et al., 2021; Tanovic, Gee, et al., 2018). IU is commonly captured using the 12-item IU questionnaire, which can be used as a single scale or two subscales (Carleton et al., 2007; Hong & Lee, 2015). The inhibitory IU subscale measures action paralysis in the face of uncertainty and the prospective IU subscale measures the desire for predictability (Carleton et al., 2007). Both the total IU score and the inhibitory/prospective subscales evidence robust psychometrics and are reliably normally distributed across community samples (Carleton et al., 2012; Hong & Lee, 2015). Importantly, a wealth of recent evidence has demonstrated that higher IU is common across a variety of mental health conditions such as anxiety, depression, schizophrenia and eating disorders (Carleton et al., 2012; McEvoy et al., 2019; Morriss & Ellett, under review). Because of this, there has been a surge of interest in characterising how IU maps onto psychological processes, in order to determine how to effectively target IU in mental health treatment and support (Einstein, 2014; Shihata et al., 2016). In the current article, we explore how individual differences in IU relate to behaviour in uncertain and changing environments using a reversal learning task. This builds on previous work investigating the role of IU in threat learning and goal-directed behaviour, reviewed in detail below.

In affective neuroscience research, there have been advancements in understanding the role of IU in psychological processes related to Pavlovian threat learning (e.g. a cue is associated with an outcome) (for review see Lonsdorf & Merz, 2017; Morriss et al 2021). In particular, previous studies have found that individuals higher in IU are sensitive to changes in the value of cues that have been predictive of aversive primary reinforcers (e.g. loud noise or electric shock). For instance, higher IU (and both subscales) is specifically associated with delayed threat extinction learning (e.g. continued responding to cues that no longer signal threat) and greater threat generalisation (e.g. responding parametrically to cues based on their similarity to a learned threat cue). Furthermore, in Pavlovian reversal learning, where the values of threatening and safe cues become inverted, individuals with higher IU demonstrate greater reversal of physiological responses (e.g. skin conductance) (Morriss et al., 2019). However, this IU-related effect in reversal learning differs based on the contingency instructions provided before reversal occurs, such that higher IU is associated with greater reversal of physiological responses when contingencies are uninstructed (e.g. participants must learn the contingencies themselves) (Morriss et al., 2019), but not when instructed (e.g. participants are given information about which cue is associated with threatening or safe outcomes) (Mertens & Morriss, 2021). Overall, this research suggests that IU plays an important role in Pavlovian threat learning processes, particularly those related to the updating of contingency information.

Currently, the literature on the role of IU and operant threat learning (e.g. an action such as a button press is reinforced through an aversive outcome) is still emerging (for review see Krypotos et al., 2015). A recent systematic review identified eleven studies that examined the role of IU in threat avoidance with primarily low-cost stakes (e.g. where failed avoidance results in little or no punishment/loss) (for review see Wong et al. 2023). The review highlighted that higher IU is associated with greater threat-driven generalisation of avoidance behaviours (e.g. avoiding visual cues that look similar to a learned threat cue) and the prospective IU subscale is associated with goal-directed and persistent avoidance. In parallel to this work, there has been a handful of studies that have started to investigate how IU impacts decision-making and actions related to reward (Ciria et al., 2021; Luhmann et al., 2011; Radell et al., 2016, 2018; Tanovic, Hajcak, et al., 2018). For instance, Luhmann et al. (2011) found that individuals with higher IU tended to select immediate smaller rewards, compared to waiting for less probable larger rewards. However, a replication attempt of this study in a larger sample only yielded a similar pattern of results with the inhibitory IU subscale and not the total IU scale (Ciria et al., 2021). In sum, these studies suggest that IU may bias behaviour towards avoidance of uncertain threats and approach towards certain rewards.

Despite this progress, there are gaps in our understanding as to how IU (and its constituents) modulates learning in environments that are changeable (for discussion, see Sandhu et al 2023) and where choices or actions result in different, non-deterministic outcomes (e.g. gain, loss) (see Carleton et al 2016; Kornilova et al 2018). These competences are important in scenarios where behaviour needs to regularly change in order to achieve a set goal. Imagine an individual who often travels between countries and has to change the way they behave in each environment (e.g. cultural norms, local laws). We ask how individual differences in how people cope with uncertainty (specifically IU and its constituents) modulate their ability to detect and adapt to such changing states of the environment.

To address this gap in the literature, we examined IU and behavioural flexibility under changing environments using reversal learning with a mixture of reward and loss outcomes. In reversal learning tasks the contingencies between actions and their outcomes change during the task (Izquierdo et al., 2017). These tasks can be set up such that participants are instructed about the occurrence but not the frequency of the reversals, and about the probabilistic nature of choice outcomes (but not the exact contingencies). This setup enables the investigation of how participants flexibly implement adaptive strategies under uncertainty and refine their internal model of the task, unconfounded by a potentially protracted trial-and-error rule-learning process which is subject to significant individual differences (Bell et al., 2018; Bell, Langdon, et al., 2019; Bell, Lindner, et al., 2019).

In this work we examined the association between IU and its constituent subscales with performance in a serial reversal task, repeated at two timepoints approximately two weeks apart. Participants were presented with potential choices which are probabilistically associated with reward or loss. The contingencies between choices and outcomes change multiple times throughout the task, with choices that were previously rewarding resulting in losses, and vice versa. To perform optimally in this task, participants therefore need to consistently select the choices that are most likely to yield a reward, and responsively adapt their choice strategy when contingencies change. Behavioural measures of task performance include accuracy, reaction time, win-stay/lose-shift behaviour, and perseverative responding (i.e., not changing response strategy after contingencies change). Performance can also be assayed by using computational modelling to fit reinforcement learning models to choice behaviour. These models posit latent contributions to task performance, such as a learning rate, or how quickly the subjective values of different choices are updated based on the experienced outcomes. Computational modelling allows us to quantify the contribution of these parameters to performance, and their association with IU. In this work we assess performance using reinforcement learning models from two families that enable us to arbitrate between contributions of choice stochasticity and reinforcement sensitivity to individual differences in behaviour. We model choice stochasticity in one family by allowing the inverse temperature – which governs the extent to which choice behaviour is driven by expected value – to freely vary; in the other we allow reinforcement sensitivity – which places an upper limit on the maximum difference between expected values – to vary. The effect of these parameters on choice behaviour is functionally the same, but the differing parameters enable the alternative fitting and interpretation of behaviour (Waltmann et al., 2022).

Expanding the hitherto limited literature on the influence of IU on adaptive learning in an environment where both gains and losses are possible, we use previously shared data (Williams et al., 2024) to test whether individual differences in IU and its subscales were associated with behavioural and computational features of reversal learning performance. We tested the stability of these associations across two timepoints, and further assessed the specificity of the IU effect by controlling for self-reported general anxiety using data collected retrospectively at a third timepoint.

## Methods

### Participants

Participants were enrolled in this study using the online recruitment platform Prolific (https://www.prolific.co/). Out of a total of 257 participants recruited to the study, 145 participants completed the study protocol at both Times 1 and 2, and passed all attention checks and performed at better than chance level in the reversal learning task (according to a binomial test) (age *M* = 34.34, *SD* = 13.33; sex = 64.14% female; please refer to Supplementary Materials for additional participant demographics and participant filtering). Of these 145 participants, 118 (age *M* = 37.64, *SD* = 13.36; sex = 68.64% female) also completed study Time 3, meaning that in addition to reversal learning performance, we have complete measures of intolerance of uncertainty (all three timepoints) and trait anxiety (Time 3) as measured using STICSA.

Participants who successfully completed Time 1 of the study were reimbursed £1.75 for their time. This payment was framed as a basic pay rate of £1.25, plus an additional 50p that would be awarded based on their performance in the reversal learning task. This was done to maximise the likelihood that participants would remain focused while completing the task. Participants who completed Time 1 of the study but failed the nonsensical attention check or the C/IE responding checks were reimbursed £1.25 for their time. Participants who failed the instructional attention check, or who did not complete their submission were not reimbursed, per participation rules on Prolific.

Participants who successfully completed Time 2 of the study were reimbursed £2.50 for their time. As in Time 1, this payment was framed as a basic rate of £1.25, plus a performance bonus of £1.25. This payment bonus was larger during Time 2 of the study than the first to encourage participants return for Time 2 of the study. Participants who completed Time 2 of the study but failed the nonsensical attention check or the C/IE responding checks were reimbursed £1.25 for their time. Participants who failed instructional attention checks, or who did not complete their submission for Time 2 were not reimbursed, per participation rules on Prolific.

Having recognised the need to control for generalised anxiety, participants who successfully completed Time 2 of the study were retrospectively invited to take part in a third session (Time 3). In Time 3 participants provided responses to the Intolerance of Uncertainty Scale (IUS-12) and the State-Trait Inventory for Cognitive and Somatic Anxiety (STICSA, described below). Of the 213 participants invited to take part in Time 3, 176 completed the questionnaires. Nine participants failed the attention check questions and were therefore not paid for their time. The remaining participants were reimbursed 50p (equivalent to an average hourly rate of £10.84/hr) for their time. There were approximately two weeks between Time 1 and Time 2 (*M* = 12.27, *SD* = 1.57 days), and 1.5 years between Time 2 and Time 3 (*M* = 522.95, *SD* = 73.95 days).

### Questionnaires

The IUS-12 was completed at Times 1, 2 and 3. STICSA was measured once at Time 3. We also added additional questions to the Intolerance of Uncertainty Scale as attentional checks, see section Careless / Insufficient Effort Responding checks below.

#### Intolerance of Uncertainty Scale 12 (IUS-12)

The IUS-12 is a 12-item self-report measure of emotional, cognitive, and behavioural responses to uncertainty (Carleton et al., 2007). The scale has excellent internal consistency, α = .91 (Carleton et al., 2007) and Cronbach’s alpha was comparable in our sample across timepoints, averaging α = .91 (for details please refer to Supplementary Materials) . For each of the items, participants are asked to rate how characteristic it is of them on a 5-point Likert scale, where 1 = not at all characteristic of me, and 5 = entirely characteristic of me. The IUS-12 has a two-factor subscale, representing 7 prospective items (e.g. *I can’t stand being taken by surprise*) and 5 inhibitory items (e.g. *When it’s time to act, uncertainty paralyses me*), which reflect the desire for predictability and paralysis in the face of uncertainty respectively (Carleton et al., 2007). Both subscales have excellent internal consistency (α = .85 for both, Carleton et al. 2007), with comparable internal consistency in our sample (average α = .87) across time (for details refer to Supplementary Materials). A total score of intolerance of uncertainty is the sum of all items, with a minimum score of 12 and maximum score of 60. Scores for subscales range from 7 to 35 for the prospective subscale and from 5 to 25 for the inhibitory subscale. A high overall score indicates higher intolerance of uncertainty, with higher scores in each subscale indicating higher prospective or inhibitory IU.

To confirm our measures of intolerance of uncertainty were stable over time, we calculated measures of reliability to test whether IU scores are stable over time as posited by the construct of IU, and whether it was appropriate to use mean scores in our analyses. The intraclass correlation coefficients (ICC) of intolerance of uncertainty and its prospective and inhibitory subscales were computed to assess their test-retest reliability across A: Time 1, 2 and 3, and B: Time 1 and 2 only. ICC estimates and their 95% CIs were computed using the *psych* package (Revelle, 2024) for each iteration based on a mean-rating (k = 3 for Times 1, 2 and 3; k = 2 for Times 1 and 2), absolute-agreement, two-way mixed-effects model.

The full ICCs are reported in Table 1 below. The ICCs for Time 1, 2 and 3 ranged from 0.90 to 0.92, and for Time 1 and 2 from 0.90 to 0.92, indicating excellent reliability according to the thresholds suggested by Koo & Li (2016). The corresponding F-tests indicated that each ICC was statistically significant (all *p*s < .001).

**Table 1.**
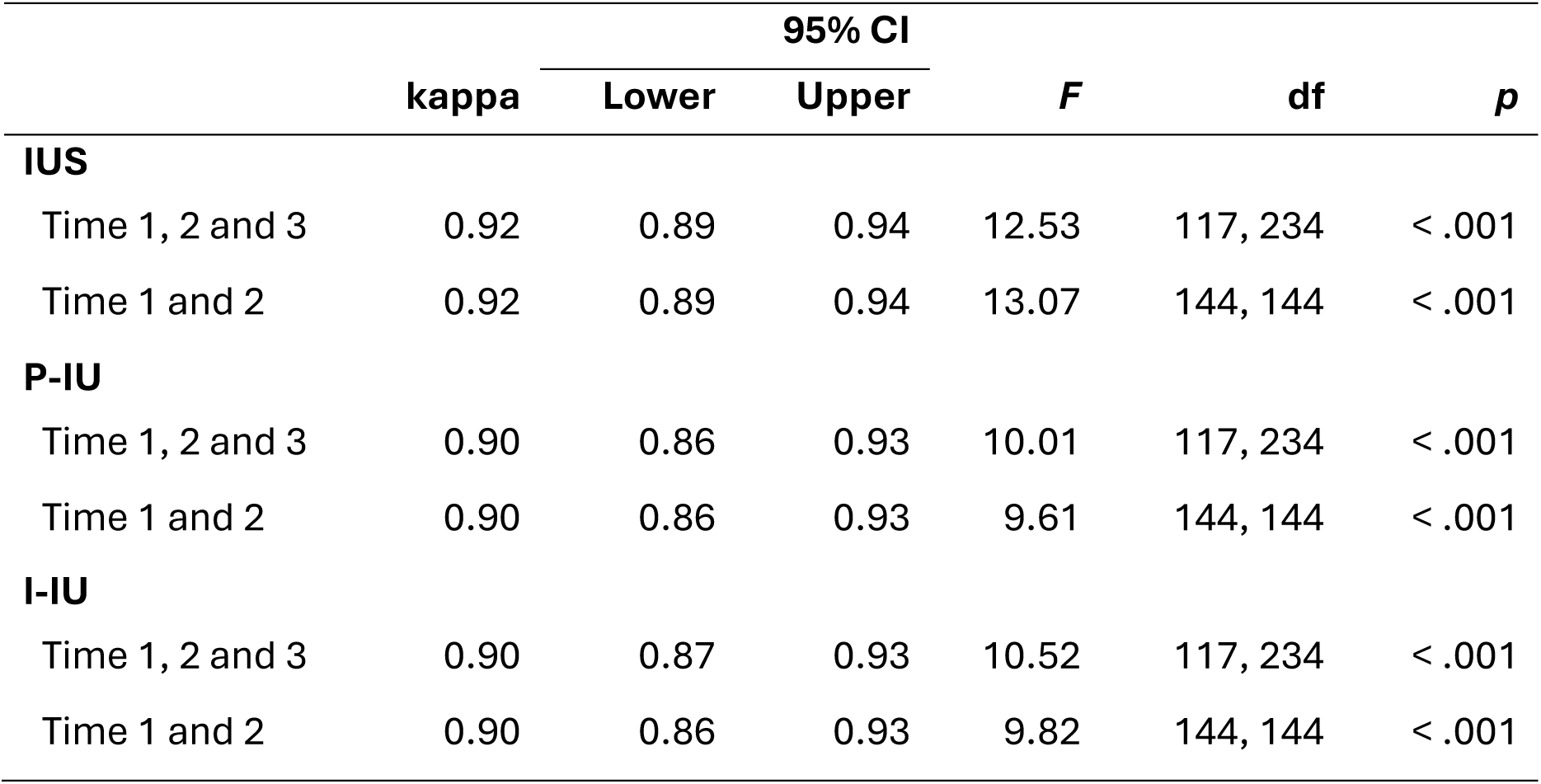
Results of Intraclass Correlation Coefficient Analyses for Intolerance of Uncertainty (IUS) and its Prospective (P-IU) and Inhibitory (I-IU) Subscales.

Given the excellent reliability of intolerance of uncertainty and its subscales across time, we concluded that there had not been major changes in IU across the three measured timepoints. Scores for intolerance of uncertainty, and its prospective and inhibitory subscales, were therefore averaged across Time 1 and Time 2 for participants who had only completed two timepoints (N = 27) and across Time 1, Time 2 and Time 3 who had completed all timepoints (N = 118). This enabled us to maximise our sample size, and use Time as a fixed effect, which in turn enables clearer interpretations in relation to the behavioural and computational modelling variables (measured at Time 1 and Time 2).

#### State-Trait Inventory for Cognitive and Somatic Anxiety (STICSA)

The STICSA (trait only: Ree et al. 2008) is a 21-item self-report measure. The scale has excellent internal consistency, commonly > .87 (Grös et al., 2007), with α = .91 in our sample. For each of the items, participants are asked to read each statement (e.g., *I feel agonised over my problems* or *My face feels hot*) and, using a 4-point Likert scale, indicate how often, in general, the statement is true of them, where 1 = not at all, and 4 = very much so. Total scores range from 21-84, with higher scores indicating higher levels of trait anxiety.

Correlations between STICSA and intolerance of uncertainty/its subscales at each time (as well as averaged across Time) were computed using the *psych* package (Revelle, 2024). Each of the correlations demonstrated a significant positive association (all *p*s < .001). Full results can be found in Supplementary Table 19 and are visualised in Supplementary Figure 11, Supplementary Figure 12 and Supplementary Figure 13.

### Reversal learning task

Participants were presented with two visually distinguishable abstract stimuli that would appear randomly either left of centre, or right of centre on screen. Participants selected one of these stimuli by pressing the corresponding arrow key on their keyboard. Participants were given up to two seconds to make a valid choice response, selecting one of the stimuli presented to them. After stimulus selection participants were presented with the outcome of their choice. Choice outcomes were either the gain or loss of a single point. Initially, one stimulus was randomly assigned as the ‘correct’ stimulus. Selection of the ‘correct’ stimulus meant participants had an 75% chance of gaining a point, and 25% chance of losing a point. The ‘incorrect’ stimulus had the inverse probabilities of the ‘correct’ choice for gaining and losing points. Outcomes for the ‘correct’ and ‘incorrect’ stimuli were pseudorandomised so the assigned outcome probabilities were true over contiguous blocks of five trials. Participants experienced ten ‘learning events’ across the task, which consisted of one initial learning event and nine reversal learning events, with each reversal occurring every 15 ± 3 trials (uniform distribution). At the point of reversal, the identity of the ‘correct’ stimulus was changed, such that the ‘correct’ stimulus became the ‘incorrect’ stimulus and vice versa. If participants did not make a valid choice response within the two second time limit, then they were told they were too slow, and lost a single point. Participants completed 150 trials of the reversal learning task (Figure 1). This task was created using the JavaScript library jsPsych (https://www.jspsych.org/) version 6.1.0, and custom JavaScript code. The relevant JavaScript code for running the task is available in the following GitHub repository: (https://github.com/bwilliams96/SR_Online_task). The reversal learning task was hosted using Gorilla (https://gorilla.sc).

**Figure 1.**
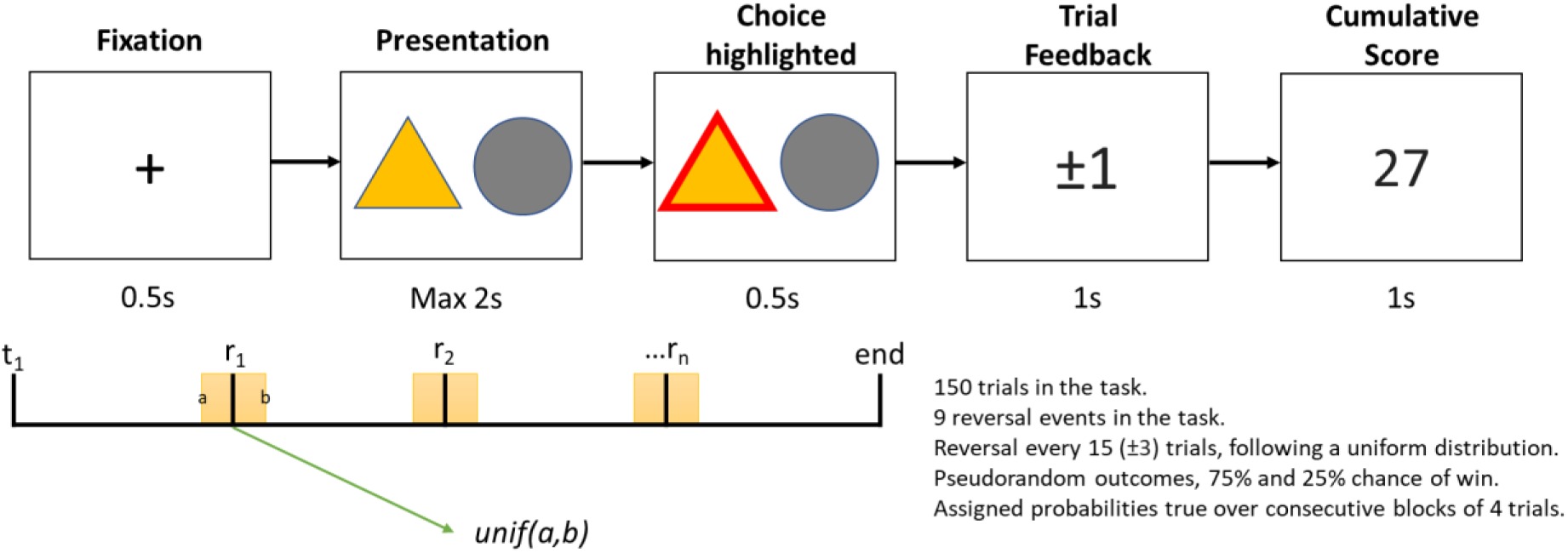
Overview of the trial-level and overall reversal learning task structure

### Checks for Careless / Insufficient Effort Responding

The reversal learning task included automated careless / insufficient effort (C/IE) responding checks that terminated the task prematurely if met. These conditions were 1) not making a valid choice over five consecutive trials or 2) not making a valid choice for over 5% of the total number of trials. We also added additional questions to the Intolerance of Uncertainty Scale to check for C/IE. We added two instructional attention check questions (“I am certain that I have read these questions carefully. Press two as your response.”, and “Worrying about good quality data is something that we as experimenters do. Therefore, we ask that you please select five as your response for this question.”) and one infrequency attention check question (“I have spent lots of time over the last two weeks worrying about the 1977 Olympics” – this question is nonsensical since no Olympics were held in 1977) (Zorowitz et al., 2021). The latter infrequency question was included since this follows best practices for using nonsensical attentional checks in online research (Huang et al., 2015; Zorowitz et al., 2021). We also added attention check questions to the STICSA. We added one instructional attention check question (“I need to respond “Moderately” to this question”), and two infrequency attention check questions (“I am living on earth” and “My skin is made of teeth”). Participants were made explicitly aware of the use of attentional checks, and that their responses to the Intolerance of Uncertainty Scale questions would not influence their bonus payment. The intolerance of uncertainty scale was hosted on the online platform Gorilla (https://gorilla.sc) for Time 1 and Time 2, and both the STICSA and intolerance of uncertainty scale were hosted on REDCap for Time 3.

Participants who passed our built-in C/IE responding checks in the reversal learning task and the intolerance of uncertainty questionnaire had their behavioural data screened again. We identified potential C/IE participants using the following performance metrics: responding using a single key for > 2/3 of all trials; responding to one stimulus for > 2/3 of all trials; reaction time < 250m/s on 10% or more trials; and accuracy (choice of correct option regardless of outcome) significantly worse than chance, based on a binomial test. These criteria are based on previously described C/IE criteria for an online reversal learning task (Zorowitz et al., 2021). Participants who failed the accuracy C/IE test were automatically removed [n = 58], because this suggested they did not learn in the task, and therefore the fitting of a reinforcement learning model to their behaviour would be inappropriate.

### Behavioural measures

Behavioural measures of task performance were derived from choices made in the reversal learning task as described in Williams et al. (2024). Accuracy is the probability of selecting the “best” (most likely to be rewarding) choice on trial n, regardless of whether a reward was obtained or not. We also calculated accuracy slopes to account for changes in accuracy across the task. To do this we calculated the proportion of “best” choices for each learning event, and estimated the intercept and beta parameters for each subject in each session using a mixed effects model. We found a model with linear terms (BIC = 25182.37) provided a better description of the data than a model with quadratic terms (BIC = 25207.61), therefore we report results using coefficients from our linear model. Perseveration is a measure of persistence in choosing the previously “best” choice after reversal, and is defined as the probability of selecting the “worst” (least likely to be rewarding) choice on trial n, after receiving two losses when making that choice following a reversal. Stay/switching behaviour is determined as the probability of making the same choice as on the previous trial, and is defined as the probability of staying both overall, and after either a win or loss on the previous trial.

Reaction time is defined as the amount of time the participant took to make a choice on trial n and, like staying behaviour, is calculated both overall, and whether a win or loss was experienced on the previous trial. We also calculated the difference in reaction time following a win and loss (win RT- loss RT). Behavioural measures were estimated from mixed effects logistic/linear (dependent on whether variable was binary) regression models that calculated estimates for each session using a model which accounted for the effect of session.

### Computational Modelling

The computational modelling of reversal learning behaviour used in this paper reuses results from Williams et al. (2024). Briefly, in Williams et al. (2024) we fit two families of models, differentiated by how expected value determined choice. The first family used an inverse temperature parameter (β) in the softmax choice function to define choice stochasticity by determining the steepness of the softmax function. The second family used a reinforcement sensitivity parameter (ρ) to determine the maximum difference in expected values between choices, placing a lower bound on choice stochasticity. Although the overall best fitting model was a reinforcement sensitivity model, the use of both model families enables us to theorise with more granularity about the component processes of interest. Broadly, the stochasticity models enable us to examine the influence of reinforcement history on exploration, while the reinforcement sensitivity models allow us to examine the influence of the nature of choice outcomes on future choice.

To determine the best fitting model within each family, we fitted a range of models with combinations of parameters that are commonly used in the reinforcement learning literature. Models had either a single inverse temperature (β) / sensitivity parameter (ρ) or separate inverse temperature (*β*_*win*/*loss*_) / sensitivity parameters (*ρ*_*win*/*loss*_) for wins and losses.

Expected values for actions were updated using prediction errors (*λ*_*t*_ − *V_t_*^*k*^), the difference in value of the actual (λ) and expected outcome (*V*) of action *k* on trial *t* (*V_t_*^*k*^). The rate of expected value updating was captured by the learning rate, with models having either a single learning rate (*α*) or separate learning rates for wins and losses (*α*^+/−^, dual learning rate models). Models either updated the expected value of only the chosen action (single update models) or of both the chosen and unchosen actions (dual update models) using the inverse outcome for updating the unchosen action. The dual update models were fit with and without a discount weight (*κ*) for the unchosen action. An in-depth explanation of model variants can be found in Williams et al. (2024).

For model fitting, each model described in the previous section was fitted to the data with one of three estimation methods, maximum likelihood (ML), maximum *a posteriori* estimation with uninformative priors (MAP0), and maximum *a posteriori* estimation with empirical priors inferred from the multivariate distribution of parameter estimates across subjects (Expectation-Maximisation – EM). The best fitting model for each family was determined using the integrated Bayesian information criterion (iBIC) from the EM fitting approach, with a lower iBIC indicating a better model fit (Huys et al., 2011, 2012; Williams et al., 2024). Here, we report associations between computational modelling parameters and IU using parameter estimates from the best-fitting model for the reinforcement sensitivity and softmax model families.

All models are described in detail in (Williams et al. 2024 and in the supplementary material. We briefly describe here the overall best fitting model which was from the reinforcement sensitivity family of models, and was a dual-update model with separate reinforcement sensitivity parameters for wins and losses, and a single learning rate. The choice rule for these models, defined as the probability of making choice *k* on trial *t* is defined as:

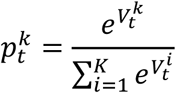

Expected value updates for chosen and unchosen actions were defined using the following formulae:

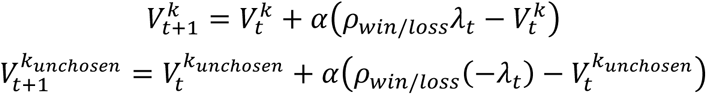

Where expected values *V* are updated based on prediction errors using the actual *λ*_*t*_ and counterfactual −*λ*_*t*_ outcome of trial *t* for chosen and unchosen actions, respectively, and scaled by the learning rate *α*.

For the softmax family of models, the best fitting model was a dual-update model with separate inverse temperatures and learning rates for wins and losses, and a discount weight for the unchosen action. The choice rule for these models, defined as the probability of making choice *k* on trial *t* is defined as:

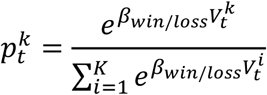

Which includes separate inverse temperature parameters *β* for wins and losses. Expected value updates for chosen and unchosen actions were defined using the following formulae:

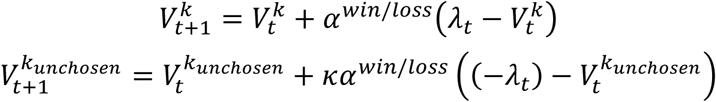

Where expected values *V* are updated based on prediction errors using the actual *λ*_*t*_ and counterfactual −*λ*_*t*_ outcome of trial *t* for chosen and unchosen actions, respectively, and scaled by the learning rate *α* (separate for wins and losses), and the discount weight *κ* for unchosen options.

### Analysis Plan

We conducted multilevel models (MLMs) in R version 4.1.3 using the *lmer* function from the *lme4* package (Bates et al., 2015). We first implemented separate MLMs to establish whether there were any effects observed across time (and outcome type (gain/loss) where relevant) for each dependent variable. The variables that were assessed for time and outcome were (1) reaction time, (2) stay, (3) reinforcement sensitivity from the reinforcement computational model, as well as (4) inverse temperature and (5) learning rate from the softmax computational model. The remaining variables (accuracy, accuracy slopes, perseveration, difference in RTs after wins and losses, learning rate from the reinforcement computational model and discount weight from the softmax computational model) were only assessed for time. For each of these MLMs, Time (Time 1, Time 2) and (where relevant) Outcome (Win, Loss) were entered at level 1 and individual subjects at level 2. Fixed effects included Time and (where relevant) Outcome and random effects included a random intercept for each individual. A maximum likelihood estimator was utilised in all models. Time and Outcome were categorical and therefore effect coded (Time: Time 1 = -1, Time 2 = 1; Outcome: Win = 1, Loss = -1). We then implemented separate MLMs to investigate the effect of intolerance of uncertainty predictors, where IUS/P-IU/I-IU scores were included as mean-centred continuous predictors at level 2 in the MLMs, with all parameters as described above.

In line with previous work (Klingelhöfer-Jens et al., 2022; Rodriguez-Sobstel et al., 2023) where significant effects or interactions with IU variables were observed, follow-up MLMs were implemented with both mean-centred IUS/P-IU/I-IU and STICSA scores included to account for any contributions of trait anxiety as measured using the STICSA and to isolate effects related to IU alone, over and above effects of trait anxiety. For the 27 participants who had not returned for Time 3 and for whom we did not have trait anxiety (STICSA) data, missing values (‘NA’) were entered. Additionally, where significant effects or interactions were observed with one IU subscale and not the other, follow-ups with both mean-centred subscale scores included in the model were conducted to assess their specificity. A significant effect or interaction with one of these predictors but not the other (IUS/P-IU/I-IU or STICSA) would indicate specificity of that predictor. To visualise these effects, plots of simple slopes are presented, alongside scatter plots of the contributing variables. We report only the corrected results in the main results section, but all results are reported in the supplementary materials.

## Results

Intolerance of uncertainty was found to have a significant effect on reinforcement sensitivity, such that a significant interaction between intolerance of uncertainty and outcome was found when entered into the model alone (*p* = .001, see Table 2) as well as when controlling for trait anxiety (IUS x Outcome: *F*(1, 357) = 6.53, *p* = .011; STICSA x Outcome: *F*(1, 357) = 0.28, *p* = .599) (Table 2). From the simple slopes, we observed that higher intolerance of uncertainty was associated with higher reinforcement sensitivity to wins but not for losses (see Figure 2A). Similarly to total intolerance of uncertainty, there was a significant interaction between the inhibitory subscale and Outcome on reinforcement sensitivity both when entered into the model alone (*p* = .001, see Table 2) as well as after controlling for trait anxiety (I-IU x Outcome: *F*(1, 357) = 6.59, *p* = .011; STICSA x Outcome: *F*(1, 357) = 0.21, *p* = .535) and for the prospective IU subscale (I-IU x Outcome: *F*(1, 435) = 11.67, *p* = .001; P-IU x Outcome: *F*(1, 435) = 1.71, *p* = .192). As with the total intolerance of uncertainty score, the simple slopes demonstrated that higher inhibitory IU was associated with higher reinforcement sensitivity to wins but not losses (see Figure 2B). Importantly, these IU-related effects were stable across time, as there were no interactions with the time of testing (I-IU x Time: *F*(1, 435) = 0.76, *p* = .385; I-IU x Time x Outcome: *F*(1, 435) = 1.31, *p* = .254).

**Figure 2.**
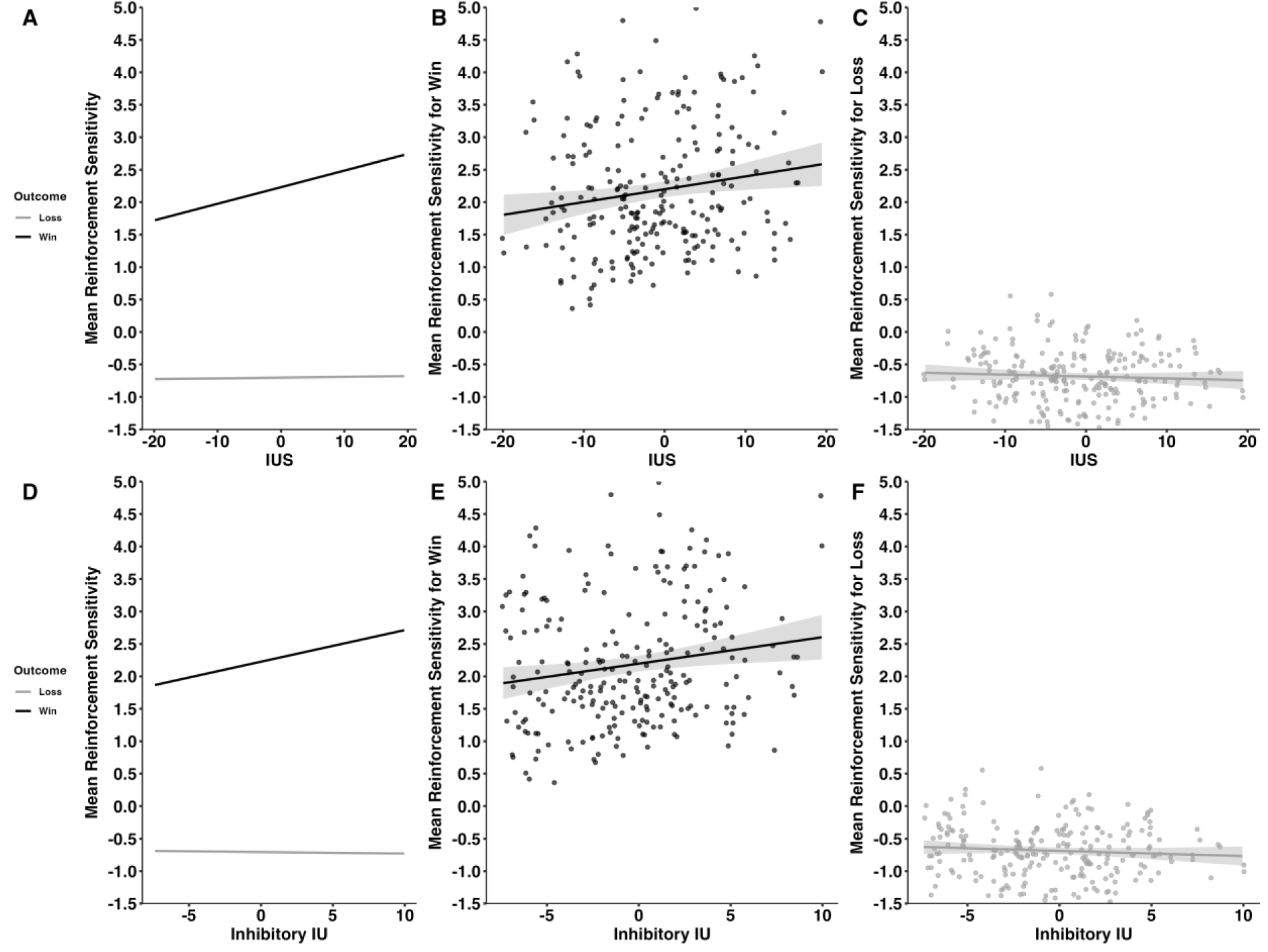
Simple Slopes Depicting Association Between (A) Intolerance of Uncertainty (IUS; mean-centred) and Reinforcement Sensitivity per Outcome and (B) its Corresponding Scatterplot for Wins and (C) for Loss, (D) Inhibitory Intolerance of Uncertainty (I-IU; mean-centred) and Reinforcement Sensitivity per Outcome and (E) its Corresponding Scatterplot for Wins and (F) for Loss. Both higher overall IUS and inhibitory IU were associated with higher reinforcement sensitivity for wins but not losses. Shaded regions represent 95% confidence intervals.

**Table 2.**
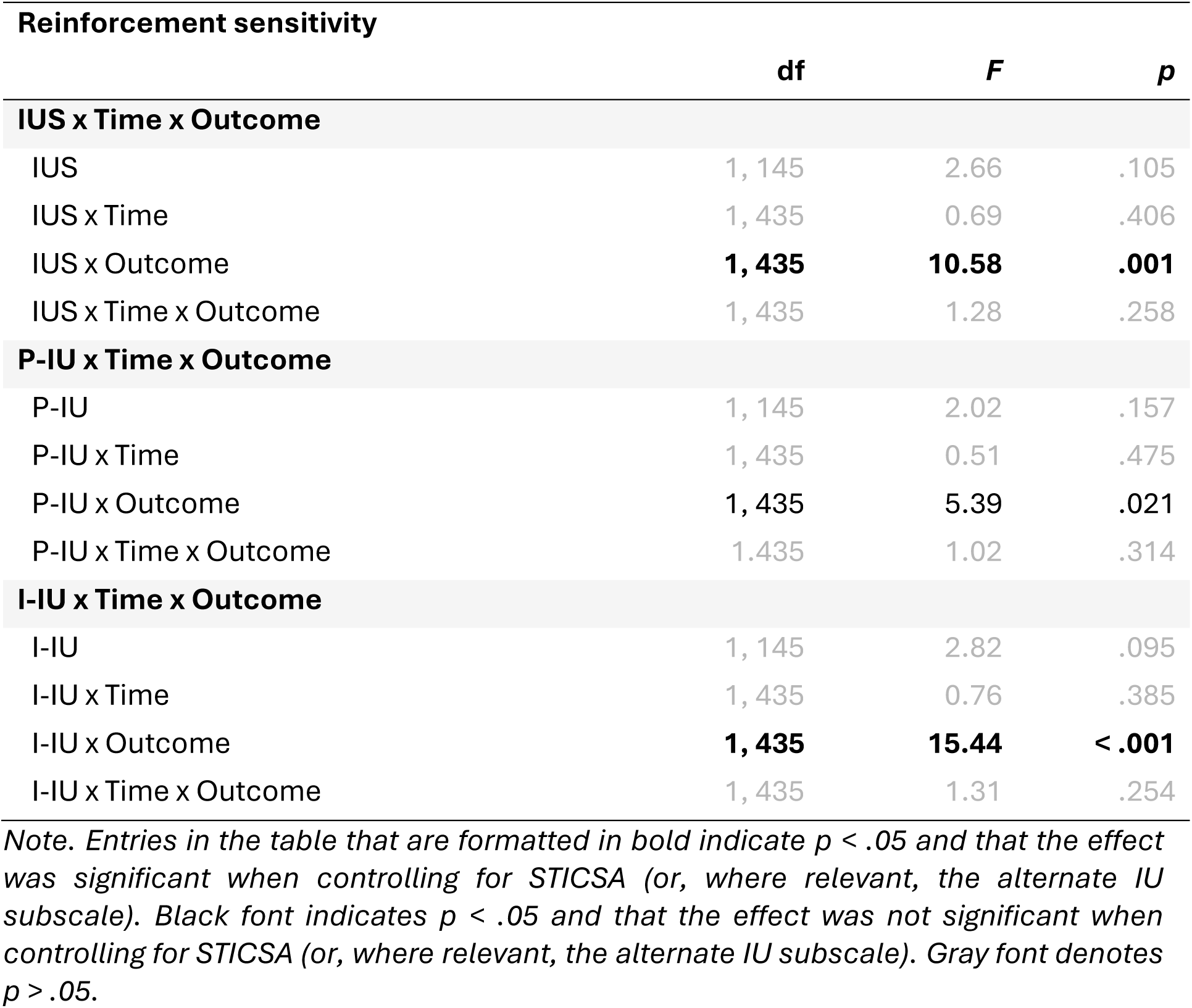
Results from MLMs Assessing Intolerance of Uncertainty, Prospective Intolerance of Uncertainty, and Inhibitory Intolerance of Uncertainty Main Effects and Interactions with Time and Outcome on Reinforcement Sensitivity.

A significant main effect of the inhibitory subscale on accuracy was found when entered into the model alone (*p* = .033, see Table 3) as well as when entered with trait anxiety (main effect of inhibitory subscale: *F*(1, 119) = 5.93, *p* = .016; main effect of trait anxiety: *F*(1, 119) = 0.60, *p* = .441) and the prospective subscale (main effect of inhibitory subscale: *F*(1, 145) = 4.87, *p* = .029; main effect of prospective subscale: *F*(1, 145) = 1.29, *p* = .258). Based on the simple slopes, we observed that a higher inhibitory subscale score was associated with higher accuracy (see Figure 3).

**Figure 3.**
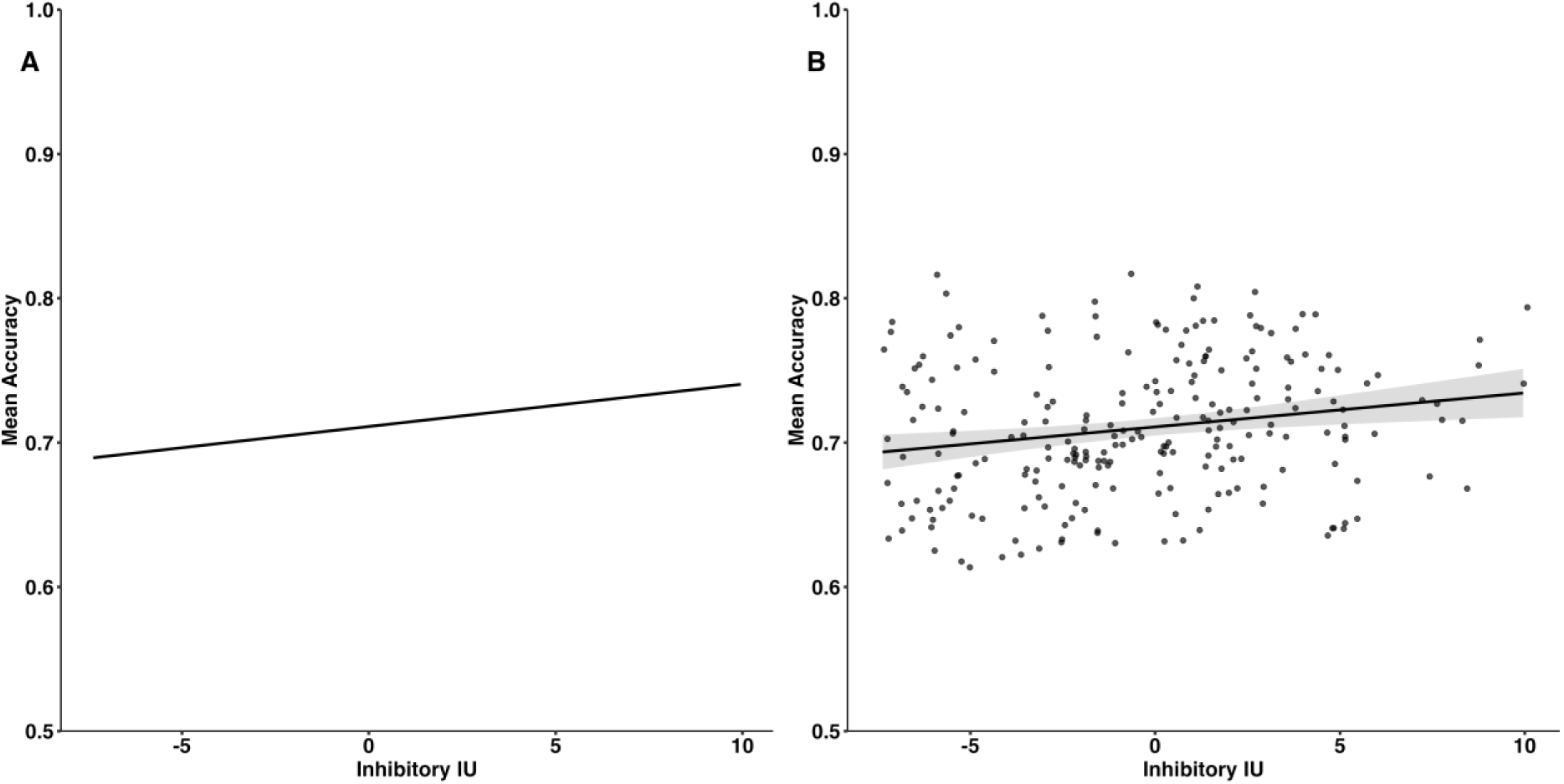
(A) Simple Slopes Depicting Association Between Inhibitory Intolerance of Uncertainty (I-IU; mean-centred) and Accuracy and (B) its Corresponding Scatterplot. Higher I-IU was associated with higher accuracy. Shaded regions represent 95% confidence intervals.

**Table 3.**
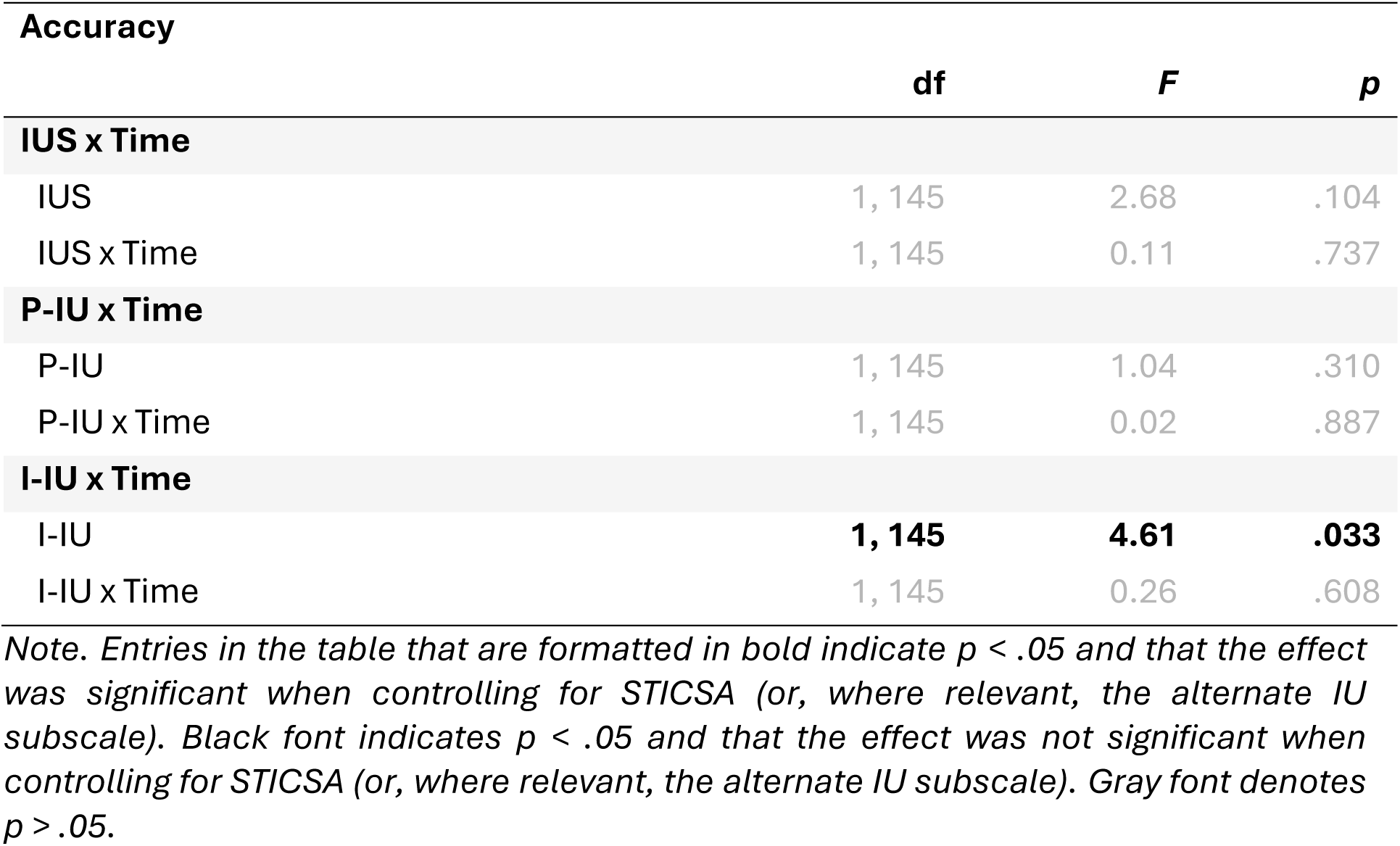
Results from MLMs Assessing Intolerance of Uncertainty, Prospective Intolerance of Uncertainty, and Inhibitory Intolerance of Uncertainty Main Effects and interactions with Time on Accuracy.

The inhibitory subscale of the intolerance of uncertainty scale was also significantly associated with perseverative behaviour during the reversal learning task, both when entered into the model alone (*p* = .035, see Table 4) as well as when entered with trait anxiety (main effect of inhibitory subscale: *F*(1, 119) = 5.19, *p* = .024; main effect of trait anxiety: *F*(1, 119) = 0.51, *p* = .477) and the prospective subscale (main effect of inhibitory subscale: *F*(1, 145) = 5.14, *p* = .025; main effect of prospective subscale: *F*(1, 145) = 1.50, *p* = .222). The simple slopes showed that higher inhibitory subscale scores were associated with fewer perseverative errors (Figure 4). These effects were also stable across time, as there were no interaction effects with time (see Table 4).

**Figure 4.**
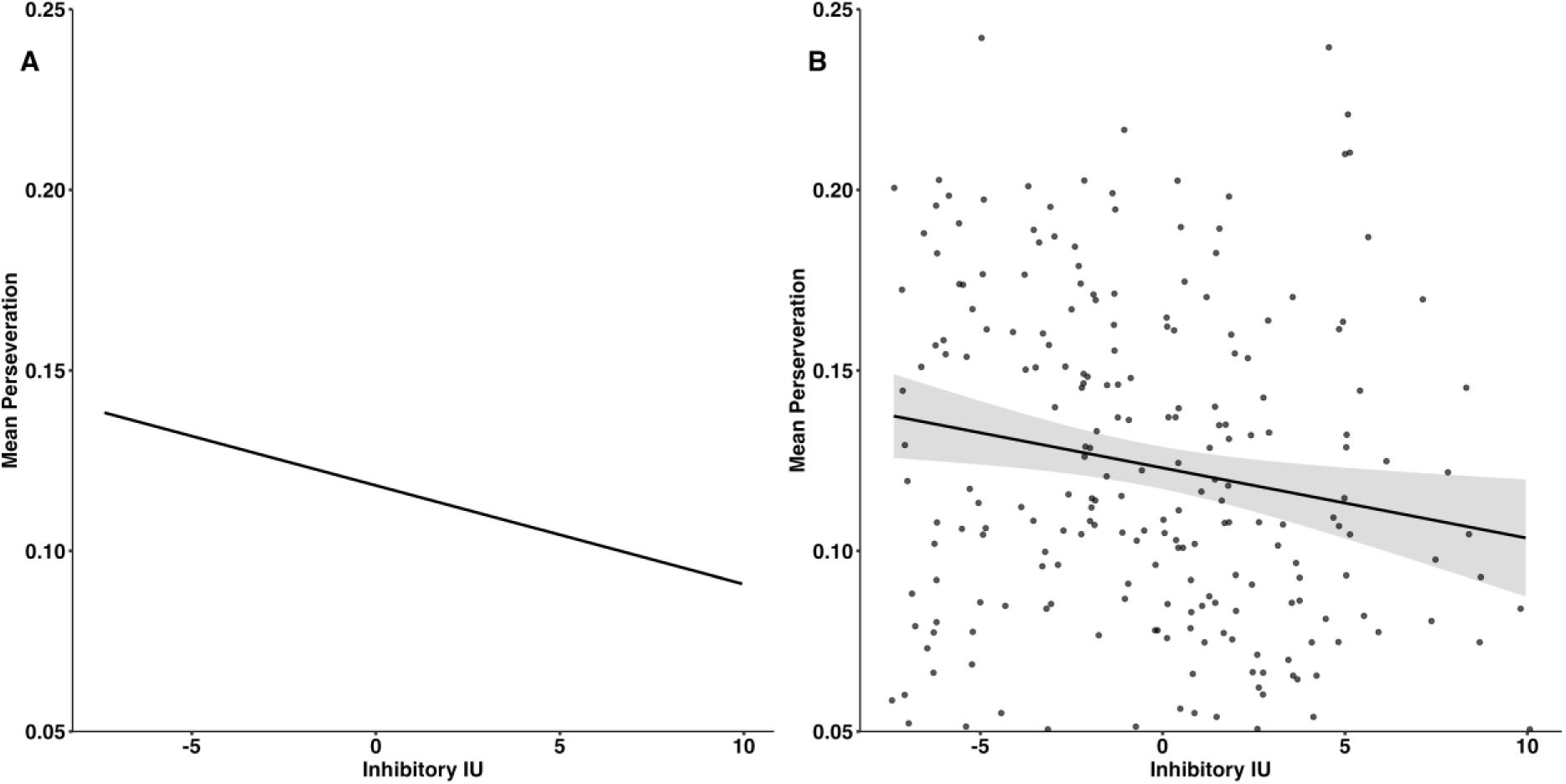
(A) Simple Slopes Depicting Association Between Inhibitory Intolerance of Uncertainty (I-IU; mean-centred) and Perseveration and (B) its Corresponding Scatterplot. Higher I-IU was associated with fewer preservative errors. Shaded regions represent 95% confidence intervals.

**Table 4.**
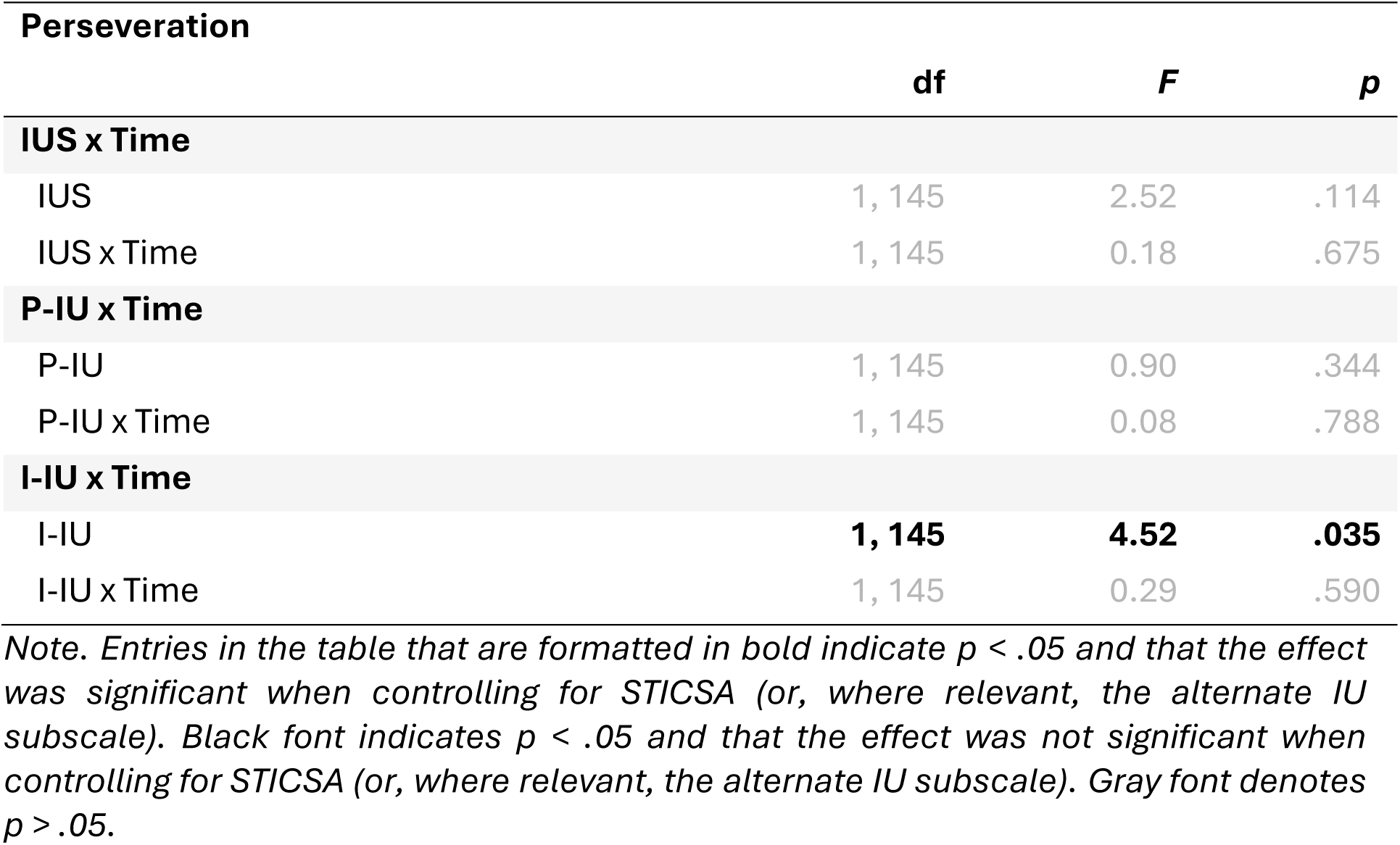
Results from MLMs Assessing Intolerance of Uncertainty, Prospective Intolerance of Uncertainty, and Inhibitory Intolerance of Uncertainty Main Effects and Interactions with Time on Perseveration.

Individual differences in P-IU were significantly associated with differential inverse temperature at Time 1 vs. Time 2 both when entered into the model alone (*p* = .021, see Table 5) as well as when controlling for trait anxiety (P-IU x Time: *F*(1, 357) = 5.45, *p* = .020; STICSA x Time: *F*(1, 357) = 0.06, *p* = .808) and for the inhibitory subscale (P-IU x Time: *F*(1, 435) = 4.25, *p* = .040; I-IU x Time: *F*(1, 435) = 0.70, *p* = .405). The simple slopes suggest that higher prospective IU is associated with higher inverse temperatures at time two (see Figure 5).

**Figure 5.**
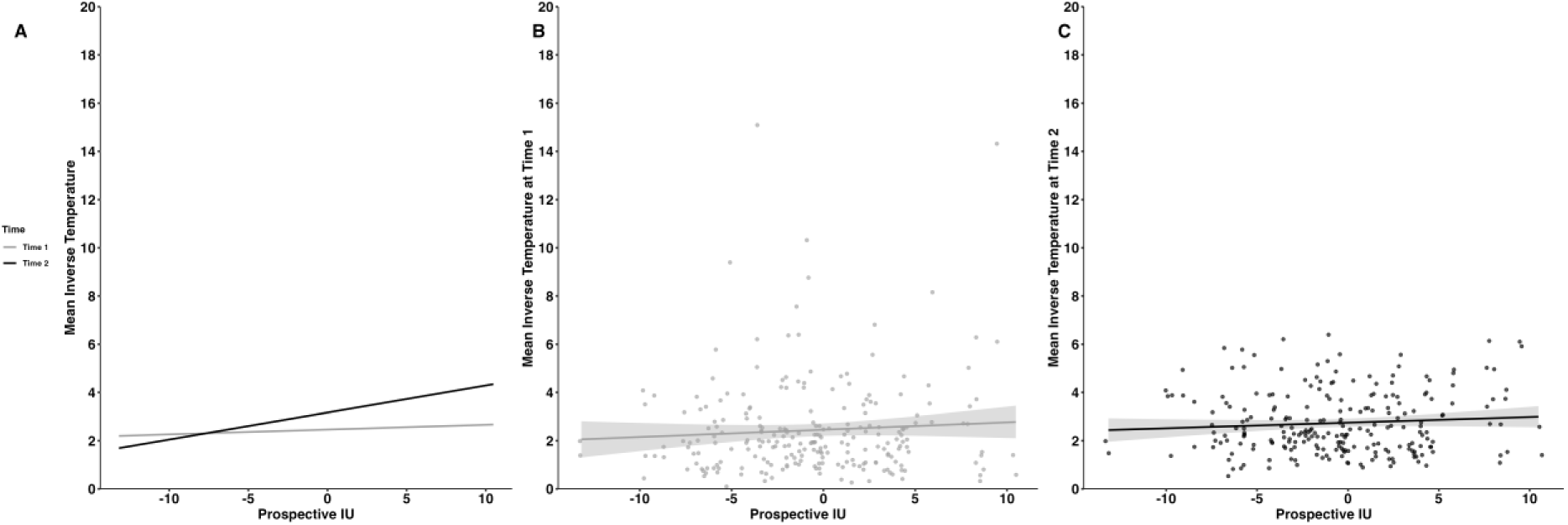
(A) Simple Slopes Depicting Association Between Prospective Intolerance of Uncertainty (P-IU; mean-centred) and Inverse Temperature per Outcome and (B) its Corresponding Scatterplot for Time 1 and (C) Time 2. Higher P-IU was associated with higher inverse temperature at both times. Shaded regions represent 95% confidence intervals.

**Table 5.**
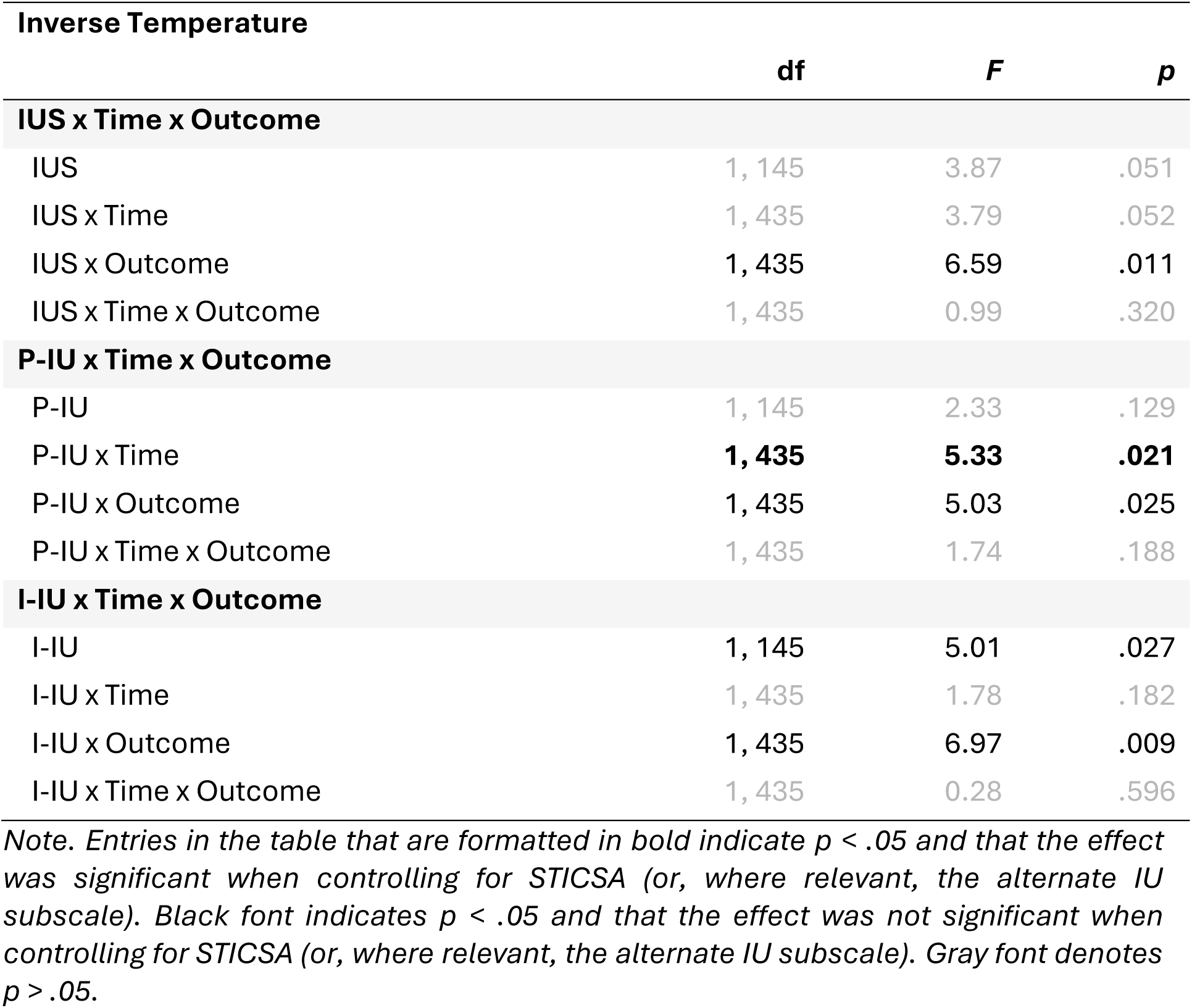
Results from MLMs Assessing Intolerance of Uncertainty, Prospective Intolerance of Uncertainty, and Inhibitory Intolerance of Uncertainty Main Effects and Interactions with Time (and Outcome) on Inverse Temperature.

Overall, our results indicate that higher levels of intolerance of uncertainty were associated with increased sensitivity to positive but not negative outcomes. Further, higher levels of inhibitory IU were associated with higher accuracy and fewer perseverative errors, that is, participants with increased inhibitory intolerance of uncertainty were more likely to select the ’best’ (most likely to be rewarding) choice and less likely to perseverate on the previously ‘best’ choice following reversal. Lastly, the prospective subscale was associated with higher inverse temperature parameter across sessions, with the association being stronger at time two compared to time one. Each of these effects was significant after controlling for trait anxiety, and the effects of the inhibitory and prospective subscales were specific to the subscale.

## Discussion

Intolerance of uncertainty is a transdiagnostic measure of psychopathology that has previously been associated with individual differences in conditioning and aversive learning paradigms. Here we investigate the role of intolerance of uncertainty in adaptive behaviour using a reversal learning task. We find that intolerance of uncertainty influences behaviour, as indexed by behavioural and latent (computational modelling) measures of task performance. Our preliminary evidence suggests that intolerance of uncertainty is associated with increased sensitivity to reinforcement when receiving positive feedback, but not when receiving negative feedback. The inhibitory subscale of intolerance of uncertainty was associated with improved performance on the task both overall (as indexed by improved accuracy) and immediately following a change in reward contingencies (as indexed by reduced perseveration). Importantly, these relationships were both stable across time and significant over and above effects of trait anxiety, and specific to the inhibitory subscale of intolerance of uncertainty. By contrast, the prospective subscale was positively associated with the extent to which reinforcement history reduced choice stochasticity in the second testing session, reflecting an inverse temperature parameter that may itself index learning across the sessions.

The specificity of our results for intolerance of uncertainty and its inhibitory and prospective subscales suggests they contribute to a behavioural phenotype positively associated with enhanced performance in environments where flexible behaviour is advantageous. More specifically, overall intolerance of uncertainty was positively correlated with reinforcement sensitivity for wins, pointing to an increased efficiency of confirmatory evidence in driving behaviour. Additionally, the inhibitory subscale of intolerance of uncertainty was associated with greater responsivity to change in the environment as indexed by increased accuracy overall combined with a decrease in perseverative responding following reversals. These behavioural results indicate that higher inhibitory intolerance of uncertainty may support increased tuning to evidence of goal-relevant state changes in the environment. Finally, higher prospective IU was associated with higher inverse temperature, but only during the second testing session. The inverse temperature parameter quantifies the extent to which choice behaviour is driven by expected value as determined by the history of experienced reinforcement, as opposed to exploration of the available options. Therefore, this finding may reflect a benefit in the transfer of learning across time points, whereby those higher in prospective IU benefit more from the prior experience of the reinforcement schedule at time point one. To summarise, we observe a phenotype characterising individuals with higher intolerance of uncertainty as being more responsive to changing environments. This is in line with findings from previous literature, including associations between the inhibitory subscale with threat avoidance (for review see, Morriss et al., 2021) and reward approach (Luhmann et al., 2011).

These findings also sit alongside previous research from the Pavlovian threat learning literature demonstrating that individual differences in intolerance of uncertainty are specifically associated with reversal of threat and safety contingencies (Mertens & Morriss, 2021; Morriss et al., 2019). Alongside this, the current study revealed that higher inhibitory intolerance of uncertainty was associated with higher reinforcement sensitivity for wins. This intolerance of uncertainty-related effect follows recent literature evidencing how higher intolerance of uncertainty is associated with the greater tendency to seek out smaller certain rewards (Ciria et al., 2021; Luhmann et al., 2011; Radell et al., 2016, 2018; Tanovic, Hajcak, et al., 2018). Taken together, and given the specificity of our findings with the inhibitory subscale, these results point to a potentially important role of the inhibitory intolerance of uncertainty (i.e. paralysis in the face of uncertainty) in the motivation to avoid uncertainty-related distress by seeking certain rewards and relying on confirmatory evidence in individuals with high inhibitory intolerance of uncertainty. This may in turn enhance contingency updating processes and related behaviours in environments where these strategies are tuned to the reinforcement regime, as is the case in the serial reversal task used here.

Overall, these findings extend our understanding of how intolerance of uncertainty influences learning in environments that are changeable (for review see Morriss et al. 2021) and where choices or actions result in different outcomes (e.g. gain, loss). Future studies could extend this line of research by manipulating different parameters of uncertainty (Morriss, 2023; Sandhu et al., 2023), volatility, task difficulty, and information availability from choice feedback, as well as measure different read outs to further probe underlying psychological processes (e.g. physiological and neural markers) (see recent special issue, Morriss, Abend, et al 2023). The extent to which intolerance of uncertainty modulates learning and choices in samples with different mental health conditions will also elucidate whether the effects observed here are specific to non-clinical samples or change on the basis of relevant symptom dimensions (e.g. anxiety and depression) (Shihata et al., 2016).

This study benefits from several key strengths. Firstly, deploying this study online meant that we could collect data from a large number of participants without the limitations of lab-based testing. To ensure our participants produced high quality data, we sourced participants from the online recruitment platform Prolific. Multiple publications demonstrate that participants on Prolific provide more high quality data than other popular online recruitment services such as Amazon Mechanical Turk and Qualtrics (Douglas et al., 2023; Peer et al., 2022). We also implemented multiple attentional and careless/insufficient effort checks in the reversal learning task and psychometric questionnaires, and only analysed data from participants who passed the complete set of checks. Secondly, our longitudinal design meant we could test whether our effects of intolerance of uncertainty on task performance varied as a function of time (and/or practice effects). Although we observed increases in reversal learning performance in our population from time one to time two, the relationship between intolerance of uncertainty and task performance was stable across testing sessions, suggesting these results are representative of stable individual differences. The only exception was the learning benefit in high perspective IU individuals in time two described above. Additionally, by using computational modelling, we examined latent contributions to task performance (such as learning rate and reinforcement sensitivity). One benefit of this approach is that putative functions can be mapped into other domains – including neural data – and there is a large body of existing literature that maps these functions to specific neural circuits (Chase et al., 2015; O’Doherty et al., 2015). Future work could aim to investigate whether interactions between individual differences in intolerance of uncertainty and task performance during reversal learning are associated with differences in neural recruitment across both typically developing individuals and clinical populations.

Our study also presented several methodological challenges. When designing the procedure for our reversal learning task we adopted approaches that aimed to optimise the performance of participants in an online setting. To incentivise performance on the reversal learning task we informed participants that we would give them a bonus payment if they did “well on the task” and “paid attention”. We operationalised “doing well” and “paying attention” as not exiting from the full screen mode which the task entered when initialised, and by not abstaining from making a choice on five or more consecutive trials or eight trials in total. Participants were informed of these conditions. The bonus for meeting these task performance criteria in the first session was an additional award of 50p, and in the second session a bonus of £1.25 was provided, and both bonus payouts were described to participant prior to the start of both task sessions. The decision to have a larger bonus pay-out in the second session was made because we wanted to maximise the number of participants who returned for the second session. However, it may also have had unintended consequences for the performance of participants in each session. For instance, there is evidence from perceptual learning studies that magnitude of reward has significant effects on task performance across trial-, block-, and session-levels (Zhang et al., 2018). Thus, we are unable to disambiguate whether between-session differences in task performance are due to practice effects, performance incentive, or an interaction between the two. This point is particularly pertinent for the observed interaction between P-IU and time for the inverse temperature parameter, which had a stronger association at time two compared to time one; it is therefore conceivable this interaction is at least partially influenced by the unequal bonus payment structure across sessions.

In conclusion, this paper provides novel evidence describing how individual differences in intolerance of uncertainty modulate behavioural flexibility in environments that are changeable and where choices or actions result in different outcomes (e.g. gain, loss). These findings contribute and extend our understanding of how intolerance of uncertainty is involved in learning and motivation in instrumental action, beyond the domains of Pavlovian conditioning and threat, and open up exciting opportunities for future research examining whether these IU-related behaviours mature across the lifespan and underpin mental health concerns.

## Acknowledgments

This research was supported by the Magdalen Vernon PhD Studentship of the School of Psychology and Clinical Language Sciences, University of Reading, awarded to Brendan Williams, a Gorilla Grant (Cauldron Science) awarded to Brendan Williams, and by the Centre for Integrative Neuroscience and Neurodynamics (CINN), University of Reading. The authors would like to thank our students and collaborators in CINN for their advice and support in setting up the study.

## Competing Interests

The authors declare no competing interests.

## Author Contributions

Brendan Williams: Conceptualization, Methodology, Software, Validation, Formal analysis, Investigation, Resources, Data Curation, Writing - Original Draft, Writing - Review & Editing, Visualization, Project administration, Funding acquisition. Claudia Rodriguez-Sobstel: Methodology, Formal analysis, Data Curation, Writing - Original Draft, Writing - Review & Editing, Visualization. Lily FitzGibbon: Conceptualization, Methodology, Software, Validation, Writing - Review & Editing. Jayne Morriss: Methodology, Writing - Original Draft, Writing - Review & Editing, Supervision, Project administration. Anastasia Christakou Conceptualization, Methodology, Resources, Writing - Review & Editing, Supervision, Project administration, Funding acquisition.

## Notes

### Competing Interest Statement

The authors have declared no competing interest.

