## Supplementary Materials for "The influence of trait intolerance of uncertainty on behavioural flexibility"

### Appendix A: Additional Participant Information

Ineligible participants were filtered out from having our study advertised to them using Prolific's participant filters (see supplementary materials for details). We identified 2,997 eligible participants who had been active over the past 90 days. During consent, participants confirmed whether they had any clinically diagnosed mental health disorders, whether they smoked, and if they took any psychoactive medication or illicit drugs, and any participants reporting yes were excluded. Participants were required to complete this study using either a laptop or desktop personal computer, due to the need of a keyboard to record keypresses during the reversal learning task. 257 participants recruited through Prolific consented to take part in the study. 251 participants completed the experiment in the first part of the study (Time 1). 2 participants were excluded after Time 1 because they failed instructional attention check questions, and 14 were excluded because they failed nonsensical attention check questions or careless / insufficient effort (C/IE) responding checks in the reversal learning task. 222 participants who were eligible for the second part of the study (Time 2) completed the experiment. 1 participant was excluded after Time 2 because they failed instructional attention check questions, and 8 were excluded because they failed nonsensical attention check questions or the C/IE responding checks.

**Supplementary Table 1**

*Participant Demographics*

| | **Learners Only (N = 145)** | **Learners and Completers (N = 118)** |
|---|---|---|
| | **M/N (SD/%)** | **M/N (SD/%)** |
| **Age** | 35.34 (13.33) | 37.64 (13.36) |
| **Sex** | | |
| Female | 93 (64.14) | 81 (68.64) |
| Male | 52 (35.86) | 37 (31.36) |
| **Country of Birth** | | |
| Belgium | 1 (0.69) | 0 (0.00) |
| Bulgaria | 3 (2.07) | 2 (1.69) |
| China | 2 (1.38) | 1 (0.85) |
| Finland | 1 (0.69) | 0 (0.00) |
| France | 1 (0.69) | 1 (0.85) |
| Greece | 4 (2.76) | 2 (1.69) |
| India | 1 (0.69) | 1 (0.85) |
| Lithuania | 1 (0.69) | 1 (0.85) |
| Nepal | 1 (0.69) | 0 (0.00) |
| Palestinian Territory | 1 (0.69) | 0 (0.00) |
| Poland | 2 (1.38) | 2 (1.69) |
| Portugal | 2 (1.38) | 2 (1.69) |

| | Learners Only (N = 145) | Learners and Completers (N = 118) |
|---|---|---|
| | **M/N (SD/%)** | **M/N (SD/%)** |
| Singapore | 1 (0.69) | 1 (0.85) |
| South Africa | 1 (0.69) | 1 (0.85) |
| Turkey | 1 (0.69) | 1 (0.85) |
| United Kingdom | 121 (83.45) | 102 (86.44) |
| United States | 1 (0.69) | 1 (0.85) |
| **Current Country of Residence** | | |
| Greece | 3 (2.07) | 2 (1.69) |
| Poland | 1 (0.69) | 1 (0.85) |
| Portugal | 3 (2.07) | 3 (2.54) |
| United Kingdom | 138 (95.17) | 112 (94.92) |
| **First Language** | | |
| Bengali | 1 (0.69) | 1 (0.85) |
| Bulgarian | 3 (2.07) | 2 (1.69) |
| Chinese | 2 (1.38) | 1 (0.85) |
| Dutch | 1 (0.69) | 0 (0.00) |
| English | 125 (86.21) | 104 (88.14) |
| Finnish | 1 (0.69) | 0 (0.00) |
| French | 1 (0.69) | 1 (0.85) |
| Greek | 4 (2.76) | 2 (1.69) |
| Lithuanian | 1 (0.69) | 1 (0.85) |
| Polish | 1 (0.69) | 1 (0.85) |
| Portuguese | 3 (2.07) | 3 (2.54) |
| Turkish | 1 (0.69) | 1 (0.85) |
| Information not provided | 1 (0.69) | 1 (0.85) |
| **Nationality** | | |
| Belgium | 1 (0.69) | 0 (0.00) |
| Bulgaria | 2 (1.38) | 2 (1.69) |
| Finland | 1 (0.69) | 0 (0.00) |
| France | 1 (0.69) | 1 (0.85) |
| Greece | 4 (2.76) | 2 (1.69) |
| Ireland | 1 (0.69) | 1 (0.85) |
| Lithuania | 1 (0.69) | 1 (0.85) |
| Nepal | 1 (0.69) | 0 (0.00) |
| Poland | 2 (1.38) | 2 (1.69) |
| Portugal | 3 (2.07) | 3 (2.54) |
| Singapore | 1 (0.69) | 1 (0.85) |
| South Africa | 1 (0.69) | 1 (0.85) |
| Turkey | 1 (0.69) | 1 (0.85) |
| United Kingdom | 124 (85.52) | 102 (86.44) |
| United States | 1 (0.69) | 1 (0.85) |
| **Employment Status** | | |
| Due to start a new job within the next month | 2 (1.38) | 1 (0.85) |

| | Learners Only (N = 145) | Learners and Completers (N = 118) |
|---|---|---|
| | **M/N (SD/%)** | **M/N (SD/%)** |
| Full-Time | 72 (49.66) | 62 (52.54) |
| Not in paid work (e.g. homemaker, retired or disabled) | 13 (8.97) | 10 (8.47) |
| Part-Time | 27 (18.62) | 24 (20.34) |
| Unemployed (and job seeking) | 10 (6.90) | 7 (5.93) |
| Other | 10 (6.90) | 9 (7.63) |
| Information not provided | 11 (7.59) | 5 (4.24) |
| **Student Status** | | |
| Yes | 31 (21.38) | 18 (15.25) |
| No | 103 (71.03) | 94 (79.66) |
| Information not provided | 11 (7.59) | 6 (5.08) |

*Note*. M = mean, N = number, SD = standard deviation.

### Appendix B: Computational modelling

The first model in the softmax family is a model-free reinforcement learning model with a single learning rate parameter ($\alpha$) and inverse temperature parameter ($\beta$). In this model the expected value ($V$) of choice $k$ on trial $t$ ($V_t^k$) is updated for the next trial ($t + 1$) by adding the product of the learning rate and the prediction error ($\lambda_t - V_t^k$), which is the difference between the actual ($\lambda$) and expected value (eq. 1).

$$V_{t+1}^k = V_t^k + \alpha\left(\lambda_t - V_t^k\right) \quad (1)$$

The probability of making choice $k$ on trial $t$ is determined by the softmax choice rule (eq. 2), and the inverse temperature parameter ($\beta$) determines the extent to which choices are based on expected value estimates. When $\beta = 0$, choices would be made completely at random; when $\beta = \infty$ the choice with the largest expected value would be deterministically chosen.

$$p_t^k = \frac{e^{\beta V_t^k}}{\sum_{i=1}^{K} e^{\beta V_t^i}} \quad (2)$$

In model one, expected values are updated at the same rate for positive and negative prediction errors. However, there is evidence that suggests that positive and negative prediction errors have asymmetric update rates with different sensitivities to wins and losses (Niv et al., 2012). Therefore, in model two included separate learning rates for wins and losses (eq. 3).

$$V_{t+1}^k = V_t^k + \alpha^{win/loss}\left(\lambda_t - V_t^k\right) \quad (3)$$

In models three and four, because choices may have dissociable sensitivities for previous wins and losses, separate inverse temperature parameters are used based on whether a win or loss was experienced on the previous trial (eq. 4); model three used a single learning rate for updating expected value (eq. 1) and model three used dual learning rates (eq. 3).

$$p_t^k = \frac{e^{\beta_{win/loss} V_t^k}}{\sum_{i=1}^{K} e^{\beta_{win/loss} V_t^i}} \quad (4)$$

Models five to eight updated the expected value choice $k$ on trial $t$ with single (models five and seven) or dual learning rates (models six and eight) and had single (models five and six) or dual (models seven and eight) inverse temperature parameters. However, expected values for the unchosen options ($k_{unchosen}$) on trial $t$ were also updated using the inverse of the actual ($\lambda$) outcome on trial $t$ (eq. 5 for models five and seven; eq. 6 for models six and eight).

$$V_{t+1}^{k_{unchosen}} = V_t^{k_{unchosen}} + \alpha\left((-\lambda_t) - V_t^{k_{unchosen}}\right) \quad (5)$$

$$V_{t+1}^{k_{unchosen}} = V_t^{k_{unchosen}} + \alpha^{+/-}\left((-\lambda_t) - V_t^{k_{unchosen}}\right) \quad (6)$$

Lastly, models nine to twelve updated chosen ($k$) choices on trial $t$ as in models five to eight (respectively), however the update of unchosen ($k_{unchosen}$) options was weighted by a discount parameter ($\kappa$) for models nine and eleven (eq. 7) and ten and twelve (eq. 8) respectively.

$$V_{t+1}^{k_{unchosen}} = V_t^{k_{unchosen}} + \kappa\alpha\left((-\lambda_t) - V_t^{k_{unchosen}}\right) \tag{7}$$

$$V_{t+1}^{k_{unchosen}} = V_t^{k_{unchosen}} + \kappa\alpha^{win/loss}\left((-\lambda_t) - V_t^{k_{unchosen}}\right) \tag{8}$$

For the reinforcement sensitivity family of models there is no inverse temperature parameter included in the softmax choice rule (eq. 9). Instead, a reinforcement sensitivity ($\rho$) parameter is used. In the softmax family of models the inverse temperature parameter determines choice stochasticity by determining the extent to which choices are driven by expected values, while the reinforcement sensitivity parameter does this by determining the maximum difference between expected values, which places a lower bound on choice stochasticity (Waltmann et al., 2022).

$$p_t^k = \frac{e^{V_t^k}}{\sum_{i=1}^{K} e^{V_t^i}} \tag{9}$$

The reinforcement sensitivity family of models broadly follow the softmax family of models with respect to how expected values are updated. The first model in the reinforcement sensitivity family is a model-free reinforcement learning model with a single learning rate parameter ($\alpha$) and reinforcement sensitivity parameter ($\rho$). When calculating a prediction error on trial $t$ in the reinforcement sensitivity family of models the actual outcome is scaled by the reinforcement sensitivity parameter ($\rho\lambda_t$) before subtracting the expected value ($V$) of choice $k$ on trial $t$ ($V_t^k$). Expected values are updated for the next trial ($t+1$) by adding the product of the learning rate and the prediction error ($\rho\lambda_t - V_t^k$) (eq. 10).

$$V_{t+1}^k = V_t^k + \alpha\left(\rho\lambda_t - V_t^k\right) \tag{10}$$

The second model in the reinforcement sensitivity family had separate learning rates for wins and losses (eq. 11). The third and fourth models included separate reinforcement sensitivities for wins and losses, coupled with symmetric (eq. 12) and asymmetric (eq. 13) learning rates for wins and losses, respectively.

$$V_{t+1}^k = V_t^k + \alpha^{win/loss}\left(\rho\lambda_t - V_t^k\right) \tag{11}$$

$$V_{t+1}^k = V_t^k + \alpha\left(\rho_{win/loss}\lambda_t - V_t^k\right) \tag{12}$$

$$V_{t+1}^k = V_t^k + \alpha^{win/loss}\left(\rho_{win/loss}\lambda_t - V_t^k\right) \tag{13}$$

Models five to eight updated expected values for both chosen and unchosen choices on trial $t$. The expected value of choice $k$ on trial $t$ was updated using a single (models five and seven) or separate learning rates (models six and eight). Models five and six included a single reinforcement sensitivity parameter, while models seven and eight included separate reinforcement sensitivities for wins and losses. Expected values for unchosen options ($k_{unchosen}$) on trial $t$ were also updated using the inverse of the actual ($\lambda$) outcome on trial $t$ (eqs. 14 and 16 for models five and seven; eqs. 15 and 17 for models six and eight).

$$V_{t+1}^{k_{unchosen}} = V_t^{k_{unchosen}} + \alpha\left(\rho(-\lambda_t) - V_t^{k_{unchosen}}\right) \tag{14}$$

$$V_{t+1}^{k_{unchosen}} = V_t^{k_{unchosen}} + \alpha^{win/loss}\left(\rho(-\lambda_t) - V_t^{k_{unchosen}}\right) \tag{15}$$

$$V_{t+1}^{k_{unchosen}} = V_t^{k_{unchosen}} + \alpha\left(\rho_{win/loss}(-\lambda_t) - V_t^{k_{unchosen}}\right) \tag{16}$$

$$V_{t+1}^{k_{unchosen}} = V_t^{k_{unchosen}} + \alpha^{win/loss}\left(\rho_{win/loss}(-\lambda_t) - V_t^{k_{unchosen}}\right) \tag{17}$$

Lastly, models nine to twelve updated chosen ($k$) choices on trial $t$ as in models five to eight (respectively), however the update of unchosen ($k_{unchosen}$) options was weighted by a discount parameter ($\kappa$) for models nine and eleven (eqs. 18 and 20) and ten and twelve (eqs. 19 and 21) respectively.

$$V_{t+1}^{k_{unchosen}} = V_t^{k_{unchosen}} + \kappa\alpha\left(\rho(-\lambda_t) - V_t^{k_{unchosen}}\right) \tag{18}$$

$$V_{t+1}^{k_{unchosen}} = V_t^{k_{unchosen}} + \kappa\alpha^{win/loss}\left(\rho(-\lambda_t) - V_t^{k_{unchosen}}\right) \tag{19}$$

$$V_{t+1}^{k_{unchosen}} = V_t^{k_{unchosen}} + \kappa\alpha\left(\rho_{win/loss}(-\lambda_t) - V_t^{k_{unchosen}}\right) \tag{20}$$

$$V_{t+1}^{k_{unchosen}} = V_t^{k_{unchosen}} + \kappa\alpha^{win/loss}\left(\rho_{win/loss}(-\lambda_t) - V_t^{k_{unchosen}}\right) \tag{21}$$

### Appendix C: Questionnaire Frequency Distributions

**Supplementary Figure 1**

*Histograms Depicting Distributions of (**A**) Intolerance of Uncertainty Scores (IUS) Across Time 1, Time 2, and Time 3, (**B**) IUS Averaged Across Times 1 and 2 for Non-Completers and Across Times 1, 2 and 3 for Completers, (**C**) Prospective Intolerance of Uncertainty (P-IU) Scores Across Time 1, Time 2, and Time 3, (**D**) P-IU Scores Averaged Across Times 1 and 2 for Non-Completers and Across Times 1, 2 and 3 for Completers, (**E**) Inhibitory Intolerance of Uncertainty (I-IU) Scores Across Time 1, Time 2, and Time 3, (**F**) I-IU Scores Averaged Across Times 1 and 2 for Non-Completers and Across Times 1, 2 and 3 for Completers, and (**G**) STICSA Scores.*

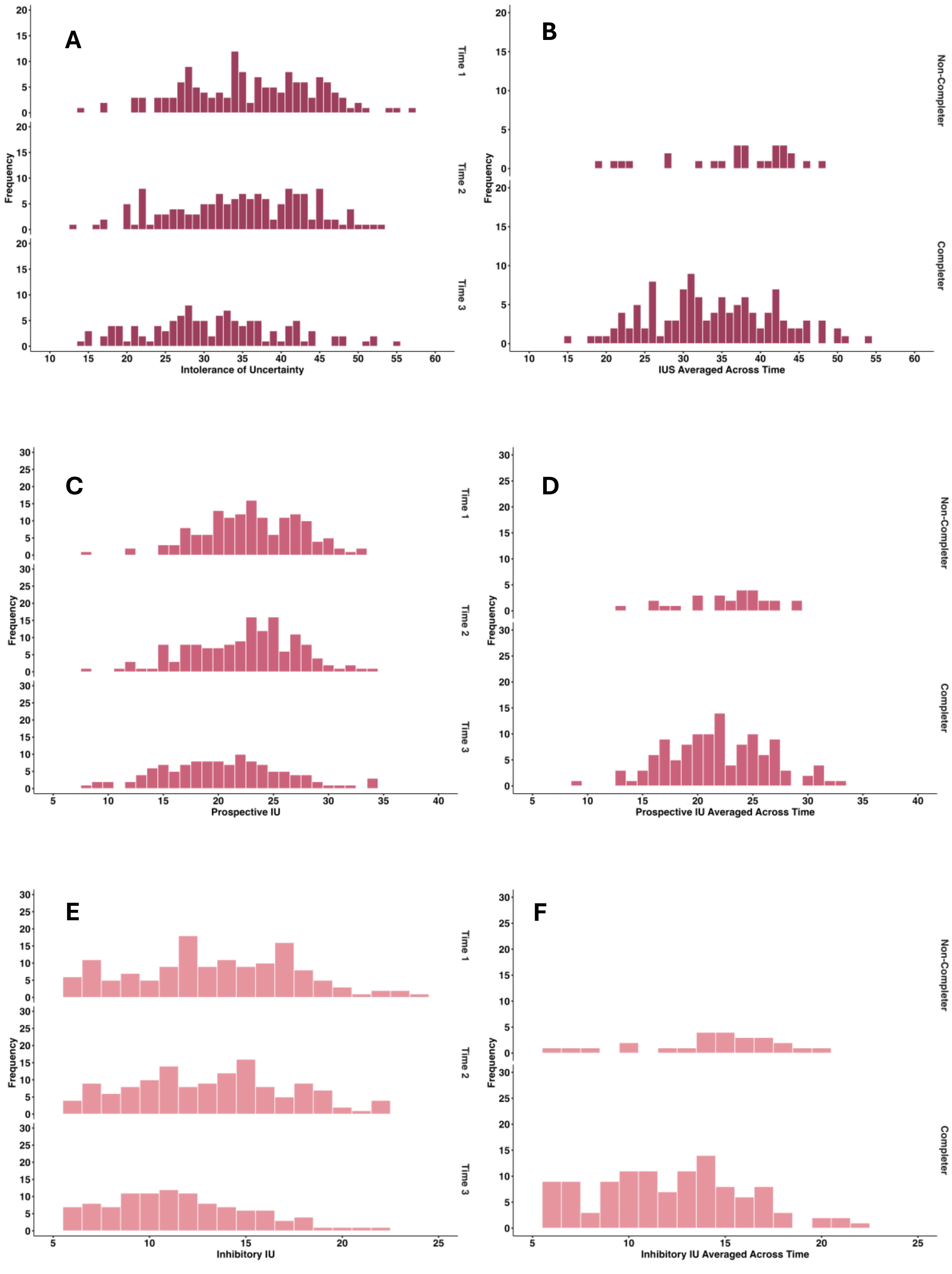

A
B
C
D
E
F
Frequency
Time 1
Time 2
Time 3
Non-Completer
Completer
Intolerance of Uncertainty
IUS Averaged Across Time
Prospective IU
Prospective IU Averaged Across Time
Inhibitory IU
Inhibitory IU Averaged Across Time

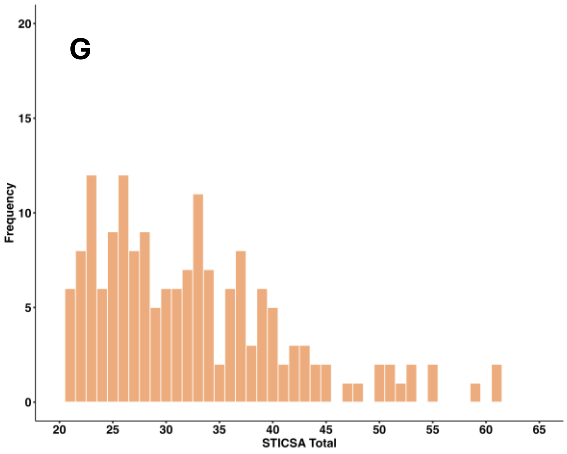
G
Frequency
20
15
10
5
0
20
25
30
35
40
45
50
55
60
65
STICSA Total

### Appendix D: Questionnaire Descriptive Statistics

**Supplementary Table 2**

*Descriptive Statistics for IUS, P-IU, I-IU and STICSA.*

| | N | Mean | SD | Range | | Skew | Kurtosis |
|---|---|---|---|---|---|---|---|
| | | | | *Min* | *Max* | | |
| **IUS** | | | | | | | |
| Time 1 | 145 | 36.03 | 8.45 | 14 | 57 | -0.08 | -0.49 |
| Time 2 | 145 | 34.79 | 8.72 | 13 | 53 | -0.21 | -0.69 |
| Time 3 | 118 | 31.34 | 8.98 | 14 | 55 | 0.28 | -0.34 |
| Average Across Time | 145 | 34.39 | 8.16 | 15 | 54.33 | -0.03 | -0.70 |
| **P-IU** | | | | | | | |
| Time 1 | 145 | 23.01 | 4.48 | 8 | 33 | -0.25 | 0.07 |
| Time 2 | 145 | 22.34 | 4.87 | 8 | 34 | -0.29 | -0.18 |
| Time 3 | 118 | 20.45 | 5.52 | 8 | 34 | 0.17 | -0.23 |
| Average Across Time | 145 | 22.09 | 4.51 | 9.33 | 33 | -0.04 | -0.29 |
| **I-IU** | | | | | | | |
| Time 1 | 145 | 13.02 | 4.52 | 5 | 24 | 0.02 | -0.71 |
| Time 2 | 145 | 12.44 | 4.51 | 5 | 22 | 0.06 | -0.84 |
| Time 3 | 118 | 10.89 | 4.04 | 5 | 22 | 0.40 | -0.45 |
| Average Across Time | 145 | 12.30 | 4.10 | 5 | 22.33 | 0.03 | -0.78 |
| **STICSA** | 118 | 32.09 | 8.23 | 21 | 59 | 0.89 | 0.45 |

### Appendix E: Behavioural and Computational Data Frequency Distributions

**Supplementary Figure 2**

*Histograms Depicting Distributions of Behavioural and Computational Modelling Data Across Time 1 and Time 2.*

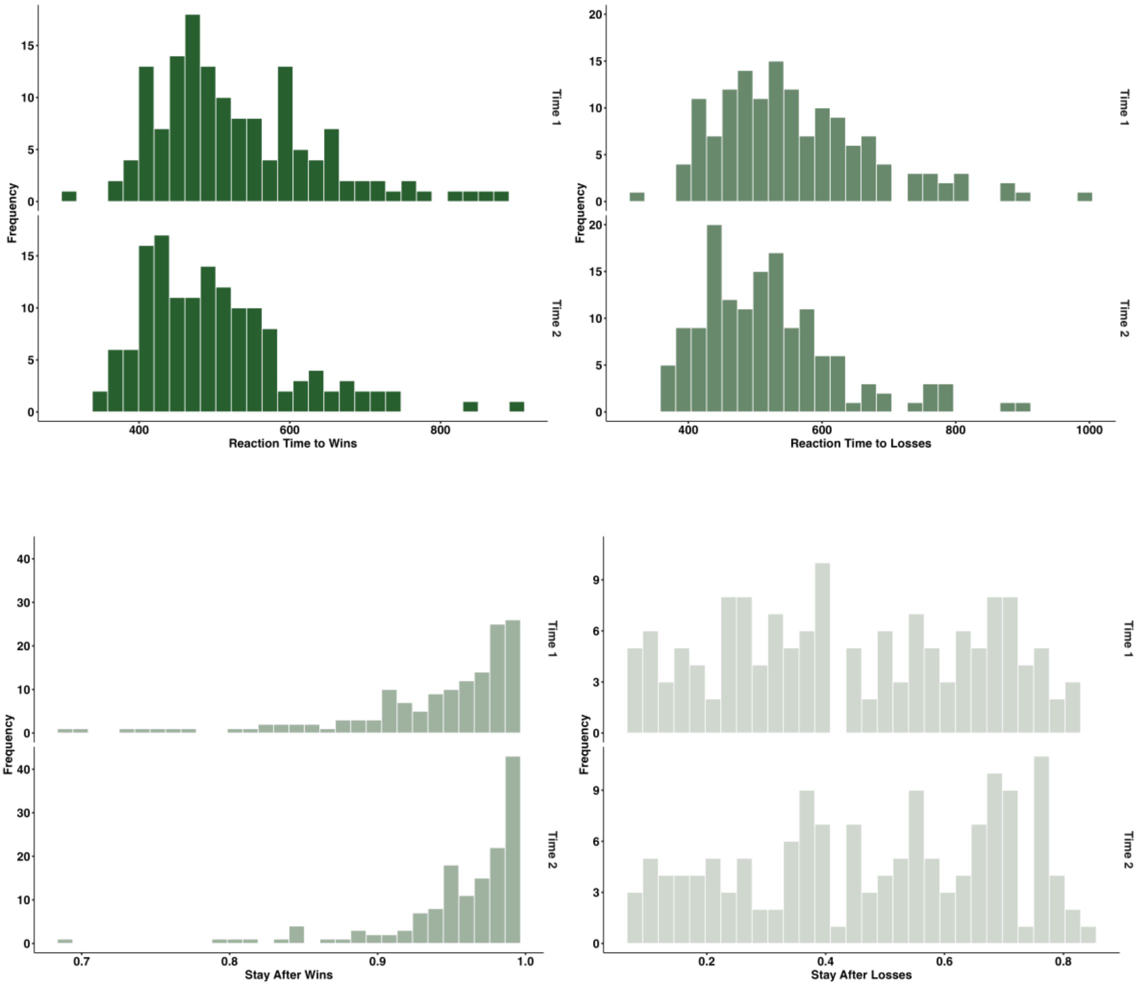

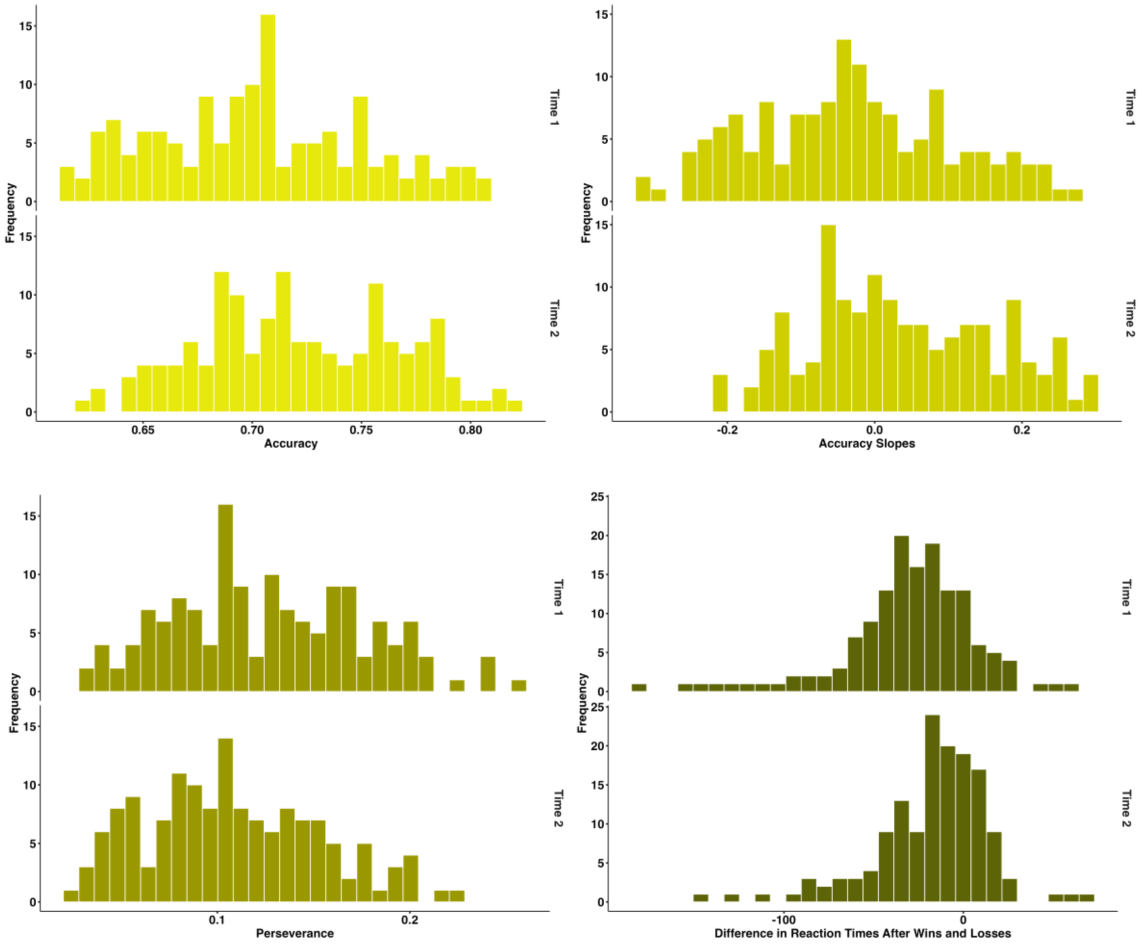
Frequency
Time 1
Time 2
Accuracy
Accuracy Slopes
Perseverance
Difference in Reaction Times After Wins and Losses

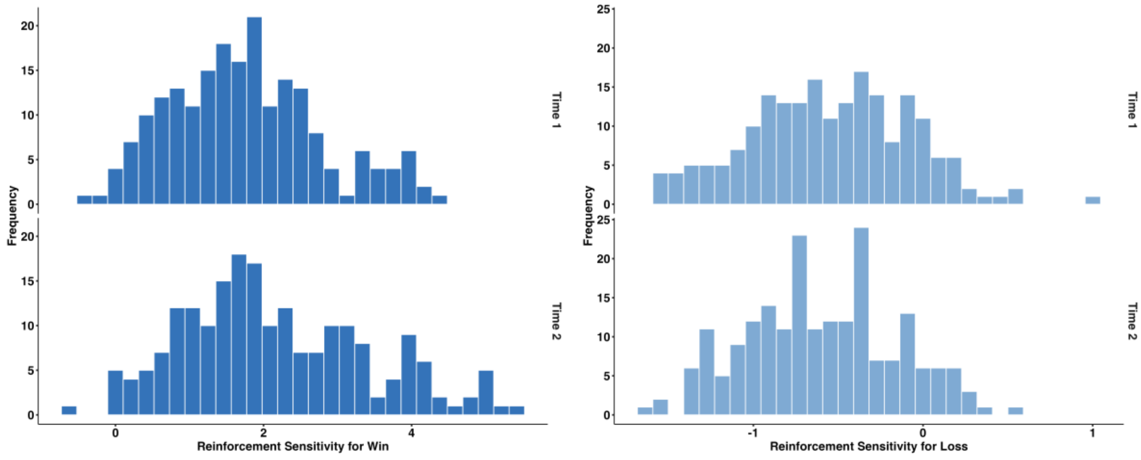
Frequency
Time 1
Time 2
Reinforcement Sensitivity for Win
Reinforcement Sensitivity for Loss

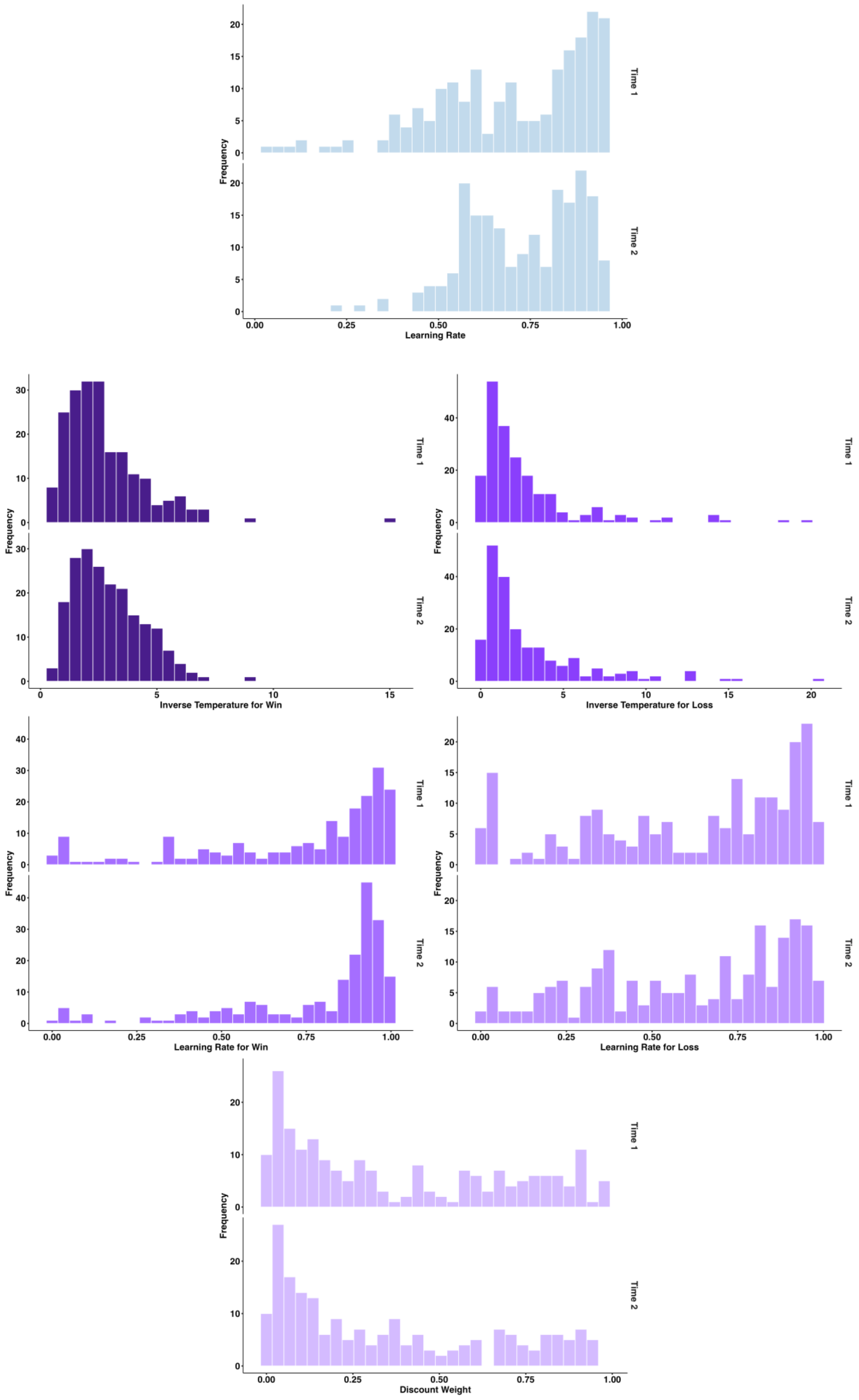
Frequency
Time 1
Time 2
Learning Rate
Inverse Temperature for Win
Inverse Temperature for Loss
Learning Rate for Win
Learning Rate for Loss
Discount Weight

### Appendix F: Behavioural and Computational Data Descriptive Statistics

**Supplementary Table 3**

*Descriptive Statistics for Behavioural and Computational Modelling Variables*

| | N | Mean | SD | Range | | Skew | Kurtosis |
|---|---|---|---|---|---|---|---|
| | | | | *Min* | *Max* | | |
| **Behavioural Variables** | | | | | | | |
| **Reaction Time** | | | | | | | |
| **Learners Only** | | | | | | | |
| **Time 1** | | | | | | | |
| Win | 145 | 529.60 | 106.69 | 316.19 | 875.03 | 0.93 | 0.73 |
| Loss | 145 | 559.83 | 118.07 | 318.09 | 986.91 | 0.96 | 0.94 |
| Time 1 Average | 290 | 544.71 | 113.35 | 316.19 | 986.91 | 0.97 | 0.97 |
| **Time 2** | | | | | | | |
| Win | 145 | 500.98 | 97.71 | 338.43 | 910.18 | 1.20 | 1.95 |
| Loss | 145 | 519.08 | 103.75 | 359.36 | 905.88 | 1.16 | 1.59 |
| Time 2 Average | 290 | 510.03 | 101.01 | 338.43 | 910.18 | 1.18 | 1.80 |
| **Win Average** | 290 | 515.29 | 103.13 | 316.19 | 910.18 | 1.06 | 1.25 |
| **Loss Average** | 290 | 539.45 | 112.81 | 317.10 | 986.91 | 1.07 | 1.27 |
| **Learners and Completers** | | | | | | | |
| **Time 1** | | | | | | | |
| Win | 118 | 533.56 | 113.07 | 316.19 | 857.03 | 0.87 | 0.42 |
| Loss | 118 | 562.84 | 124.79 | 317.10 | 986.91 | 0.92 | 0.68 |
| Time 1 Average | 236 | 548.20 | 119.72 | 316.19 | 986.91 | 0.92 | 0.70 |
| **Time 2** | | | | | | | |
| Win | 118 | 507.02 | 101.51 | 338.43 | 910.18 | 1.19 | 1.78 |
| Loss | 118 | 527.23 | 108.13 | 359.36 | 905.88 | 1.11 | 1.29 |
| Time 2 Average | 236 | 517.13 | 105.14 | 338.43 | 910.18 | 1.16 | 1.55 |
| **Win Average** | 236 | 520.29 | 108.04 | 316.19 | 910.18 | 1.03 | 1.01 |
| **Loss Average** | 236 | 545.04 | 117.87 | 317.10 | 986.91 | 1.03 | 1.04 |
| **Stay** | | | | | | | |
| **Learners Only** | | | | | | | |
| **Time 1** | | | | | | | |
| Win | 145 | 0.94 | 0.06 | 0.69 | 0.99 | -1.76 | 3.00 |
| Loss | 145 | 0.44 | 0.21 | 0.07 | 0.82 | 0.00 | -1.25 |
| Time 1 Average | 290 | 0.69 | 0.29 | 0.07 | 0.99 | -0.61 | -1.05 |
| **Time 2** | | | | | | | |
| Win | 145 | 0.96 | 0.05 | 0.69 | 0.99 | -2.35 | 7.17 |
| Loss | 145 | 0.48 | 0.22 | 0.07 | 0.83 | -0.26 | -1.14 |
| Time 2 Average | 290 | 0.72 | 0.28 | 0.07 | 0.99 | -0.74 | -0.78 |
| **Win Average** | 290 | 0.95 | 0.06 | 0.69 | 0.99 | -2.07 | 4.76 |
| **Loss Average** | 290 | 0.46 | 0.22 | 0.07 | .083 | -0.13 | -1.21 |
| **Learners and Completers** | | | | | | | |
| **Time 1** | | | | | | | |
| Win | 118 | 0.94 | 0.06 | 0.69 | 0.99 | -1.70 | 2.72 |
| Loss | 118 | 0.42 | 0.21 | 0.07 | 0.82 | 0.08 | -1.19 |

| | N | Mean | SD | Range | | Skew | Kurtosis |
|---|---|---|---|---|---|---|---|
| | | | | *Min* | *Max* | | |
| Time 1 Average | 236 | 0.68 | 0.30 | 0.07 | 0.99 | -0.56 | -1.15 |
| **Time 2** | | | | | | | |
| Win | 118 | 0.95 | 0.05 | 0.69 | 0.99 | -2.21 | 5.97 |
| Loss | 118 | 0.46 | 0.21 | 0.07 | 0.83 | -0.21 | -1.11 |
| Time 2 Average | 236 | 0.71 | 0.29 | 0.07 | 0.99 | -0.68 | -0.89 |
| **Win Average** | 236 | 0.94 | 0.06 | 0.69 | 0.99 | -1.96 | 4.10 |
| **Loss Average** | 236 | 0.44 | 0.21 | 0.07 | 0.83 | -0.06 | -1.17 |
| **Accuracy** | | | | | | | |
| **Learners Only** | | | | | | | |
| Time 1 | 145 | 0.70 | 0.05 | 0.61 | 0.81 | 0.19 | -0.78 |
| Time 2 | 145 | 0.72 | 0.05 | 0.62 | 0.82 | 0.06 | -0.86 |
| **Learners and Completers** | | | | | | | |
| Time 1 | 118 | 0.70 | 0.05 | 0.61 | 0.81 | 0.18 | -0.83 |
| Time 2 | 118 | 0.72 | 0.05 | 0.62 | 0.82 | 0.10 | -0.84 |
| **Accuracy Slopes** | | | | | | | |
| **Learners Only** | | | | | | | |
| Time 1 | 145 | -0.03 | 0.13 | -0.31 | 0.28 | 0.15 | -0.65 |
| Time 2 | 145 | 0.04 | 0.12 | -0.22 | 0.30 | 0.18 | -0.89 |
| **Learners and Completers** | | | | | | | |
| Time 1 | 118 | -0.03 | 0.13 | -0.30 | 0.28 | 0.12 | -0.74 |
| Time 2 | 118 | 0.04 | 0.12 | -0.22 | 0.30 | 0.20 | -0.84 |
| **Perseveration** | | | | | | | |
| **Learners Only** | | | | | | | |
| Time 1 | 145 | 0.13 | 0.05 | 0.03 | 0.26 | 0.26 | -0.60 |
| Time 2 | 145 | 0.11 | 0.05 | 0.03 | 0.22 | 0.34 | -0.61 |
| **Learners and Completers** | | | | | | | |
| Time 1 | 118 | 0.13 | 0.05 | 0.03 | 0.26 | 0.20 | -0.71 |
| Time 2 | 118 | 0.11 | 0.05 | 0.03 | 0.22 | 0.34 | -0.61 |
| **Difference in Reaction Times After Wins and Losses** | | | | | | | |
| **Learners Only** | | | | | | | |
| Time 1 | 145 | -30.22 | 37.65 | -180.56 | 64.52 | -1.08 | 2.59 |
| Time 2 | 145 | -18.09 | 32.29 | -148.08 | 69.05 | -0.96 | 2.42 |
| **Learners and Completers** | | | | | | | |
| Time 1 | 118 | -29.28 | 38.28 | -180.57 | 55.56 | -1.26 | 2.78 |
| Time 2 | 118 | -20.21 | 33.58 | -148.08 | 69.05 | -0.97 | 2.00 |
| **Reinforcement Sensitivity Computational Modelling Variables** | | | | | | | |
| **Reinforcement Sensitivity** | | | | | | | |
| **Learners Only** | | | | | | | |
| **Time 1** | | | | | | | |
| Win | 145 | 2.06 | 0.92 | 0.36 | 4.43 | 0.58 | -0.32 |
| Loss | 145 | -0.72 | 0.42 | -1.56 | 0.56 | 0.23 | -0.20 |
| Time 1 Average | 290 | 0.67 | 1.56 | -1.56 | 4.43 | 0.42 | -1.03 |
| **Time 2** | | | | | | | |
| Win | 145 | 2.52 | 1.15 | 0.42 | 5.50 | 0.59 | -0.50 |
| Loss | 145 | -0.73 | 0.40 | -1.62 | 0.58 | 0.31 | 0.02 |
| Time 2 Average | 290 | 0.90 | 1.84 | -1.62 | 5.50 | 0.52 | -0.94 |
| **Win Average** | 290 | 2.29 | 1.06 | 0.36 | 5.50 | 0.69 | -0.12 |
| **Loss Average** | 290 | -0.72 | 0.41 | -1.62 | 0.58 | 0.27 | -0.08 |
| **Learners and Completers** | | | | | | | |

| | N | Mean | SD | Range | | Skew | Kurtosis |
|---|---|---|---|---|---|---|---|
| | | | | *Min* | *Max* | | |
| **Time 1** | | | | | | | |
| Win | 118 | 1.98 | 0.84 | 0.36 | 4.01 | 0.57 | -0.29 |
| Loss | 118 | -0.70 | 0.43 | -1.56 | 0.56 | 0.17 | -0.20 |
| Time 1 Average | 236 | 0.64 | 1.50 | -1.56 | 4.01 | 0.37 | -1.10 |
| **Time 2** | | | | | | | |
| Win | 118 | 2.40 | 1.07 | 0.42 | 5.07 | 0.52 | -0.62 |
| Loss | 118 | -0.71 | 0.41 | -1.53 | 0.58 | 0.26 | -0.11 |
| Time 2 Average | 236 | 0.85 | 1.75 | -1.53 | 5.07 | 0.48 | -1.02 |
| **Win Average** | 236 | 2.19 | 0.98 | 0.36 | 5.07 | 0.65 | -0.21 |
| **Loss Average** | 236 | -0.70 | 0.42 | -1.56 | 0.58 | 0.21 | -0.13 |
| **Learning Rate** | | | | | | | |
| **Learners Only** | | | | | | | |
| Time 1 | 145 | 0.71 | 0.18 | 0.35 | 0.97 | -0.13 | -1.38 |
| Time 2 | 145 | 0.71 | 0.14 | 0.34 | 0.95 | -0.04 | -1.08 |
| **Learners and Completers** | | | | | | | |
| Time 1 | 118 | 0.73 | 0.17 | 0.38 | 0.97 | -0.17 | -1.41 |
| Time 2 | 118 | 0.73 | 0.14 | 0.43 | 0.95 | -0.04 | -1.24 |
| **Softmax Computational Modelling Variables** | | | | | | | |
| **Inverse Temperature** | | | | | | | |
| **Learners Only** | | | | | | | |
| **Time 1** | | | | | | | |
| Win | 145 | 2.61 | 1.31 | 0.46 | 6.40 | 0.83 | 0.05 |
| Loss | 145 | 2.69 | 2.98 | 0.10 | 18.36 | 2.69 | 8.50 |
| Time 1 Average | 290 | 2.65 | 2.30 | 0.10 | 18.36 | 3.04 | 13.62 |
| **Time 2** | | | | | | | |
| Win | 145 | 3.09 | 1.38 | 0.89 | 6.87 | 0.51 | -0.59 |
| Loss | 145 | 3.37 | 3.50 | 0.23 | 20.52 | 2.09 | 4.98 |
| Time 2 Average | 290 | 3.23 | 2.66 | 0.23 | 20.52 | 2.53 | 9.58 |
| **Win Average** | 290 | 2.85 | 1.36 | 0.46 | 6.87 | 0.66 | -0.33 |
| **Loss Average** | 290 | 3.03 | 3.27 | 0.10 | 20.52 | 2.36 | 6.46 |
| **Learners and Completers** | | | | | | | |
| **Time 1** | | | | | | | |
| Win | 118 | 2.51 | 1.28 | 0.53 | 6.40 | 1.05 | 0.58 |
| Loss | 118 | 2.37 | 2.54 | 0.10 | 15.09 | 2.72 | 8.78 |
| Time 1 Average | 236 | 2.44 | 2.01 | 0.10 | 15.09 | 2.83 | 12.02 |
| **Time 2** | | | | | | | |
| Win | 118 | 2.96 | 1.29 | 0.89 | 6.14 | 0.50 | -0.69 |
| Loss | 118 | 3.13 | 3.46 | 0.23 | 20.52 | 2.42 | 6.83 |
| Time 2 Average | 236 | 3.05 | 2.60 | 0.23 | 20.52 | 2.94 | 12.72 |
| **Win Average** | 236 | 2.73 | 2.36 | 0.53 | 6.40 | -0.21 | 0.08 |
| **Loss Average** | 236 | 2.75 | 3.05 | 0.10 | 20.52 | 2.66 | 8.51 |
| **Learning Rate** | | | | | | | |
| **Learners Only** | | | | | | | |
| **Time 1** | | | | | | | |
| Win | 145 | 0.77 | 0.23 | 0.09 | 1.00 | -1.03 | -0.06 |
| Loss | 145 | 0.69 | 0.24 | 0.11 | 0.99 | -0.60 | -0.99 |
| Time 1 Average | 290 | 0.73 | 0.24 | 0.09 | 1.00 | -0.80 | -0.61 |
| **Time 2** | | | | | | | |
| Win | 145 | 0.80 | 0.20 | 0.12 | 0.99 | -1.15 | 0.29 |

| | N | Mean | SD | Range | | Skew | Kurtosis |
|---|---|---|---|---|---|---|---|
| | | | | *Min* | *Max* | | |
| Loss | 145 | 0.64 | 0.25 | 0.12 | 0.99 | -0.34 | -1.22 |
| Time 2 Average | 290 | 0.72 | 0.24 | 0.12 | 0.99 | -0.72 | -0.72 |
| **Win Average** | 290 | 0.79 | 0.21 | 0.09 | 1.00 | -1.11 | 0.19 |
| **Loss Average** | 290 | 0.67 | 0.25 | 0.11 | 0.99 | -0.47 | -1.12 |
| **Learners and Completers** | | | | | | | |
| **Time 1** | | | | | | | |
| Win | 118 | 0.78 | 0.22 | 0.09 | 1.00 | -1.13 | 0.13 |
| Loss | 118 | 0.70 | 0.24 | 0.11 | 0.99 | -0.70 | -0.83 |
| Time 1 Average | 236 | 0.74 | 0.23 | 0.09 | 1.00 | -0.90 | -0.44 |
| **Time 2** | | | | | | | |
| Win | 118 | 0.80 | 0.20 | 0.12 | 0.99 | -1.11 | 0.16 |
| Loss | 118 | 0.66 | 0.25 | 0.12 | 0.99 | -0.46 | -1.12 |
| Time 2 Average | 236 | 0.73 | 0.24 | 0.12 | 0.99 | -0.78 | -0.63 |
| **Win Average** | 236 | 0.79 | 0.21 | 0.09 | 1.00 | -1.14 | 0.24 |
| **Loss Average** | 236 | 0.68 | 0.24 | 0.11 | 0.99 | -0.58 | -0.98 |
| **Discount Weight** | | | | | | | |
| **Learners Only** | | | | | | | |
| Time 1 | 145 | 0.39 | 0.31 | 0.00 | 0.98 | 0.32 | -1.41 |
| Time 2 | 145 | 0.33 | 0.29 | 0.01 | 0.93 | 0.67 | -0.96 |
| **Learners and Completers** | | | | | | | |
| Time 1 | 118 | 0.42 | 0.31 | 0.00 | 0.98 | 0.18 | -1.47 |
| Time 2 | 118 | 0.35 | 0.30 | 0.01 | 0.93 | 0.58 | -1.11 |

### Appendix G: Results of reversal learning performance over time

We assessed whether measures of participant performance on our reversal learning task over two testing sessions were correlated. To assess task performance, we quantified both direct (behavioural, estimated using mixed effects regression models), and latent (computational, estimated by fitting reinforcement learning models) measures of task behaviour. We found that participant behaviour across two instances of performing the reversal learning task were significantly positively correlated for behavioural and computational measures of task performance (all *p*s < .001, see ***Supplementary Table 4***). To explore whether measures of task performance varied as a function of time and/or outcome, we used MLMs to model main effects of time, previous outcome (where relevant) and their interactions.

**Supplementary Table 4**

*Correlations of Behavioural and Computational Variables Between Time 1 and Time 2*

| | ***N*** | ***r*** | ***p*** | **95% CI** | |
|---|---|---|---|---|---|
| | | | | **Lower** | **Upper** |
| **Behavioural** | | | | | |
| Reaction Time to Win | 145 | 0.84 | < .001 | 0.78 | 0.88 |
| Reaction Time to Loss | 145 | 0.80 | < .001 | 0.74 | 0.86 |
| Stay After Wins | 145 | 0.66 | <. 001 | 0.56 | 0.74 |
| Stay After Losses | 145 | 0.78 | < .001 | 0.84 | 0.88 |
| Accuracy | 145 | 0.64 | < .001 | 0.53 | 0.73 |
| Accuracy Slopes | 145 | 0.38 | < .001 | 0.23 | 0.51 |
| Perseveration | 145 | 0.54 | < .001 | 0.41 | 0.65 |
| Difference in Reaction Times After Wins and Losses | 145 | 0.73 | < .001 | 0.64 | 0.80 |
| **Reinforcement sensitivity model (computational)** | | | | | |
| Reinforcement Sensitivity for Win | 145 | 0.82 | < .001 | 0.76 | 0.87 |
| Reinforcement Sensitivity for Loss | 145 | 0.85 | < .001 | 0.80 | 0.89 |
| Learning Rate | 145 | 0.77 | < .001 | 0.69 | 0.83 |
| **Softmax model (computational)** | | | | | |
| Inverse Temperature for Win | 145 | 0.67 | < .001 | 0.67 | 0.81 |
| Inverse Temperature for Loss | 145 | 0.53 | < .001 | 0.40 | 0.64 |
| Learning Rate for Win | 145 | 0.31 | < .001 | 0.16 | 0.45 |
| Learning Rate for Loss | 145 | 0.45 | < .001 | 0.31 | 0.57 |
| Discount Weight | 145 | 0.69 | < .001 | 0.60 | 0.77 |

#### Behavioural measures of reversal learning performance

Participant's reaction times were significantly quicker to make a choice following a win on the previous trial than when they experienced a loss (Win: *M* = 515.29, *SD* = 103.13; Loss: *M* = 539.45, *SD* = 112.81; main effect of Outcome: $F(1, 435) = 48.19$, $p < .001$), and reaction times overall were faster at Time 2 than Time 1 (Time 1: $M = 544.71$, $SD = 113.35$; Time 2: $M = 510.03$, $SD = 101.01$; main effect of time: $F(1, 435) = 99.31$, $p < .001$). However, there were no significant interactions between main effects of outcome and time (Time x Outcome: $F(1, 435) = 3.04$, $p = .083$) (**Supplementary Figure *3A***). We also found that the difference in reaction times between wins and losses was significantly lower at Time 2 than at Time 1, where the delta was closer to zero (Time 1: $M = -30.22$, $SD = 37.65$; Time 2: $M = -18.09$, $SD = 32.23$; main effect of Time: $F(1, 145) = 31.10$, $p < .001$) (**Supplementary Figure *3B***).

**Supplementary Figure 3**

*Main Effects of **(A)** Time and Outcome on Reaction Time and **(B)** Time on Difference in Reaction Times After Wins and Losses. Individual datapoints presented here are reaction time and difference in reaction times after wins and losses averaged for each participant per time and outcome. Filled circles denote **(A)** mean of reaction time per time and outcome, and **(B)** difference in reaction times after wins and losses per time. Error bars denote the standard deviation. Reaction times were significantly faster for wins than for losses across both times, and reaction times were significantly faster at Time 2 than Time 1. Difference in reaction times after wins and losses was significantly lower at Time 1 than Time 2.*

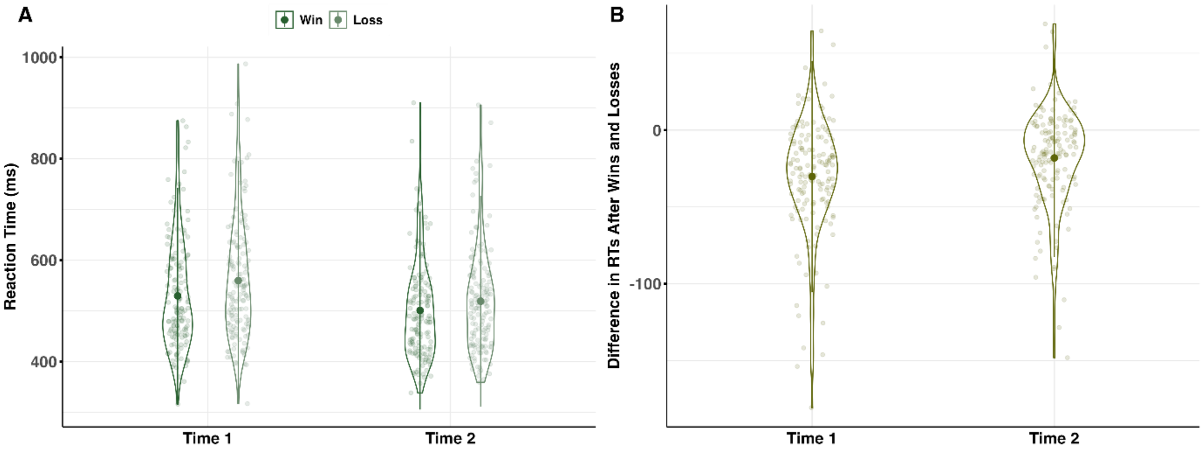


After experiencing a win, participants were significantly more likely to repeat their previous choice on the next trial (stay behaviour) than when they experienced a loss (Win: $M = 0.95$, $SD = 0.06$; Loss: $M = 0.46$, $SD = 0.22$; main effect of Outcome: $F(1, 435) = 2132.20$, $p < .001$), and overall staying behaviour significantly increased between sessions (Time 1: $M = 0.69$ , $SD = 0.29$; Time 2: $M = 0.72$, $SD = 0.28$; main effect of Time: $F(1, 435) = 7.62$, $p = .006$), but no significant interaction was observed between outcome and time (Time x Outcome: $F(1, 435) = 1.03$, $p = .311$) (**Supplementary Figure *4***).

**Supplementary Figure 4**

*Main Effect of Time and Outcome on Staying Behaviour. Individual datapoints presented here are the proportion of stay decisions for each participant per time and outcome. Filled circles denote mean proportion of stay decisions across participants per time and outcome. Error bars denote the standard deviation. Stay behaviour was significantly higher following wins than following losses, and increased significantly from Time 1 to Time 2.*

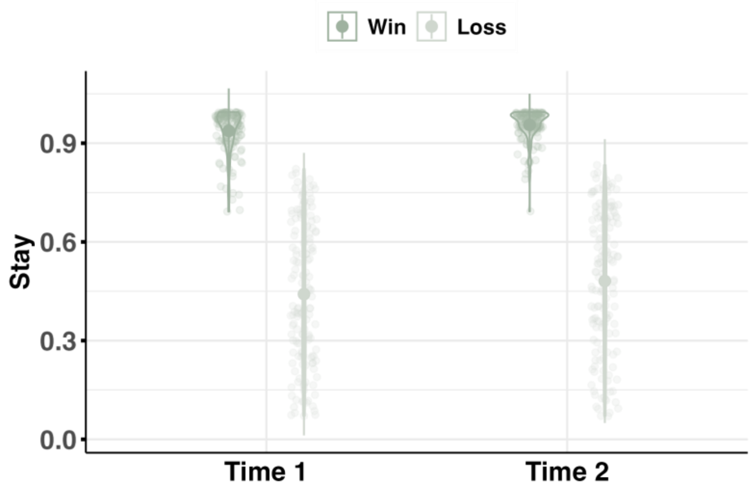


Choice accuracy, regardless of outcome received, significantly increased as a function of time (Time 1: *M* = 0.70, *SD* = 0.05; Time 2: *M* = 0.72, *SD* = 0.05; main effect of Time: $F(1, 145) = 31.78$, $p < .001$) (**Supplementary Figure *5A***), and changes in accuracy across successive reversals (as indexed by accuracy slopes) were significantly higher at Time 2 than Time 1 (Time 1: *M* = -0.03, *SD* = 0.13; Time 2: *M* = 0.04, *SD* = 0.12; main effect of Time: $F(1, 145) = 33.16$, $p < .001$) (**Supplementary Figure *5B***). Additionally, participants made significantly fewer perseverative errors following the reversal of reward contingencies at Time 2 than during Time 1 (Time 1: *M* = 0.13, *SD* = 0.05; Time 2: *M* = 0.11, *SD* = 0.04; main effect of Time: $F(1, 145) = 24.24$, $p < .001$) (**Supplementary Figure *5C***).

**Supplementary Figure 5**

*Main effect of Time on **(A)** Accuracy, **(B)** Accuracy Slope and **(C)** Perseveration. Individual datapoints presented here are accuracy (proportion of choices of the "correct" stimulus), accuracy slope, and perseveration for each participant per time. Filled circles denote mean accuracy, accuracy slope and perseveration across participants per time. Error bars denote the standard deviation. Accuracy significantly increased from Time 1 to Time 2, and changes in accuracy across successive reversals (accuracy slope) were significantly higher at Time 2 than Time 1. Perseveration was significantly lower at Time 2 than Time 1.*

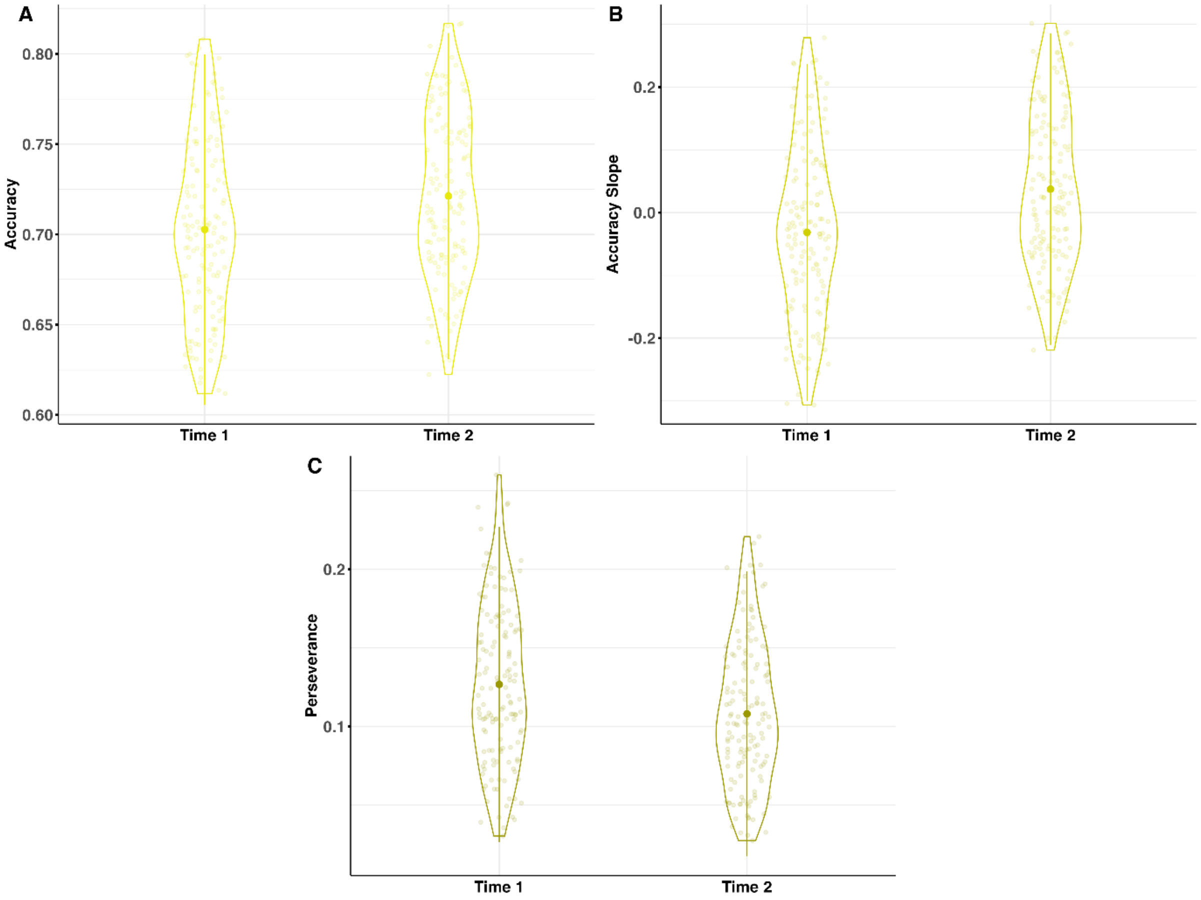


Overall, our behavioural performance measures indicate that participants got better at the reversal learning task as a function of time. Overall reaction time decreased, and the difference in reaction time following wins and losses was significantly different, becoming closer to zero at Time than Time 1. However, this change in reaction time and reduced reaction time delta for outcome did not come at the expense of accuracy. Indeed, participants were more accurate and made fewer perseverative errors at Time 2 than at Time 1.

#### Computational measures of reversal learning performance

The best fitting model overall came from the reinforcement sensitivity family of models. Reinforcement sensitivity was significantly higher for win than loss outcomes (Loss: *M* = -0.72, *SD* = 0.41; Win: *M* = 2.29, *SD* = 1.06; main effect of Outcome: *F*(1, 435) = 2424.14, *p* < .001), and significantly increased from Time 1 to Time 2 (Time 1: *M* = 0.67, *SD* = 1.56; Time 2; *M* = 0.90, *SD* = 1.84; main effect of Time: *F*(1, 435) = 13.32, *p* < .001) (**Error! Reference source not found.**).There was also a significant interaction between outcome and time (Time x Outcome: *F*(1, 435) = 14.81, *p* <.001), such that reinforcement sensitivity varied significantly in response to wins (*p* < .001) but not losses (*p* = .514). Learning rate for the reinforcement sensitivity family did not significantly change across time (Time 1: *M* = 0.71, *SD* = 0.18; Time 2: *M* = 0.71, *SD* = 0.14; main effect of Time: *F*(1, 145) = 0.08, *p* = .772) (**Supplementary Figure 6**).

**Supplementary Figure 6**

*Main Effects of Time and Outcome on Reinforcement Sensitivity. Individual datapoints presented here are reinforcement sensitivity averaged for each participant per time and outcome. Filled circles denote mean of reinforcement sensitivity per time and outcome. Error bars denote the standard deviation. Reinforcement sensitivity was significantly higher for wins than loss outcomes and increased significantly from time 1 to time 2. Reinforcement sensitivity further varied significantly in response to wins but not losses.*

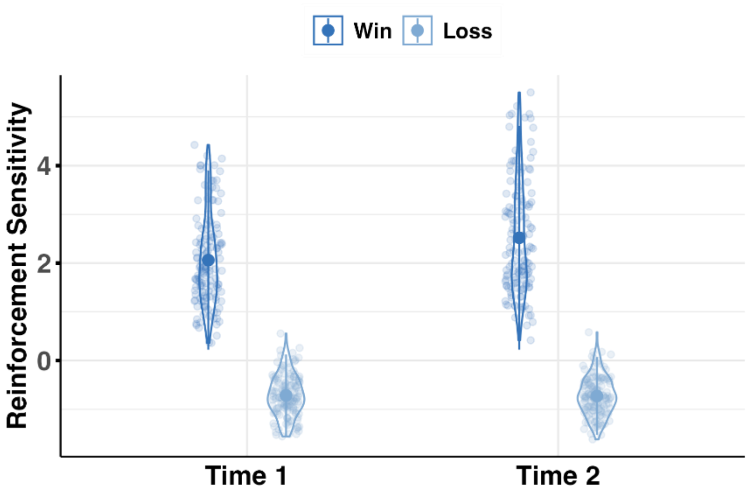


For the best fitting softmax family model, inverse temperature was significantly increased from Time 1 to Time 2 (Time 1: $M$ = 2.65, $SD$ = 2.30; Time 2: $M$ = 3.23, $SD$ = 2.66; main effect of Time $F(1, 435) = 15.61$, $p < .001$), but no main effect of previous outcome nor interaction was observed (main effect of Outcome: $F(1, 435) = 1.44$, $p = .231$; Time x Outcome: $F(1, 435) = 0.46$, $p = .500$) (**Supplementary Figure *7A***). Similarly to the learning rate for the best fitting reinforcement sensitivity model, learning rate did not vary as a function of time (main effect of Time: $F(1, 435) = 0.24$, $p = .623$). However learning rate was significantly higher for wins ($M$ = 0.79, $SD$ = 0.21) than for losses ($M$ = 0.67, $SD$ = 0.25) across both times (main effect of Outcome: $F(1, 435) = 54.97$, $p < .001$), and there was a significant interaction (Time x Outcome: $F(1, 435) = 6.61$, $p = .010$), such that the effect of time on learning rate decreased significantly for losses ($p = .025$) but not wins ($p = .113$) (**Supplementary Figure *7B***). Lastly, the rate of expected value updating for the unchosen option varied as a function of time, such that the discount weight was significantly higher in Time 1 than Time 2 (Time 1: $M$ = 0.39, $SD$ = 0.31; Time 2: $M$ = 0.33, $SD$ = 0.29; main effect of Time: $F(1, 145) = 8.59$, $p = .004$) (**Supplementary Figure *7C***).

**Supplementary Figure 7**

*Main Effects of **(A)** Time and Outcome on Inverse Temperature, **(B)** Time and Outcome on Learning Rate and **(C)** Time on Discount Weight. Individual datapoints presented here are inverse temperature, learning rate, and discount weight averaged for each participant per time (and outcome). Filled circles denote mean of inverse temperature, learning rate, and discount weight per time (and outcome). Error bars denote the standard deviation.*

*Inverse temperature significantly increased from Time 1 to Time 2, but did not vary depending on outcome. Learning rate did not change across time but was significantly higher for wins than for losses across both times, and the effect of time on learning rate decreased significantly for losses but not for wins. Discount weight was significantly higher at Time 1 than at Time 2.*

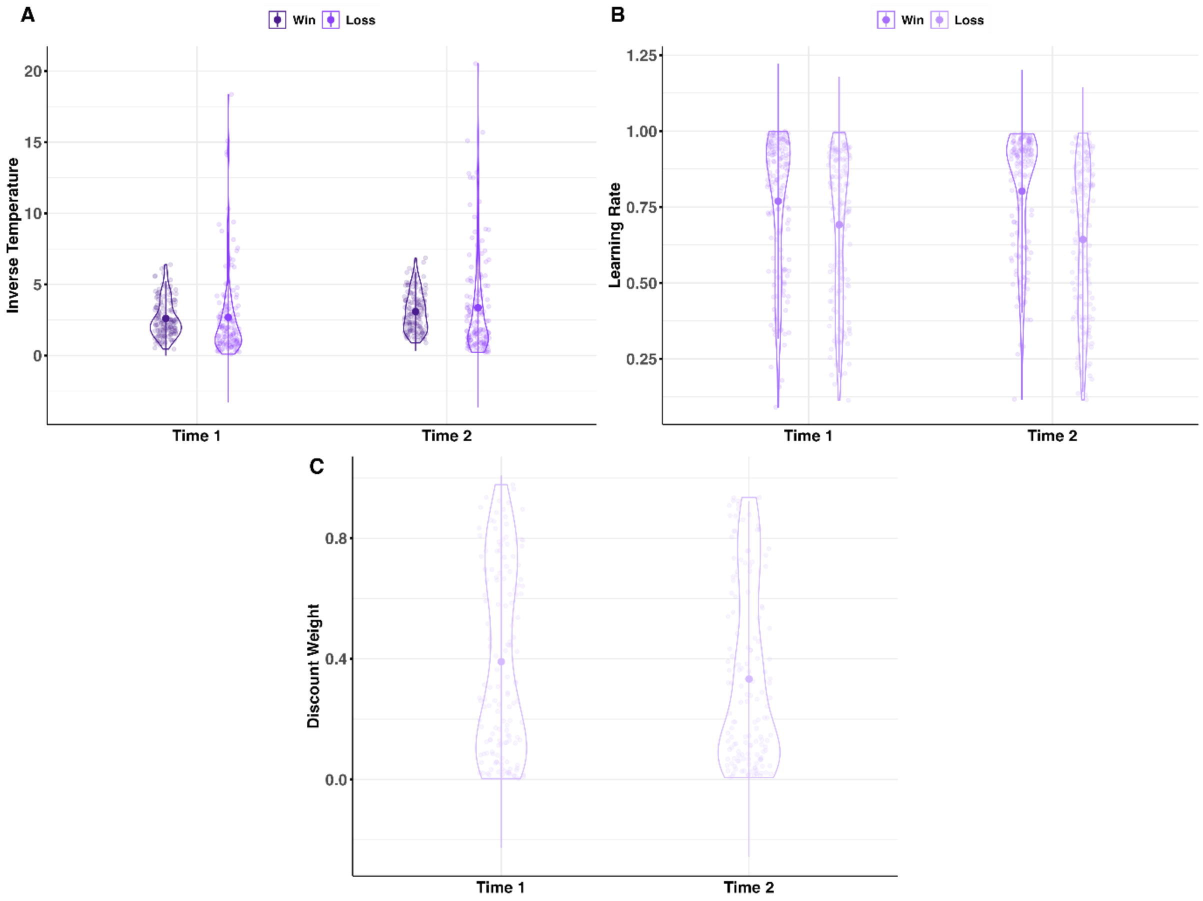


Overall, our computational performance measures also indicate that task performance improved across the two sessions. Reinforcement sensitivity, which determines the maximum difference in expected value of the "better" and "worse" choice, increased over time, meaning choices could be more easily discriminated between. However, the rate at which expected values did not change varied as a function of time. For the best fitting softmax model, inverse temperatures (extent to which choice is driven by expected value) increased from Time 1 to Time 2 but did not vary as a function of outcome, suggesting that expected value was more important for choice behaviour in the second session regardless of the outcome received. Learning rate also did not vary as a function of time for the softmax model, but was significantly higher for wins than losses and subject to an interaction between outcome and time, meaning wins were more informative for updating expected values than losses, and the impact of losses for expected value updating was significantly lower in the second session. The rate of expected value updating for the unchosen option also varied as a function of time, with the extent of counterfactual value updating decreasing as a function of time.

### Appendix H: Full Results of IU-Related Main Analyses

#### Behavioural Variables

##### Reaction Time

###### *Intolerance of Uncertainty Scale (IUS)*

A significant main effect of IUS on reaction times was observed when IUS was entered into the model alone [$p$ = .020, see ***Supplementary Table 5***]. However, the effect of IUS was no longer significant when entered into the model together with STICSA [IUS: $F(1, 119) = 3.36$, $p$ = .069; STICSA: $F(1, 119) = 0.11$, $p$ = .743]. Individual differences in IUS were not significantly associated with effects of Time or Outcome on reaction times (see ***Supplementary Table 5***, min $p$ = .057).

###### *Prospective Intolerance of Uncertainty (P-IU)*

There were no significant relationships observed between P-IU and reaction times (see ***Supplementary Table 5***, min $p$ = .074).

###### *Inhibitory Intolerance of Uncertainty (I-IU)*

The analyses revealed a significant main effect of I-IU on reaction times when entered into the model alone [$p$ = .008, see Table 8], however, the effect of I-IU did not hold after controlling for STICSA [I-IU: $F(1, 119) = 3.77$, $p$ = .055; STICSA: $F(1, 119) = 0.13$, $p$ = .718] or for P-IU [I-IU: $F(1, 145) = 2.67$, $p$ = .104; P-IU: $F(1, 145) = 0.04$, $p$ = .839]. I-IU was not significantly associated with differential reaction times across Time or Outcome (see ***Supplementary Table 5***, min $p$ = .240).

**Supplementary Table 5**

*Results from MLMs Assessing Intolerance of Uncertainty, Prospective Intolerance of Uncertainty, and Inhibitory Intolerance of Uncertainty Main Effects and Interactions with Time (and Outcome) on Reaction Time*

| **Reaction Time** | **df** | ***F*** | ***p*** |
|---|---|---|---|
| **IUS x Time x Outcome** | | | |
| IUS | 1, 145 | 5.53 | .020 |
| IUS x Time | 1, 435 | 3.65 | .057 |
| IUS x Outcome | 1, 435 | 0.80 | .372 |
| IUS x Time x Outcome | 1, 435 | 0.16 | .690 |
| **P-IU x Time x Outcome** | | | |
| P-IU | 1, 145 | 3.24 | .074 |
| P-IU x Time | 1, 435 | 1.97 | .162 |
| P-IU x Outcome | 1, 435 | 0.37 | .543 |
| P-IU x Time x Outcome | 1, 435 | 0.25 | .618 |
| **I-IU x Time x Outcome** | | | |
| I-IU | 1, 145 | 7.31 | .008 |
| I-IU x Time | 1, 435 | 5.13 | .240 |

| Reaction Time | | | |
|---|---|---|---|
| | **df** | ***F*** | ***p*** |
| I-IU x Outcome | 1, 435 | 1.23 | .268 |
| I-IU x Time x Outcome | 1, 435 | 0.06 | .807 |

*Note*. Entries in the table that are formatted in bold indicate $p < .05$ and that the effect was significant when controlling for STICSA (or, where relevant, the alternate IU subscale). Black font indicates $p < .05$ and that the effect was not significant when controlling for STICSA (or, where relevant, the alternate IU subscale). Gray font denotes $p > .05$.

#### Stay

##### *Intolerance of Uncertainty Scale (IUS)*

There were no significant relationships observed between IUS and stay behaviour (see ***Supplementary Table 6***, min $p = .184$).

##### *Prospective Intolerance of Uncertainty (P-IU)*

There analyses did not reveal any significant relationships between P-IU and stay behaviour (see ***Supplementary Table 6***, min $p = .239$).

##### *Inhibitory Intolerance of Uncertainty (I-IU)*

There were no significant relationships observed between I-IU and stay behaviour (see ***Supplementary Table 6***, min $p = .177$).

**Supplementary Table 6**

*Results from MLMs Assessing Intolerance of Uncertainty, Prospective Intolerance of Uncertainty, and Inhibitory Intolerance of Uncertainty Main Effects and Interactions with Time (and Outcome) on Stay*

| Stay | | | |
|---|---|---|---|
| | **df** | ***F*** | ***p*** |
| **IUS x Time x Outcome** | | | |
| IUS | 1, 145 | 1.78 | .184 |
| IUS x Time | 1, 435 | 0.16 | .691 |
| IUS x Outcome | 1, 435 | 0.20 | .654 |
| IUS x Time x Outcome | 1, 435 | 0.31 | .577 |
| **P-IU x Time x Outcome** | | | |
| P-IU | 1, 145 | 1.40 | .239 |
| P-IU x Time | 1, 435 | 0.16 | .692 |
| P-IU x Outcome | 1, 435 | 0.26 | .608 |
| P-IU x Time x Outcome | 1.435 | 0.43 | .511 |
| **I-IU x Time x Outcome** | | | |
| I-IU | 1, 145 | 1.84 | .177 |
| I-IU x Time | 1, 435 | 0.13 | .723 |
| I-IU x Outcome | 1, 435 | 0.11 | .743 |
| I-IU x Time x Outcome | 1, 435 | 0.15 | .697 |

*Note*. Entries in the table that are formatted in bold indicate $p < .05$ and that the effect was significant when controlling for STICSA (or, where relevant, the alternate IU subscale). Black font indicates $p < .05$ and that the effect was not significant when controlling for STICSA (or, where relevant, the alternate IU subscale). Gray font denotes $p > .05$.

#### Accuracy

##### *Intolerance of Uncertainty Scale (IUS)*

There were no significant relationships observed between IUS and accuracy (see ***Supplementary Table 7***, min $p = .104$).

##### *Prospective Intolerance of Uncertainty (P-IU)*

There were no significant relationships observed between P-IU and accuracy (see ***Supplementary Table 7***, min $p = .310$).

##### *Inhibitory Intolerance of Uncertainty (I-IU)*

The analyses revealed a significant main effect of I-IU on overall accuracy when I-IU was entered into the model alone [$p = .033$, see ***Supplementary Table 7***] as well as when entered with STICSA [I-IU: $F(1, 119) = 5.93$, $p = .016$; STICSA: $F(1, 119) = 0.60$, $p = .441$] and P-IU [I-IU: $F(1, 145) = 4.87$, $p = .029$; P-IU: $F(1, 145) = 1.29$, $p = .258$]. A follow-up partial correlational test controlling for STICSA showed that higher I-IU was significantly associated with higher accuracy [$r(119) = 0.22$, $p = .018$] (see ***Supplementary Figure 8***). This relationship further held when controlling for P-IU [$r(145) = 0.18$, $p = .031$]. Individual differences in I-IU were not related to variations in accuracy across time [$p = .608$, see ***Supplementary Table 7***].

**Supplementary Figure 8**

*Scatterplot With Histograms Depicting the Partial Correlation Between Inhibitory Intolerance of Uncertainty (Controlling for STICSA) and Accuracy.*

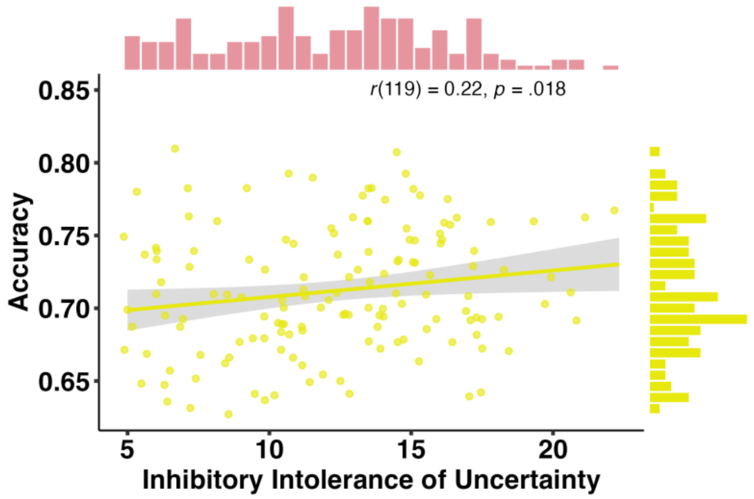

*Note*. The distribution of Inhibitory IU (I-IU) scores is displayed on top of the figure in pink, and the distribution of accuracy is displayed on the right side of the figure in yellow. Shaded areas represent 95% confidence intervals. Higher I-IU was associated with higher levels of accuracy. This relationship further held when controlling for P-IU [$r(145) = 0.18$, $p = .031$].

**Supplementary Table 7**

*Results from MLMs Assessing Intolerance of Uncertainty, Prospective Intolerance of Uncertainty, and Inhibitory Intolerance of Uncertainty Main Effects and Interactions with Time (and Outcome) on Accuracy*

| **Accuracy** | **df** | ***F*** | ***p*** |
|---|---|---|---|
| **IUS x Time** | | | |
| IUS | 1, 145 | 2.68 | .104 |
| IUS x Time | 1, 145 | 0.11 | .737 |
| **P-IU x Time** | | | |
| P-IU | 1, 145 | 1.04 | .310 |
| P-IU x Time | 1, 145 | 0.02 | .887 |
| **I-IU x Time** | | | |
| I-IU | **1, 145** | **4.61** | **.033** |
| I-IU x Time | 1, 145 | 0.26 | .608 |

*Note*. Entries in the table that are formatted in bold indicate $p < .05$ and that the effect was significant when controlling for STICSA (or, where relevant, the alternate IU subscale). Black font indicates $p < .05$ and that the effect was not significant when controlling for STICSA (or, where relevant, the alternate IU subscale). Gray font denotes $p > .05$.

#### Accuracy Slope

##### *Intolerance of Uncertainty Scale (IUS)*

There analyses did not reveal any significant relationships between IUS and accuracy slopes (see ***Supplementary Table 8***, min $p = .170$).

##### *Prospective Intolerance of Uncertainty (P-IU)*

There were no significant relationships observed between P-IU and accuracy slope (see ***Supplementary Table 8***, min $p = .400$).

##### *Inhibitory Intolerance of Uncertainty (I-IU)*

We did not observe any significant relationships between I-IU and accuracy slopes (see ***Supplementary Table 8***, min $p = .070$).

**Supplementary Table 8**

*Results from MLMs Assessing Intolerance of Uncertainty, Prospective Intolerance of Uncertainty, and Inhibitory Intolerance of Uncertainty Main Effects and Interactions with Time (and Outcome) on Accuracy Slope*

| **Accuracy slope** | **df** | ***F*** | ***p*** |
|---|---|---|---|
| **IUS x Time** | | | |
| IUS | 1, 145 | 1.91 | .170 |
| IUS x Time | 1, 145 | 0.29 | .590 |
| **P-IU x Time** | | | |
| P-IU | 1, 145 | 0.71 | .400 |
| P-IU x Time | 1, 145 | 0.09 | .767 |
| **I-IU x Time** | | | |
| I-IU | 1, 145 | 3.33 | .070 |
| I-IU x Time | 1, 145 | 0.56 | .455 |

*Note*. Entries in the table that are formatted in bold indicate $p < .05$ and that the effect was significant when controlling for STICSA (or, where relevant, the alternate IU subscale). Black font indicates $p < .05$ and that the effect was not significant when controlling for STICSA (or, where relevant, the alternate IU subscale). Gray font denotes $p > .05$.

#### Perseveration

##### *Intolerance of Uncertainty Scale (IUS)*

There were no significant relationships observed between IUS and perseveration (see ***Supplementary Table 9***, min $p = .114$).

##### *Prospective Intolerance of Uncertainty (P-IU)*

There analyses did not reveal any significant relationships between P-IU and perseveration (see ***Supplementary Table 9***, min $p = .344$).

##### *Inhibitory Intolerance of Uncertainty (I-IU)*

There was a significant main effect of I-IU on overall perseveration when I-IU was entered into the model alone [$p = .035$, see ***Supplementary Table 9***] as well as when entered with STICSA [I-IU: $F(1, 119) = 5.19$, $p = .0024$; STICSA: $F(1, 119) = 0.509$, $p = .477$] and P-IU [I-IU: $F(1, 145) = 5.14$, $p = .025$; P-IU: $F(1, 145) = 1.50$, $p = .222$]. A follow-up partial correlational test controlling for STICSA showed that higher levels of I-IU were significantly associated with lower perseveration [$r(119) = -0.20$, $p = .026$] (see ***Supplementary Figure 9***). This relationship further held when controlling for P-IU [$r(145) = -0.19$, $p = .026$]. Individual differences in I-IU did not differentially affect perseveration across time (see ***Supplementary Table 9***).

##### Supplementary Figure 9

*Scatterplot With Histograms Depicting the Partial Correlation Between Inhibitory Intolerance of Uncertainty (Controlling for STICSA) and Perseveration.*

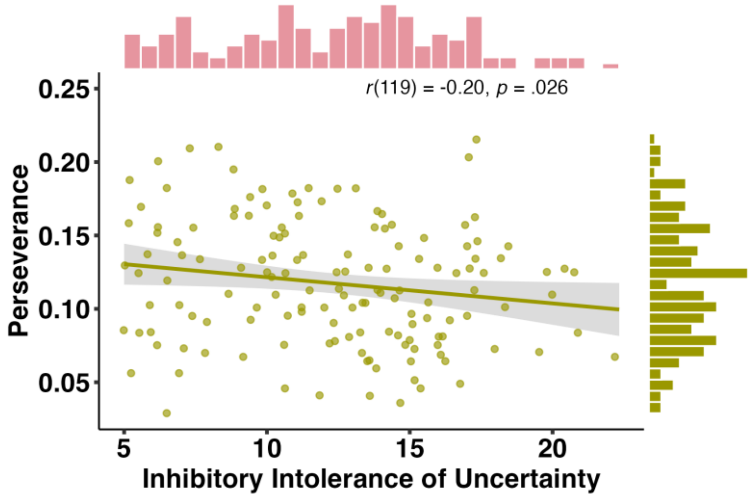


*Note*. The distribution of Inhibitory IU (I-IU) scores is displayed on top of the figure in pink, and the distribution of perseveration is displayed on the right side of the figure in green. Shaded areas represent 95% confidence intervals. Higher I-IU was associated with lower perseveration. This relationship further held when controlling for P-IU [$r(145) = -0.19$, $p = .026$].

##### Supplementary Table 9

*Results from MLMs Assessing Intolerance of Uncertainty, Prospective Intolerance of Uncertainty, and Inhibitory Intolerance of Uncertainty Main Effects and Interactions with Time (and Outcome) on Perseveration*

| **Perseveration** | **df** | ***F*** | ***p*** |
|---|---|---|---|
| **IUS x Time** | | | |
| IUS | 1, 145 | 2.52 | .114 |
| IUS x Time | 1, 145 | 0.18 | .675 |
| **P-IU x Time** | | | |
| P-IU | 1, 145 | 0.90 | .344 |
| P-IU x Time | 1, 145 | 0.08 | .788 |
| **I-IU x Time** | | | |
| I-IU | **1, 145** | **4.52** | **.035** |
| I-IU x Time | 1, 145 | 0.29 | .590 |

*Note*. Entries in the table that are formatted in bold indicate $p < .05$ and that the effect was significant when controlling for STICSA (or, where relevant, the alternate IU subscale). Black font indicates $p < .05$ and that the effect was not significant when controlling for STICSA (or, where relevant, the alternate IU subscale). Gray font denotes $p > .05$.

#### Difference in Reaction Times After Wins And Losses

##### *Intolerance of Uncertainty Scale (IUS)*

There were no significant relationships observed between IUS and differences in reaction times after wins and losses (see ***Supplementary Table 10***, min $p$ = .203).

##### *Prospective Intolerance of Uncertainty (P-IU)*

We did not observe any significant relationships between P-IU and differences in reaction times after wins and losses (see ***Supplementary Table 10***, min $p$ = .110).

##### *Inhibitory Intolerance of Uncertainty (I-IU)*

Differences in reaction times after wins and losses were not significantly affected by individual differences in I-IU (see ***Supplementary Table 10***, min $p$ = .153).

**Supplementary Table 10**

*Results from MLMs Assessing Intolerance of Uncertainty, Prospective Intolerance of Uncertainty, and Inhibitory Intolerance of Uncertainty Main Effects and Interactions with Time (and Outcome) on Difference in Reaction Times After Wins and Losses*

| **Difference in reaction times after wins and losses** | **df** | ***F*** | ***p*** |
|---|---|---|---|
| **IUS x Time** | | | |
| IUS | 1, 145 | 1.34 | .250 |
| IUS x Time | 1, 145 | 1.63 | .203 |
| **P-IU x Time** | | | |
| P-IU | 1, 145 | 0.62 | .432 |
| P-IU x Time | 1, 145 | 2.59 | .110 |
| **I-IU x Time** | | | |
| I-IU | 1, 145 | 2.06 | .153 |
| I-IU x Time | 1, 145 | 0.61 | .438 |

*Note*. Entries in the table that are formatted in bold indicate $p$ < .05 and that the effect was significant when controlling for STICSA (or, where relevant, the alternate IU subscale). Black font indicates $p$ < .05 and that the effect was not significant when controlling for STICSA (or, where relevant, the alternate IU subscale). Gray font denotes $p$ > .05.

#### Reinforcement Sensitivity Model Variables

#### Reinforcement sensitivity

##### *Intolerance of Uncertainty Scale (IUS)*

There was a significant interaction between IUS and Outcome when IUS was entered into the model alone [$p$ = .001] as well as after controlling for STICSA [IUS x Outcome: $F(1, 357)$ = 6.53, $p$ = .011; STICSA x Outcome: $F(1, 357)$ = 0.28, $p$ = .599]. A follow-up partial correlational test controlling for STICSA showed that higher IUS was significantly associated with higher reinforcement sensitivity to wins [*r*(119) = 0.18, *p* = .049] (see ***Supplementary Figure 10***). IUS was not significantly related to reinforcement sensitivity for loss [*r*(119) = 0.02, *p* = .838]. Individual differences in IUS did not differentially affect reinforcement sensitivity across time [*p* = .406, see ***Supplementary Table 11***].

**Supplementary Figure 10**

*Scatterplot With Histograms Depicting the Partial Correlation Between Intolerance of Uncertainty (Controlling for STICSA) and Reinforcement Sensitivity for Win.*

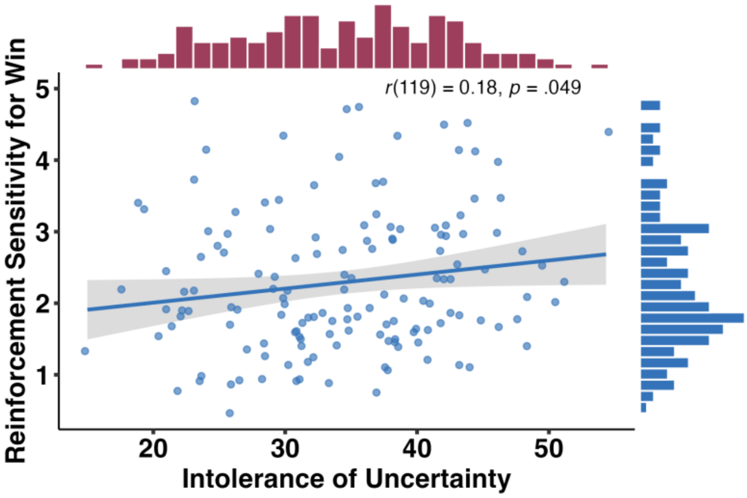


*Note*. The distribution of intolerance of uncertainty scores (IUS) is displayed on top of the figure in pink, and the distribution of reinforcement sensitivity for win is displayed on the right side of the figure in blue. Shaded areas represent 95% confidence intervals. Higher IUS was associated with higher reinforcement sensitivity for wins.

##### *Prospective Intolerance of Uncertainty (P-IU)*

As with overall IUS, individual differences in P-IU were not significantly associated with overall reinforcement sensitivity [*p* = .157, see ***Supplementary Table 11***]. However, individual differences in P-IU were significantly associated with varying reinforcement sensitivity in response to wins vs. losses when entered into the model alone [*p* = .021] and when controlling for STICSA [P-IU x Outcome: *F*(1, 357) = 4.83, *p* = .029; STICSA x Outcome: *F*(1, 357) = 1.00, *p* = .317] but not when controlling for I-IU [P-IU x Outcome: *F*(1, 435) = 1.71, *p* = .192; I-IU x Outcome: *F*(1, 435) = 11.67, *p* = .001]. P-IU was not related to differential reinforcement sensitivity across time [*p* = .475, ***Supplementary Table 11***].

##### *Inhibitory Intolerance of Uncertainty (I-IU)*

Similarly to IUS and P-IU, overall I-IU did not significantly affect reinforcement sensitivity [*p* = .095, see ***Supplementary Table 11***]. However, there was a significant interaction between I-IU and Outcome when I-IU was entered into the model alone [*p* = .001] as well as after controlling for STICSA [I-IU x Outcome: *F*(1, 357) = 6.59, *p* = .011; STICSA x Outcome: *F*(1, 357) = 0.21, *p* = .535] and P-IU [I-IU x Outcome: *F*(1, 435) = 11.67, *p* = .001; P-IU x Outcome: *F*(1, 435) = 1.71, *p* = .192]. Whilst higher I-IU was associated with higher reinforcement sensitivity for win and lower reinforcement sensitivity for loss, these follow-up partial correlations controlling for STICSA were non-significant [Reinforcement sensitivity for win: $r(119) = 0.17$, $p = .072$; Reinforcement sensitivity for loss: $r(119) = -0.02$, $p = .840$]. Individual differences in I-IU did not differentially affect reinforcement sensitivity across time [$p = .385$, see ***Supplementary Table 11***].

##### Supplementary Table 11

*Results from MLMs Assessing Intolerance of Uncertainty, Prospective Intolerance of Uncertainty, and Inhibitory Intolerance of Uncertainty Main Effects and Interactions with Time (and Outcome) on Reinforcement Sensitivity*

| **Reinforcement sensitivity** | **df** | ***F*** | ***p*** |
|---|---|---|---|
| **IUS x Time x Outcome** | | | |
| IUS | 1, 145 | 2.66 | .105 |
| IUS x Time | 1, 435 | 0.69 | .406 |
| IUS x Outcome | **1, 435** | **10.58** | **.001** |
| IUS x Time x Outcome | 1, 435 | 1.28 | .258 |
| **P-IU x Time x Outcome** | | | |
| P-IU | 1, 145 | 2.02 | .157 |
| P-IU x Time | 1, 435 | 0.51 | .475 |
| P-IU x Outcome | 1, 435 | 5.39 | .021 |
| P-IU x Time x Outcome | 1.435 | 1.02 | .314 |
| **I-IU x Time x Outcome** | | | |
| I-IU | 1, 145 | 2.82 | .095 |
| I-IU x Time | 1, 435 | 0.76 | .385 |
| I-IU x Outcome | **1, 435** | **15.44** | **< .001** |
| I-IU x Time x Outcome | 1, 435 | 1.31 | .254 |

*Note*. Entries in the table that are formatted in bold indicate $p < .05$ and that the effect was significant when controlling for STICSA (or, where relevant, the alternate IU subscale). Black font indicates $p < .05$ and that the effect was not significant when controlling for STICSA (or, where relevant, the alternate IU subscale). Gray font denotes $p > .05$.

#### Learning rate

##### *Intolerance of Uncertainty Scale (IUS)*

There were no significant relationships observed between IUS and learning rate (see ***Supplementary Table 12***, min $p = .080$).

##### *Prospective Intolerance of Uncertainty (P-IU)*

As with overall IUS, individual differences in P-IU were not significantly associated with overall learning rate [$p = .733$, see ***Supplementary Table 12***]. Individual differences in P-IU were significantly associated with varying learning rate at Time 1 vs. Time 2 when entered into the model alone [$p = .036$]. However, this interaction did not hold when controlling for STICSA [P-IU x Time: $F(1, 119) = 2.56$, $p = .112$; STICSA x Time: $F(1, 119) =$

1.46, *p* = .229] or when controlling for I-IU [P-IU x Time: *F*(1, 145) = 3.74, *p* = .055; I-IU x Time: *F*(1, 145) = 0.67, *p* = .414].

###### *Inhibitory Intolerance of Uncertainty (I-IU)*

There were no significant relationships observed between I-IU and learning rate (see ***Supplementary Table 12***, min *p* = .238).

**Supplementary Table 12**

*Results from MLMs Assessing Intolerance of Uncertainty, Prospective Intolerance of Uncertainty, and Inhibitory Intolerance of Uncertainty Main Effects and Interactions with Time (and Outcome) on Learning Rate from the Reinforcement Sensitivity Computational Model*

| **Learning Rate** | **df** | ***F*** | ***p*** |
|---|---|---|---|
| **IUS x Time** | | | |
| IUS | 1, 145 | 0.08 | .775 |
| IUS x Time | 1, 145 | 3.11 | .080 |
| **P-IU x Time** | | | |
| P-IU | 1, 145 | 0.12 | .733 |
| P-IU x Time | 1, 145 | 4.48 | .036 |
| **I-IU x Time** | | | |
| I-IU | 1, 145 | 0.90 | .344 |
| I-IU x Time | 1, 145 | 1.40 | .238 |

*Note*. Entries in the table that are formatted in bold indicate *p* < .05 and that the effect was significant when controlling for STICSA (or, where relevant, the alternate IU subscale). Black font indicates *p* < .05 and that the effect was not significant when controlling for STICSA (or, where relevant, the alternate IU subscale). Gray font denotes *p* > .05.

#### Softmax Model Variables

##### Inverse temperature

###### *Intolerance of Uncertainty Scale (IUS)*

A significant interaction between IUS and Outcome was observed when IUS was entered into the model alone [*p* = .011, see ***Supplementary Table 13***]. However, the IUS x Outcome interaction was no longer significant when entered into the model together with STICSA [IUS x Outcome: *F*(1, 357) = 2.16, *p* = .143; STICSA x Outcome: *F*(1, 357) = 1.19, *p* = .276]. Individual differences in IUS were not significantly associated overall or with effects of Time on inverse temperature (see ***Supplementary Table 13***, min *p* = .051).

###### *Prospective Intolerance of Uncertainty (P-IU)*

Individual differences in P-IU were not significantly associated with overall inverse temperature [*p* = .129, see ***Supplementary Table 13***]. P-IU was significantly associated with varying inverse temperature in response to wins vs. losses when entered into the model alone [*p* = .025] but not after controlling for STICSA [P-IU x Outcome: *F*(1, 357) = 3.30, *p* = .070; STICSA x Outcome: *F*(1, 357) = 1.26, *p* = .262] or for I-IU [P-IU x Outcome: *F*(1, 435) = 0.05, *p* = .830; I-IU x Outcome: *F*(1, 435) = 2.05, *p* = .153]. However, individual differences in P-IU were significantly associated with differential inverse temperature at Time 1 vs. Time 2 both when entered into the model alone [*p* = .021, see ***Supplementary Table 13***] as well as when controlling for STICSA [P-IU x Time: *F*(1, 357) = 5.45, *p* = .020; STICSA x Time: *F*(1, 357) = 0.06, *p* = .808] and for I-IU [P-IU x Time: *F*(1, 435) = 4.25, *p* = .040; I-IU x Time: *F*(1, 435) = 0.70, *p* = .405]. Higher levels of P-IU were associated with higher inverse temperature both at Time 1 and Time 2, but neither of these relationships were statistically significant [Time 1: *r*(119) = 0.14, *p* = .125; Time 2: *r*(119) = 0.14, *p* = .125].

##### *Inhibitory Intolerance of Uncertainty (I-IU)*

The analyses revealed a significant main effect of I-IU as well as an interaction between I-IU and Outcome on inverse temperature when I-IU was entered into the model alone [I-IU: *p* = .027; I-IU x Outcome: *p* = .009, see ***Supplementary Table 13***]. However, neither the main effect of I-IU or the I-IU x Outcome interaction remained significant after controlling for STICSA [I-IU: *F*(1, 119) = 1.53, *p* = .219; STICSA: *F*(1, 119) = 0.24, *p* = .623; I-IU x Outcome: *F*(1, 357) = 0.62, *p* = .431; STICSA x Outcome: *F*(1, 357) = 2.12, *p* = .147] or for P-IU [I-IU: *F*(1, 145) = 2.81, *p* = .096; P-IU: *F*(1, 145) = 0.17, *p* = .684; I-IU x Outcome: *F*(1, 435) = 2.05, *p* = .153; P-IU x Outcome: *F*(1, 435) = 0.05, *p* = .830]. Individual differences in I-IU were further not significantly associated with differential inverse temperature at Time 1 vs. Time 2 (see ***Supplementary Table 13***, min *p* = .182).

##### Supplementary Table 13

*Results from MLMs Assessing Intolerance of Uncertainty, Prospective Intolerance of Uncertainty, and Inhibitory Intolerance of Uncertainty Main Effects and Interactions with Time (and Outcome) on Inverse Temperature*

| **Inverse Temperature** | **df** | ***F*** | ***p*** |
|---|---|---|---|
| **IUS x Time x Outcome** | | | |
| IUS | 1, 145 | 3.87 | .051 |
| IUS x Time | 1, 435 | 3.79 | .052 |
| IUS x Outcome | 1, 435 | 6.59 | .011 |
| IUS x Time x Outcome | 1, 435 | 0.99 | .320 |
| **P-IU x Time x Outcome** | | | |
| P-IU | 1, 145 | 2.33 | .129 |
| P-IU x Time | **1, 435** | **5.33** | **.021** |
| P-IU x Outcome | 1, 435 | 5.03 | .025 |
| P-IU x Time x Outcome | 1, 435 | 1.74 | .188 |
| **I-IU x Time x Outcome** | | | |
| I-IU | 1, 145 | 5.01 | .027 |
| I-IU x Time | 1, 435 | 1.78 | .182 |
| I-IU x Outcome | 1, 435 | 6.97 | .009 |

| **Inverse Temperature** | | | |
| --- | --- | --- | --- |
| | **df** | ***F*** | ***p*** |
| I-IU x Time x Outcome | 1, 435 | 0.28 | .596 |

*Note*. Entries in the table that are formatted in bold indicate *p* < .05 and that the effect was significant when controlling for STICSA (or, where relevant, the alternate IU subscale). Black font indicates *p* < .05 and that the effect was not significant when controlling for STICSA (or, where relevant, the alternate IU subscale). Gray font denotes *p* > .05.

#### Learning Rate

##### *Intolerance of Uncertainty Scale (IUS)*

There were no significant relationships observed between IUS and learning rate (see ***Supplementary Table 14***, min *p* = .394).

##### *Prospective Intolerance of Uncertainty (P-IU)*

Learning rate was not significantly affected by individual differences in P-IU (see ***Supplementary Table 14***, min *p* = .170).

##### *Inhibitory Intolerance of Uncertainty (I-IU)*

We did not observe any significant relationships between I-IU and learning rate (see ***Supplementary Table 14***, min *p* = .337).

**Supplementary Table 14**

*Results from MLMs Assessing Intolerance of Uncertainty, Prospective Intolerance of Uncertainty, and Inhibitory Intolerance of Uncertainty Main Effects and Interactions with Time (and Outcome) on Learning Rate from the Softmax Computational Model*

| **Learning rate** | | | |
| --- | --- | --- | --- |
| | **df** | ***F*** | ***p*** |
| **IUS x Time x Outcome** | | | |
| IUS | 1, 145 | 0.38 | .536 |
| IUS x Time | 1, 435 | 1.45 | .229 |
| IUS x Outcome | 1, 435 | 0.61 | .434 |
| IUS x Time x Outcome | 1, 435 | 0.73 | .394 |
| **P-IU x Time x Outcome** | | | |
| P-IU | 1, 145 | 1.02 | .314 |
| P-IU x Time | 1, 435 | 1.89 | .170 |
| P-IU x Outcome | 1, 435 | 0.29 | .588 |
| P-IU x Time x Outcome | 1, 435 | 0.51 | .476 |
| **I-IU x Time x Outcome** | | | |
| I-IU | 1, 145 | 0.02 | .902 |
| I-IU x Time | 1, 435 | 0.79 | .375 |
| I-IU x Outcome | 1, 435 | 0.92 | .337 |
| I-IU x Time x Outcome | 1, 435 | 0.83 | .362 |

*Note*. Entries in the table that are formatted in bold indicate $p < .05$ and that the effect was significant when controlling for STICSA (or, where relevant, the alternate IU subscale). Black font indicates $p < .05$ and that the effect was not significant when controlling for STICSA (or, where relevant, the alternate IU subscale). Gray font denotes $p > .05$.

#### Discount weight

##### *Intolerance of Uncertainty Scale (IUS)*

Discount weight was not significantly affected by individual differences in IUS (see ***Supplementary Table 15***, min $p$ = .452).

##### *Prospective Intolerance of Uncertainty (P-IU)*

Discount weight was not significantly affected by individual differences in P-IU (see ***Supplementary Table 15***, min $p$ = .234).

##### *Inhibitory Intolerance of Uncertainty (I-IU)*

Discount weight was not significantly affected by individual differences in I-IU (see ***Supplementary Table 15***, min $p$ = .336).

**Supplementary Table 15**

*Results from MLMs Assessing Intolerance of Uncertainty, Prospective Intolerance of Uncertainty, and Inhibitory Intolerance of Uncertainty Main Effects and Interactions with Time (and Outcome) on Discount Weight*

| **Discount weight** | **df** | ***F*** | ***p*** |
|---|---|---|---|
| **IUS x Time** | | | |
| IUS | 1, 145 | 0.24 | .623 |
| IUS x Time | 1, 145 | 0.57 | .452 |
| **P-IU x Time** | | | |
| P-IU | 1, 145 | 0.00 | .986 |
| P-IU x Time | 1, 145 | 1.43 | .234 |
| **I-IU x Time** | | | |
| I-IU | 1, 145 | 0.93 | .336 |
| I-IU x Time | 1, 145 | 0.04 | .850 |

*Note*. Entries in the table that are formatted in bold indicate $p < .05$ and that the effect was significant when controlling for STICSA (or, where relevant, the alternate IU subscale). Black font indicates $p < .05$ and that the effect was not significant when controlling for STICSA (or, where relevant, the alternate IU subscale). Gray font denotes $p > .05$.

### Appendix I: Reliability, internal consistency of intolerance of uncertainty measures and their associations with trait anxiety

#### Reliability and internal consistency

To confirm our measures of intolerance of uncertainty were stable over time, we calculated measures of reliability and internal consistency to confirm stability of our measures, justifying the use of mean scores in our analyses.

The intraclass correlation coefficients (ICC) of intolerance of uncertainty and its prospective and inhibitory subscales were computed to assess their test-retest reliability across A: Time 1, 2 and 3, and B: Time 1 and 2 only. ICC estimates and their 95% CIs were computed using the *psych* package (Revelle, 2024) for each iteration based on a mean-rating (k = 3 for Times 1, 2 and 3; k = 2 for Times 1 and 2), absolute-agreement, two-way mixed-effects model.

The full ICCs are reported in **Supplementary Table *16*** below (and are identical to the table presented in the main manuscript). The ICCs for Time 1, 2 and 3 ranged from 0.90 to 0.92, and for Time 1 and 2 from 0.90 to 0.92, indicating excellent reliability according to the thresholds suggested by Koo & Li (2016). The corresponding F-tests indicated that each ICC was statistically significant (all *p*s < .001).

Given the excellent reliability of intolerance of uncertainty and its subscales across time, we concluded that there had not been major changes in IU across the three measured timepoints and expected the same to be true of trait anxiety, assessed here using STICSA only at Time 3. Scores for intolerance of uncertainty, and its prospective and inhibitory subscales were therefore averaged across Time 1 and Time 2 for participants who had only completed two timepoints (N = 27) and across Time 1, Time 2 and Time 3 who had completed all timepoints (N = 118). This was done to maintain a higher and better-powered sample size, with Time as a fixed effect, to enable clearer interpretations to be made in relation to the behaviour and computation modelling variables, which were only measured at Time 1 and Time 2.

**Supplementary Table 16**

*Results of Intraclass Correlation Coefficient Analyses for Intolerance of Uncertainty (IUS) and its Prospective (P-IU) and Inhibitory (I-IU) Subscales.*

| | | 95% CI | | | | |
|---|---|---|---|---|---|---|
| | **kappa** | **Lower** | **Upper** | ***F*** | **df** | ***p*** |
| **IUS** | | | | | | |
| Time 1, 2 and 3 | 0.92 | 0.89 | 0.94 | 12.53 | 117, 234 | < .001 |
| Time 1 and 2 | 0.92 | 0.89 | 0.94 | 13.07 | 144, 144 | < .001 |
| **P-IU** | | | | | | |
| Time 1, 2 and 3 | 0.90 | 0.86 | 0.93 | 10.01 | 117, 234 | < .001 |
| Time 1 and 2 | 0.90 | 0.86 | 0.93 | 9.61 | 144, 144 | < .001 |

| | kappa | 95% CI | | *F* | df | *p* |
|---|---|---|---|---|---|---|
| | | Lower | Upper | | | |
| **I-IU** | | | | | | |
| Time 1, 2 and 3 | 0.90 | 0.87 | 0.93 | 10.52 | 117, 234 | < .001 |
| Time 1 and 2 | 0.90 | 0.86 | 0.93 | 9.82 | 144, 144 | < .001 |

Correlations between intolerance of uncertainty/its subscales at each time were computed using the *psych* package. Each of the correlations demonstrated a significant positive association (all *ps* < .001), suggesting that intolerance of uncertainty and its subscale scores were strongly correlated across time. Full results can be found in **Supplementary Table *17***.

##### Supplementary Table 17

*Correlations Between Intolerance of Uncertainty Variables (IUS/P-IU/I-IU) at Each Time.*

| | *n* | *r* | *p* | 95% CI | |
|---|---|---|---|---|---|
| | | | | Lower | Upper |
| **IUS** | | | | | |
| Time 1 & Time 2 | 145 | 0.86 | < .001 | 0.81 | 0.90 |
| Time 1 & Time 3 | 118 | 0.76 | < .001 | 0.67 | 0.83 |
| Time 2 & Time 3 | 118 | 0.76 | < .001 | 0.68 | 0.83 |
| **P-IU** | | | | | |
| Time 1 & Time 2 | 145 | 0.81 | < .001 | 0.75 | 0.86 |
| Time 2 & Time 3 | 118 | 0.73 | < .001 | 0.63 | 0.80 |
| Time 2 & Time 3 | 118 | 0.73 | < .001 | 0.63 | 0.80 |
| **I-IU** | | | | | |
| Time 1 & Time 2 | 145 | 0.82 | < .001 | 0.75 | 0.86 |
| Time 1 & Time 3 | 118 | 0.73 | < .001 | 0.63 | 0.80 |
| Time 2 & Time 3 | 118 | 0.73 | < .001 | 0.64 | 0.81 |

Internal consistency of the intolerance of uncertainty scale, as well as its subscales, was computed separately at Time 1, 2 and 3 using the *cocron* package (Diedenhofen, 2016). Internal consistency for STICSA was computed for Time 3 only, as this was the only timepoint at which trait anxiety was measured.

All internal consistencies are reported in **Supplementary Table *18***. Intolerance of uncertainty plus its prospective and inhibitory subscales, as well as STICSA, demonstrated very good internal consistency, with Cronbach's a ranging from 0.87 to 0.92 for IUS and its subscales, and Cronbach's a of 0.91 for STICSA.

##### Supplementary Table 18

*Internal Consistency for Intolerance of Uncertainty (IUS), its Prospective (P-IU) and Inhibitory (I-IU) Subscales and STICSA.*

| | Cronbach's a | 95% CI | |
|---|---|---|---|
| | | Lower | Upper |
| **IUS** | | | |
| Time 1 | 0.90 | 0.88 | 0.92 |
| Time 2 | 0.91 | 0.89 | 0.93 |
| Time 3 | 0.92 | 0.90 | 0.94 |
| **P-IU** | | | |
| Time 1 | 0.87 | 0.84 | 0.90 |
| Time 2 | 0.87 | 0.84 | 0.90 |
| Time 3 | 0.87 | 0.84 | 0.90 |
| **I-IU** | | | |
| Time 1 | 0.88 | 0.86 | 0.91 |
| Time 2 | 0.88 | 0.86 | 0.91 |
| Time 3 | 0.87 | 0.84 | 0.90 |
| **STICSA** | 0.91 | 0.89 | 0.93 |

#### Associations with trait anxiety

Correlations between intolerance of uncertainty/its subscales at each time (as well as averaged across Time) and STICSA were computed using the *psych* package. Each of the correlations demonstrated a significant positive association (all *ps* < .001). Full results can be found in **Supplementary Table *19*** and are visualised in **Supplementary Figure *11***, **Supplementary Figure *12*** and **Supplementary Figure *13***.

Overall, our measures of intolerance of uncertainty (including its subscales) were stable over time and highly reliable across all timepoints in our sample. Intolerance of uncertainty and its subscales were further strongly positively related across time, as well as with our measure of trait anxiety (STICSA).

**Supplementary Table 19**

*Correlations Between Intolerance of Uncertainty Variables (IUS/P-IU/I-IU) and STICSA at Each Time, as well as Averaged Across Time 1 and 2.*

| | *n* | *r* | *p* | 95% CI | |
|---|---|---|---|---|---|
| | | | | Lower | Upper |
| **IUS & STICSA** | | | | | |
| Time 1 & STICSA | 118 | 0.52 | < .001 | 0.38 | 0.64 |
| Time 2 & STICSA | 118 | 0.53 | < .001 | 0.38 | 0.65 |
| Time 3 & STICSA | 118 | 0.62 | < .001 | 0.49 | 0.72 |
| Average Across Time & STICSA | 118 | 0.60 | < .001 | 0.47 | 0.70 |
| **P-IU & STICSA** | | | | | |
| Time 1 & STICSA | 118 | 0.43 | < .001 | 0.27 | 0.56 |
| Time 2 & STICSA | 118 | 0.45 | < .001 | 0.30 | 0.59 |
| Time 3 & STICSA | 118 | 0.56 | < .001 | 0.43 | 0.68 |
| Average Across Time & STICSA | 118 | 0.53 | <.001 | 0.39 | 0.65 |
| **I-IU & STICSA** | | | | | |
| Time 1 & STICSA | 118 | 0.55 | < .001 | 0.41 | 0.66 |
| Time 2 & STICSA | 118 | 0.54 | < .001 | 0.39 | 0.65 |

|  | n | r | p | 95% CI |  |
|---|---|---|---|---|---|
|  |  |  |  | Lower | Upper |
| Time 3 & STICSA | 118 | 0.60 | < .001 | 0.47 | 0.71 |
| Average Across Time & STICSA | 118 | 0.61 | < .001 | 0.48 | 0.71 |

**Supplementary Figure 11**

*Scatterplots with Histograms Depicting Correlation Between IUS and STICSA at **(A)** Time 1, **(B)** Time 2, **(C)** Time 3, and **(D)** IUS Averaged Across Times 1 and 2. The distribution of IUS scores is displayed on top of the figures in pink, and the distribution of STICSA is displayed on the right side of the figures in orange. Shaded areas represent 95% of confidence intervals. Higher IUS at each Time was associated with higher levels of STICSA.*

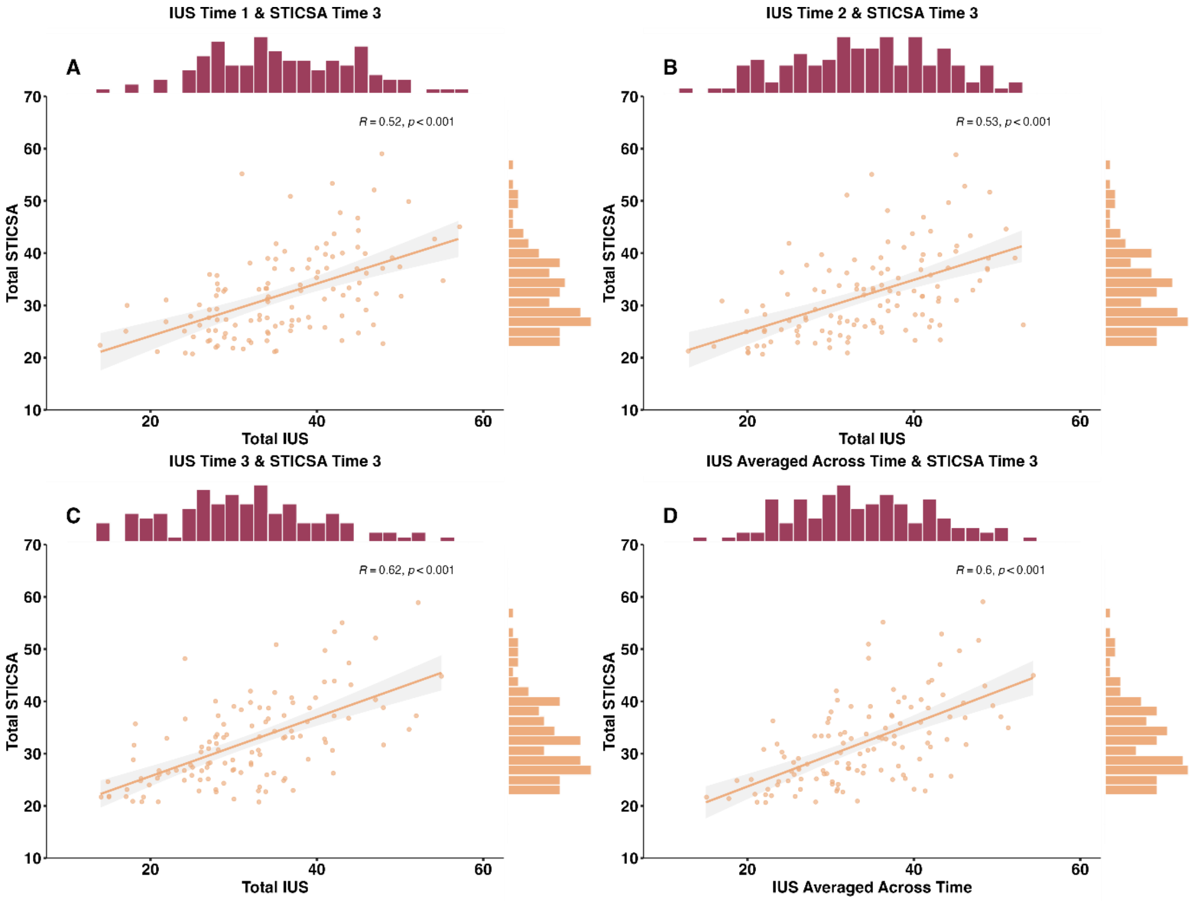


**Supplementary Figure 12**

*Scatterplots with Histograms Depicting Correlation Between Prospective Intolerance of Uncertainty (P-IU and STICSA at **(A)** Time 1, **(B)** Time 2, **(C)** Time 3, and **(D)** P-IU Averaged Across Times 1 and 2. The distribution of P-IU scores is displayed on top of the figures in pink, and the distribution of STICSA is displayed on the right side of the figures in orange. Shaded areas represent 95% of confidence intervals. Higher P-IU at each Time was associated with higher levels of STICSA.*

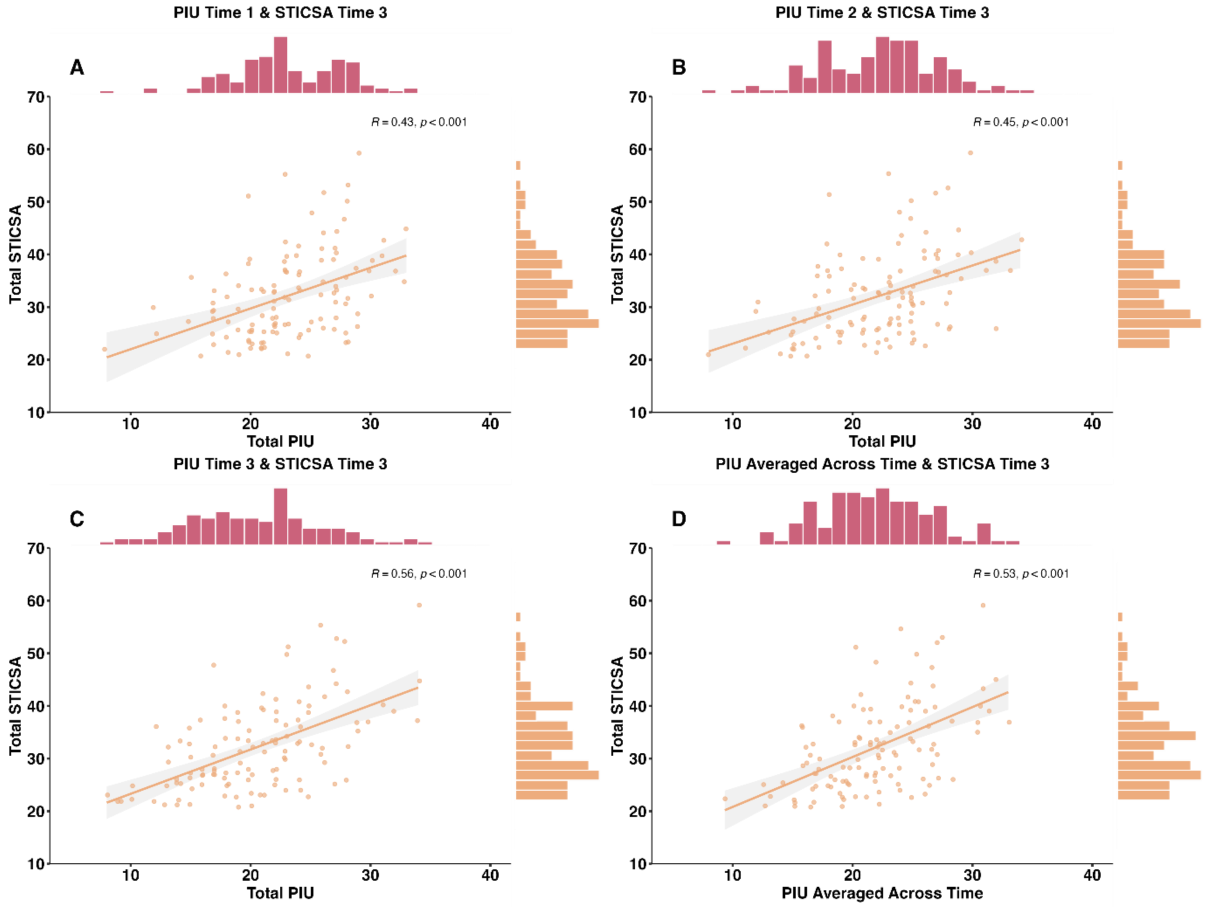


#### Supplementary Figure 13

*Scatterplots with Histograms Depicting Correlation Between Inhibitory Intolerance of Uncertainty (I-IU and STICSA at* ***(A)*** *Time 1,* ***(B)*** *Time 2,* ***(C)*** *Time 3, and* ***(D)*** *I-IU Averaged Across Times 1 and 2. The distribution of I-IU scores is displayed on top of the figures in pink, and the distribution of STICSA is displayed on the right side of the figures in orange. Shaded areas represent 95% of confidence intervals. Higher I-IU at each Time was associated with higher levels of STICSA.*

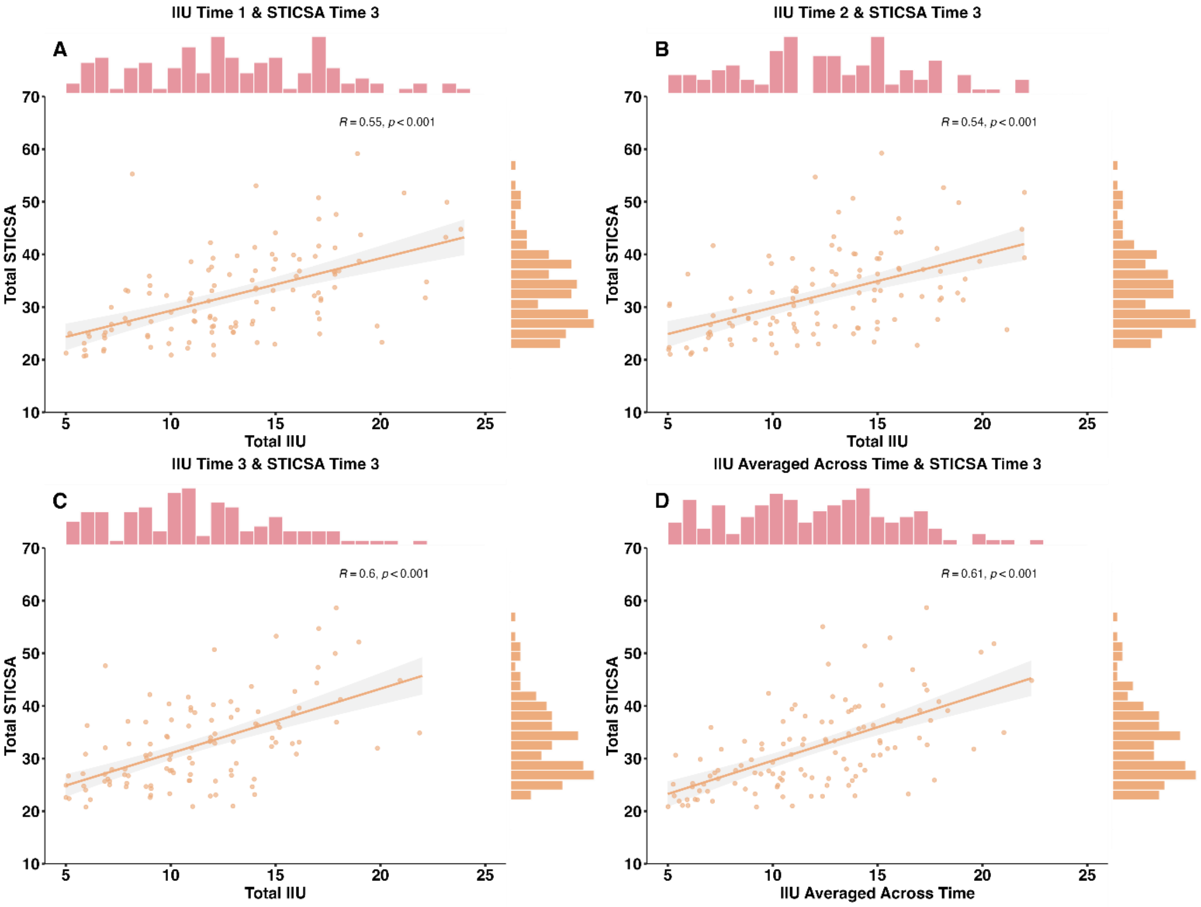

IIU Time 1 & STICSA Time 3
A
R = 0.55, p < 0.001
Total STICSA
Total IIU
IIU Time 2 & STICSA Time 3
B
R = 0.54, p < 0.001
Total STICSA
Total IIU
IIU Time 3 & STICSA Time 3
C
R = 0.6, p < 0.001
Total STICSA
Total IIU
IIU Averaged Across Time & STICSA Time 3
D
R = 0.61, p < 0.001
Total STICSA
IIU Averaged Across Time

### Appendix J: Summary of MLM Results

**Supplementary Table 20**

*Summary of MLM Results*

| | Reaction Time | Stay | Accuracy | Accuracy Slope | Perseveration | Difference in Reaction Times | Reinforcement Sensitivity | Learning Rate | Inverse Temperature | Learning Rate | Discount Weight |
|---|---|---|---|---|---|---|---|---|---|---|---|
| **Time x Outcome** | | | | | | | | | | | |
| Time | < .001 | .006 | < .001 | <. 001 | < .001 | < .001 | < .001 | .772 | < .001 | .623 | .004 |
| Outcome | .001 | < .001 | | | | | <. 001 | | .231 | < .001 | |
| Time x Outcome | .083 | .311 | | | | | < .001 | | .500 | .010 | |
| **IUS x Time (x Outcome)** | | | | | | | | | | | |
| IUS | .020 | .184 | .104 | .170 | .114 | .250 | .105 | .775 | .051 | .536 | .623 |
| IUS x Time | .057 | .691 | .737 | .590 | .675 | .203 | .406 | .080 | .052 | .229 | .452 |
| IUS x Outcome | .372 | .654 | | | | | **.001** | | .011 | .434 | |
| IUS x Time x Outcome | .690 | .577 | | | | | .258 | | .320 | .394 | |
| **P-IU x Time (x Outcome)** | | | | | | | | | | | |
| P-IU | .074 | .239 | .310 | .400 | .344 | .432 | .157 | .733 | .129 | .314 | .986 |
| P-IU x Time | .162 | .692 | .887 | .767 | .788 | .110 | .475 | .036 | **.021** | .170 | .234 |
| P-IU x Outcome | .543 | .608 | | | | | .021 | | .025 | .588 | |
| P-IU x Time x Outcome | .618 | .511 | | | | | .314 | | .188 | .476 | |
| **I-IU x Time (x Outcome)** | | | | | | | | | | | |
| I-IU | .008 | .177 | **.033** | .070 | **.035** | .153 | .095 | .344 | .027 | .902 | .336 |

| | Reaction Time | Stay | Accuracy | Accuracy Slope | Perseveration | Difference in Reaction Times | Reinforcement Sensitivity | Learning Rate | Inverse Temperature | Learning Rate | Discount Weight |
|---|---|---|---|---|---|---|---|---|---|---|---|
| I-IU x Time | .240 | .723 | .608 | .455 | .590 | .438 | .385 | .238 | .182 | .375 | .850 |
| I-IU x Outcome | .268 | .743 | | | | | **< .001** | | .009 | .337 | |
| I-IU x Time x Outcome | .807 | .697 | | | | | .254 | | .596 | .362 | |

### Appendix K: Summary of Post-Hoc Correlation Results

**Supplementary Table 21**

*Summary of Post-Hoc Correlation Results*

| | *r* | df | *p* |
|---|---|---|---|
| **Accuracy** | | | |
| **Main effect of I-IU** | | | |
| Partial correlation controlling for STICSA | **0.22** | **119** | **.018** |
| Partial correlation controlling for P-IU | **0.18** | **145** | **.031** |
| **Perseveration** | | | |
| **Main effect of I-IU** | | | |
| Partial correlation controlling for STICSA | **-0.20** | **119** | **.026** |
| Partial correlation controlling for P-IU | **-0.19** | **145** | **.026** |
| **Reinforcement Sensitivity** | | | |
| **IUS x Outcome Interaction** | | | |
| **Win** | | | |
| Partial correlation controlling for STICSA | **0.18** | **119** | **.049** |
| **Loss** | | | |
| Partial correlation controlling for STICSA | 0.02 | 119 | .838 |
| **I-IU x Outcome Interaction** | | | |
| **Win** | | | |
| Partial correlation controlling for STICSA | 0.17 | 119 | .072 |
| Partial correlation controlling for P-IU | 0.14 | 145 | .095 |
| **Loss** | | | |
| Partial correlation controlling for STICSA | -0.02 | 119 | .840 |
| Partial correlation controlling for P-IU | -0.17 | 145 | .037 |
| **Inverse Temperature** | | | |
| **P-IU x Time Interaction** | | | |
| **Time 1** | | | |
| Partial correlation controlling for STICSA | 0.14 | 119 | .125 |
| Partial correlation controlling for I-IU | -0.11 | 145 | .202 |
| **Time 2** | | | |
| Partial correlation controlling for STICSA | 0.14 | 119 | .125 |
| Partial correlation controlling for I-IU | 0.04 | 145 | .656 |

### Appendix L: Results from STICSA MLMs

We conducted additional MLMs to test the effects of trait anxiety (STICSA) on the dependent variables of interest. The MLMs were coded as described in the main manuscript, only instead with mean-centred STICSA scores entered as the main individual differences independent variable. In cases of a significant main effect or interaction, we conducted similar post-hoc MLMs to test whether the STICSA effects would hold after controlling for IUS.

##### Supplementary Table 22

*Results from MLMs Assessing STICSA Main Effects and Interactions with Time (and Outcome) on Behavioural and Computational Data*

| | df | *F* | *p* |
|---|---|---|---|
| **Reaction Times** | | | |
| STICSA | 1, 119 | 3.12 | .080 |
| STICSA x Time | **1, 357** | **22.29** | **< .001**[1] |
| STICSA x Outcome | 1, 357 | 0.74 | 0.393 |
| STICSA x Time x Outcome | 1, 357 | 0.04 | .841 |
| **Stay** | | | |
| STICSA | 1, 119 | 0.02 | .893 |
| STICSA x Time | 1, 357 | 0.06 | .803 |
| STICSA x Outcome | 1, 357 | 1.17 | .280 |
| STICSA x Time x Outcome | | | |
| **Accuracy** | | | |
| STICSA | 1, 119 | 0.80 | .374 |
| STICSA x Time | 1, 119 | 0.10 | .756 |
| **Accuracy Slope** | | | |
| STICSA | 1, 119 | 0.27 | .603 |
| STICSA x Time | 1, 119 | 0.10 | .758 |
| **Perseveration** | | | |
| STICSA | 1, 119 | 0.72 | .396 |
| STICSA x Time | 1, 119 | 0.31 | .578 |
| **Difference in Reaction Times** | | | |
| STICSA | 1, 119 | 1.15 | .287 |
| STICSA x Time | 1, 119 | 0.39 | .531 |
| **Reinforcement Sensitivity** | | | |
| STICSA | 1, 119 | 0.18 | .676 |
| STICSA x Time | 1, 357 | 0.02 | .878 |
| STICSA x Outcome | 1, 357 | 6.47 | .011[2] |
| STICSA x Time x Outcome | 1, 357 | 0.11 | .738 |

[1] The STICSA x Time interaction on RTs remained significant after controlling for IUS [STICSA x Time: $F(1, 357) = 16.99$, $p < .001$; IUS x Outcome: $F(1, 357) = 0.32$, $p = .569$].

[2] The STICSA x Outcome interaction on reinforcement sensitivity did not hold after controlling for IUS [STICSA x Outcome: $F(1, 357) = 0.28$, $p = .599$; IUS x Outcome: $F(1, 357) = 6.53$, $p = .011$].

| | df | *F* | *p* |
|---|---|---|---|
| **Learning Rate** | | | |
| STICSA | 1, 119 | 0.08 | .777 |
| STICSA x Time | 1, 119 | 0.17 | .679 |
| **Inverse Temperature** | | | |
| STICSA | 1, 119 | 2.48 | .118 |
| STICSA x Time | 1, 357 | 1.36 | .245 |
| STICSA x Outcome | 1, 357 | 5.94 | .015[3] |
| STICSA x Time x Outcome | 1, 357 | 0.68 | .411 |
| **Learning Rate** | | | |
| STICSA | 1, 119 | 0.63 | .430 |
| STICSA x Time | 1, 357 | 0.11 | .745 |
| STICSA x Outcome | 1, 357 | 0.38 | .537 |
| STICSA x Time x Outcome | 1, 357 | 0.86 | .353 |
| **Discount Weight** | | | |
| STICSA | 1, 119 | 0.66 | .417 |
| STICSA x Time | 1, 119 | 0.18 | .674 |

*Note*. Entries in the table that are formatted in bold indicate $p < .05$ and that the effect was significant when controlling for IUS. Black font indicates $p < .05$ and that the effect was not significant when controlling for IUS. Gray font denotes $p > .05$.

[3] The STICSA x Outcome interaction on inverse temperature did not hold after controlling for IUS [STICSA x Outcome: $F(1, 357) = 1.19$, $p = .276$; IUS x Outcome: $F(1, 357) = 2.16$, $p = .143$].
